# Neonatal social communication and single genes predict the variability of post-pubertal social behavior in a mouse model of paternal 15q11-13 duplication

**DOI:** 10.64898/2026.01.17.700083

**Authors:** Takahira Yamauchi, Kota Tamada, Takeshi Takano, Mitsuteru Nakamura, Mariel Barbachan E. Silva, Kenny Ye, Hitoshi Inada, Takaki Tanifuji, Takeshi Hiramoto, Lucas Stevens, Gina Kang, Marisa Esparza, Takefumi Kikusui, Noriko Osumi, Pilib Ó Broin, Toru Takumi, Noboru Hiroi

## Abstract

Mental illnesses associated with high-risk copy number variations (CNVs) are characterized by incomplete penetrance and variable severity, with their underlying mechanisms remaining inadequately understood. We hypothesized that such phenotypic variability is evident from the neonatal stage and is, at least in part, attributable to individual differences in the expression levels of CNV-encoded genes in the brain. We conducted an analysis of the quantitative and functional structure of neonatal social communication, assessed post-pubertal social interaction, and evaluated the brain expression levels of genes within the same cohort of a mouse model of paternal human 15q11-13 duplication, a high-risk factor variably associated with neurodevelopmental disorders.

Subsequently, computational methods were utilized to identify predictive variables for the variability of post-pubertal social interaction. Mice harboring the 15q11-13 duplication exhibited distinctive call sequences characterized by diverse connections, which lacked the incentive value necessary for effective social communication with mother mice. The neonatal call sequences and the expression levels of *Magel2*, along with, to a lesser extent, *Herc2* and *Ndn*, in the prefrontal cortex of the 15q11-13 duplication model were predictive of post-pubertal social interaction. Our findings demonstrate that variability in post-pubertal social interaction—a dimensional characteristic of neurodevelopmental disorders—can be predicted by the variability of neonatal social communication and is influenced by the expression levels of specific CNV-encoded genes in the prefrontal cortex. This computational approach has the potential to predict the developmental trajectories of various dimensions of mental illness among CNV carriers in humans and to identify CNV-encoded driver genes in preclinical models, thereby providing potential mechanistic bases for the development of gene-based therapeutic strategies.

## INTRODUCTION

Copy number variations (CNVs) represent chromosomal deletions and duplications of several million base pairs within and are associated with significantly elevated risks of developing mental illnesses, including schizophrenia, autism spectrum disorder (ASD), intellectual disability, bipolar disorder, mood disorders, attention-deficit/hyperactivity disorder, and various other psychiatric disorders [1]. These genetic variations serve as a reliable entry point for investigating the mechanistic underpinnings of psychiatric disorders[2–4]. However, a significant challenge in formulating therapeutic strategies based on CNV is that each CNV encompasses multiple genes, and it remains unclear whether all encoded genes and their molecular pathways are functionally relevant to a specific phenotype and how each gene contributes to a wide range of phenotypes [3, 5, 6].

One potential strategy to address this challenge is to identify variants among CNV-encoded genes in individuals without CNVs. Recent large-scale analyses have detected ultra-rare variants of CNV-encoded single genes in idiopathic cases of schizophrenia and ASD [7–12]. Although these studies have provided valuable insights, considerable interpretative challenges persist. Firstly, the limited number of identified variants for each gene does not allow for statistically robust associations between variants and psychiatric disorders. Secondly, the absence of ultra-rare variants of a CNV-encoded gene does not constitute evidence of its lack of contribution to psychiatric disorders. Indeed, studies with larger sample sizes tend to uncover more ultra-rare single-gene variants of CNVs compared to those with smaller sample sizes. Due to the technical challenges of identifying CNV-encoded driver genes in humans, it is often postulated, without empirical validation, that all genes within each CNV functionally contribute to mental illness or specific phenotypes.

Phenotypic variability is another aspect of human CNVs. Not all individuals harboring CNVs are diagnosed with a psychiatric disorder (i.e., incomplete penetrance), and the same CNVs may contribute to diverse disorders in different individuals (i.e., pleiotropy). Furthermore, the severity of each disorder varies among individual CNV carriers (i.e., variable expressivity) [3, 5]. Environmental, stochastic, prenatal, and postnatal factors play a role in these variabilities. While individual variation in CNV-encoded gene expression in the brain is another potential contributor, quantifying CNV-encoded gene expression levels and their variability in the human brain during and around symptomatic phases is not feasible.

The developmental origins of phenotypic variability constitute another fundamental question that remains unresolved. While the presence of severe core symptoms is a prerequisite for clinical diagnosis, various social, cognitive, and motor measures deviate from normative standards among infants who are later diagnosed with psychiatric disorders [13–15]. Atypical preverbal vocalizations are one such measure [16–18] and has been proposed as a prerequisite for the development of subsequent social behavior impairments [16, 19]. However, establishing a prospective relationship between neonatal indicators and later phenotypes in the same individuals requires years, or even decades, of observation in humans.

Duplications at human chromosome 15q11-13 raise the risk of developing intellectual disability, ASD, and schizophrenia[1]. Maternally-derived 15q11-13 duplication s are more frequently observed in individuals with various neurodevelopmental phenotypes, compared to paternally-derived cases [20–32]. However, paternally-derived duplications and triplications are variably linked to an elevated risk for ASD, intellectual disability, developmental delays, epilepsy/seizures, behavioral problems, or a combination of these diagnoses.[22, 26–28, 32–40] However, the source of phenotypic variability remains unclear. Ultra-rare protein-truncating variants of several 15q11-13 genes, such as *GABRA5* and *GABRB3* in schizophrenia[10], *MAGEL2 in ASD*[8], and *GABRA5 and ATP10A* in bipolar disorder[41], have been identified with high odds ratios. One patient diagnosed with ASD has been reported to carry a small paternally inherited duplication limited to a small region including *TUBGCP5, CYFIP1, NIPA2, NIPA1, MKRN3, MAGEL2*, and *NDN* [27]. While these associations with single genes are suggestive, they have not consistently reached statistical significance due to their rarity, and the contributions of each encoded gene to social dimensions remain unclear.

The technical limitations inherent in human studies pose challenges to mechanism-based predictions and therapeutic advancements. A genetic mouse model provides a complementary approach to address these methodological challenges[3–5, 42]. Although mouse models do not fully replicate the entire symptomatology of each clinically defined psychiatric disorder, they offer a means to model certain aspects of mental illness [2, 3, 5]. Deficits in social dimensions are observed in both idiopathic and CNV-linked schizophrenia and ASD[15, 43–47]. CNV-encoded single genes in mouse models are implicated in cognitive and social dimensions[3–5, 48–50] that are adversely affected in humans with CNVs [51–55] and in idiopathic schizophrenia and ASD [3].

To evaluate the hypothesis that the variability of post-pubertal social behavior has a neonatal origin and is influenced by expression levels of CNV-encoded genes, we utilized a mouse model of paternally-inherited 15q11-13 duplication [56]. We sequentially assessed neonatal vocalizations, post-pubertal social interactions, and gene expression in the post-pubertal brains of the same cohort in the paternal duplication model of 15q11-13. A machine learning algorithm was employed to identify predictive variables for post-pubertal social interaction based on a pooled set of neonatal social communication variables and post-pubertal gene expression data. Our results identified specific predictive neonatal and gene variables associated with post-pubertal social behavior. This computational approach to a preclinical model offers a powerful and complementary means to enhance our understanding of the developmental and genetic origins of phenotypic variability linked to dimensions of CNV-associated mental illness.

## METHODS

### Ethics Approval

The use of animal subjects complied with protocols approved by the Animal Care and Use Committee of the Albert Einstein College of Medicine (Animal Welfare Assurance A3312-01), the University of Texas Health Science Center at San Antonio (Animal Welfare Assurance 3345-01, 20190084AR), the RIKEN Center for Brain Science (2018-056), and Kobe University School of Medicine (P200104-R12), in accordance with guidelines established by the National Institutes of Health.

### Mice

We utilized a congenic mouse model with a duplication of the 15q11-13 region; the original mouse lineage [56] was backcrossed to C57BL/6J mice for over 10 generations to reduce the unequal genetic backgrounds between wild-type and mutant littermates[4]. Each breeding pair consisted of one male paternal 15q11-13 duplication (Dup/+) model mouse (aged 10–20 weeks) [56] and one female C57BL/6J mouse (aged 10 to 30 weeks) sourced from Japan SLC, Inc., Shizuoka, Japan. Offspring were housed in groups of two to four male wild-type and Dup/+ littermates per cage. The minimum sample size was determined through power analyses based on prior studies. All male mice in each litter were used. The litter sizes were indistinguishable between +/+ and Dup/+ mice (+/+, Average=4.6, SEM=0.327; Dup/+, Average=4.7, SEM=0.334; Mann-Whitney U test, U=350.5, p=0.850).

Male mice were randomly assigned to experimental groups, and experimenters were blinded to genotypes during group allocation and testing (see **Supplementary Table S2** for sample sizes).

We did not use female mice for three primary reasons. First, initiating testing of post-pubertal social interactions at the same estrous stage across all female mice at 10 weeks of age was technically difficult. Second, conducting social interaction assessments and subsequent sacrifices at the same estrous stage posed technical challenges. Third, brain gene expression fluctuates throughout the estrous cycle in mice[57].

Genotyping was performed using forward (5′-ATATGTACTTTTGCATATAGTATAC-3′) and reverse (5′-AGAGGAGGGCCTTACTAATTACTTA-3′) primers.

### Behavioral Assays

Ultrasonic vocalization was assessed during maternal separation at postnatal days (P) P8 and P12, and post-pubertal social behavior was evaluated at 10 weeks of age. Neonatal vocalizations were analyzed both qualitatively and quantitatively, while post-pubertal social behavior was recorded and manually rated following established methodologies [50, 58–64].

To evaluate the incentive value of neonatal vocalizations, we measured the approach behavior of C57BL/6J mothers that had not previously been exposed to the vocalizations of either +/+ or Dup/+ pups from the 15q11-13 duplication model, following our procedures [61](see **Supplementary Materials**).

### Quantitative Reverse Transcription Polymerase Chain Reaction

The mice were euthanized after the completion of behavioral testing, . The prefrontal cortex (the entire cortex anterior to Bregma +1.98) and the remaining forebrain (excluding the olfactory bulb) were dissected, frozen, and utilized for quantitative reverse transcription polymerase chain reaction (qRT-PCR) analyses (see **Supplementary Materials** and **Supplementary Table S1**). The cerebellum and brainstem were excluded from this analysis.

### Computational analyses

We conducted Shannon entropy analysis, the Markov model, UMAP (uniform manifold approximation and projection), and the Lasso (least absolute shrinkage and selection operator) regression model in accordance with our previously published methodologies [59, 61, 65, 66] (see **Methods** and **Supplementary Materials**).

### Statistical Analysis

We used SPSS (v29.0.2.0 (20), IBM Corporation). Group means were compared using analysis of variance. In instances where interaction effects were statistically significant, Student’s T-tests were employed to elucidate the nature of the interaction; two-sided t-tests were applied for comparisons between two groups. A p-value of less than 0.05 was considered statistically significant. The significance level for multiple tests was adjusted using the Benjamini–Hochberg correction at false discovery rates (FDR) of 5%, 10% and 25%. Data that violated the assumptions of homogeneity of variance or normality were analyzed using linear mixed effect models with the individual animal as the random effect or nonparametric tests. All statistical values are detailed in **Supplementary Table S1**.

## Results

### Characterization of Wave Shapes and Patterns of Neonatal Vocalizations

Our previous work indicated that paternal Dup/+ pups exhibited a significantly higher frequency of ultrasonic vocalizations compared to their +/+ littermates on postnatal days 7 (P7) and 14 (P14). However, no significant differences were observed between the two groups before or after this time frame [56]. Consequently, in the current study, we selected postnatal days 8 (P8) and 12 (P12) as the periods of interest for evaluating ultrasonic vocalizations.

Consistent with this previous observation of ours, the total number of ultrasonic vocalizations at P12 is greater in Dup/+ pups than in +/+ littermates, although no statistically significant differences are noted between the two genotypes at P8 (**Figure 1A, inset**).

**Figure 1.**
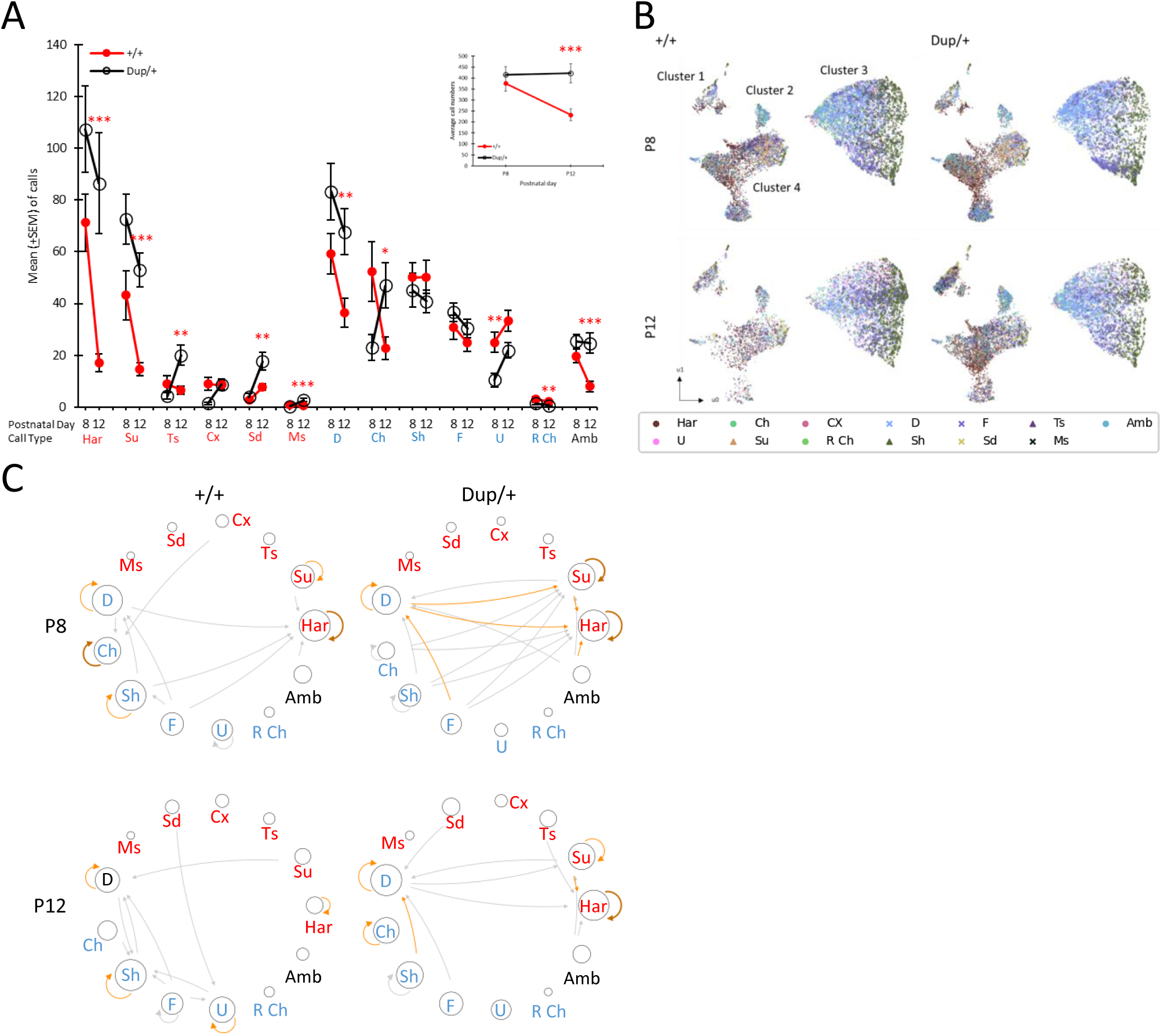
**A**. The number of 13 call types (mean±SEM) emitted at postnatal day (P) P8 and P12 for Dup/+ (white circle with black line) and +/+ (red circle with red line) mice. As the assumptions of normality and homogeneity of variance were violated in many cases (see **Table S2**-Figure 1), we used Mann-Whitney U tests. The Benjamini–Hochberg method adjusted the significance at the false detection rate of 5%. The original p values of cases that remained statistically significant after this adjustment are indicated as *, p<0.05; **, p<0.01; ***, p<0.001. **A. Inset**: The number of total ultrasonic vocalizations (mean±SEM) emitted at P8 and P12 for Dup/+ and +/+ mice. The two genotypes differed, but this effect depended on age (genotype, F(1,51)=8.354, p=0.006; age, F(1,51)=4.626, p=0.036; genotype × age, F(1,51)=5.463, p=0.023). ***, p<0.001, as determined by t-tests. Har, harmonic; Su, step-up; Ts, two-steps; Cx, complex; Sd, step-down; Ms, multiple-steps; D, downward; Ch, chevron; Sh, short; F, flat; U, upward; R Ch, reverse chevron; Amb, ambiguous. Red call type names are those of more than one wave or complex waves; blue call type names are those of relatively simple waves[59, 61]. P8: +/+, n=29; Dup/+, n=25. P12: +/+, n=28; Dup/+, n=25. One +/+ pup emitted no call at P12 and was thus omitted from analysis. **B**. UMAP (uniform manifold approximation and projection) of the quantitative parameters of calls. The calls separated into four major clusters. Each dot represents a call emitted by a pup. Call types are added with distinct colors to indicate the locations of each call type in clusters. P, postnatal day. **C**. Markov models for the +/+ and Dup/+ mice at P8 and P12. The size of each call circle represents the non-Markov proportion of each call type (see **Figures S2**). Lines and their thicknesses represent the connections among call types and their significance, respectively (see **Table S2_**Figure 1C). Dark brown, p>0.1923; light brown, p>0.1539; grey, p>0.1154. Har, harmonic; Su, step-up; Ts, two-steps; Cx, complex; Sd, step-down; Ms, multiple-steps; D, downward; Ch, chevron; Sh, short; F, flat; U, upward; R Ch, reverse chevron; Amb, ambiguous.

Neonatal mouse vocalizations are characterized by a diverse array of wave shapes and patterns [67], with these parameters being differentially influenced by specific genes or sets of genes associated with neurodevelopmental disorders [59, 61, 63, 65, 68, 69]. To further investigate the impact of the Dup/+ genotype on various call types, we conducted an analysis utilizing our previously published methodology for call classification [65] (**Figure S1; Supplementary Materials,** Call-type classification). At P12, Dup/+ pups emitted a greater number of harmonic, step-up, two-step, step-down, multiple-step, downward, chevron, and ambiguous calls compared to +/+ littermates.

Conversely, Dup/+ pups exhibited lower frequencies of upward calls at P8 and reverse chevron calls at P12 in comparison to +/+ pups (**Figure 1A**).

To determine whether the observed differences in call counts could be attributed to an overall increase in vocalizations by Dup/+ pups, we analyzed the proportions of each call type within each genotype.

Dup/+ pups displayed significantly higher proportions of harmonic, step-up, two-step, multiple-step, and ambiguous call types relative to +/+ littermates. In contrast, Dup/+ pups proportionally emitted fewer short, upward, and reverse chevron calls than their +/+ counterparts (**Figure S2**).

In summary, the Dup/+ genotype significantly influences the emission of harmonic, step-up, two-step, multiple-step, short, upward, reverse chevron, and ambiguous calls beyond what would be expected based solely on their absolute numbers. The genotype-dependent variations in the number of step-down, downward, and chevron calls can be attributed to their higher frequencies observed in Dup/+ pups. Although the counts of the short call type did not show significant differences between the two genotypes, Dup/+ pups exhibited a proportionally lower emission of these calls compared to +/+ pups.

### Characterization of Acoustic Features of Neonatal Vocalizations

In addition to categorizing neonatal calls by wave shapes and patterns, each call exhibits a distinct set of quantitative acoustic properties. We employed the Uniform Manifold Approximation and Projection (UMAP) method [70] to elucidate the dimensionality of the quantitative features of calls. UMAP is a dimensionality reduction technique based on Riemannian geometry and algebraic topology, which effectively clusters data exhibiting similar quantitative characteristics.

The quantitative acoustic parameters of the calls, analyzed using VocalMat [71], include: a) bandwidth measured in Hz; b) maximum, mean, and minimum frequencies in Hz of a call or its principal components; and c) maximum, mean, and minimum intensities of each call. The UMAP analysis resulted in the formation of four principal spatial clusters based on these quantitative properties.

Subsequently, we incorporated the categorical classification of calls into the UMAP framework (**Figure 1B**).

Various call types were represented within or across these clusters (**Figure 1B**). Each cluster displayed a unique set of call-type representations (**Figure S3-1** to **S3-4**), but the shapes and positions of the four clusters did not seem to differ between the Dup/+ and +/+ genotypes, suggesting that the Dup/+ genotype did not substantially influence the overall quantitative features of the calls.

Conversely, the density of certain calls exhibited genotype-dependent variations. The genotype-dependent differences in call numbers, indicated by the dots in **Figure 1B**, were more pronounced in the data from postnatal day 12 (P12). Specifically, Dup/+ pups exhibited a greater number of calls in Clusters 1, 2, and 4 compared to +/+ pups at P12, whereas an opposite trend was observed in Cluster 3. These patterns align with the categorization analysis of call types (see **Figure 1A; S3-1 to S3-4**).

Notably, the increased density of calls observed in Cluster 1 of P12 Dup/+ mice, compared to P8 Dup/+ (**Figure 1B**), is attributed to higher levels of harmonic and step-up call types (see **Figure S3-1**). The increased calls in Cluster 1 from P8 to P12 in Dup/+ pups reflect more step-up, two-step, complex, multiple-step, and downward call types emitted (see **Figure S3**).

### Determination of Call Sequences

The vocal call sequences of pups have significant biological implications. When presented with the complete call sequence of a pup, mothers exhibit a strong maternal response; however, if the sequence is randomized while retaining all call types and amplitudes, its ability to elicit a maternal approach is diminished[61]. Furthermore, the call sequences of pups with a gene-dose alteration associated with neurodevelopmental disorders (e.g., *Tbx1*) lack incentive value for maternal engagement[61].

Calls are emitted with varying inter-call intervals. While it remains unclear whether a functional unit exists within a series of mouse pup calls, we operationally defined a call sequence as comprising inter-call intervals shorter than those theoretically expected from a given number of calls emitted within a specified testing period (i.e., Poisson distribution)[59, 61, 65]. This methodology was applied to the calls of Dup/+ and +/+ littermates (**Figure S4**). The distribution curves of observed inter-call intervals intersected with those of the theoretical inter-call intervals at 362.41 ms and 408.66 ms for +/+ and Dup/+ pups, respectively, at P8; the intersection points were 366.55 ms and 314.99 ms for +/+ and Dup/+ pups, respectively, at P12 . When an inter-call interval exceeded the intersection point values, calls preceding and succeeding the interval were classified as the last and first calls of two consecutive, distinct sequences, respectively; calls with inter-call intervals shorter than the intersection point values were considered components of a sequence.

### Degree of Unpredictability of Call Selection and Sequencing

Following the definition of the call sequence, we examined whether calls are emitted unpredictably as single instances or in sequences. Shannon entropy scores were utilized to assess the degree of unpredictability in the selection of call types (H0), the distribution of distinct calls within the selected call types (H1), and the distributions of distinct call types in two-call sequences (H2), three-call sequences (H3), and four-call sequences (H4).

The degree of unpredictability decreased at each H level, indicating the presence of predictable elements as pups select call types and establish connections within sequences. Entropy scores consistently declined at each H level for both Dup/+ and +/+ pups at P8 and P12. The choice of call types and their connections had equal levels of unpredictability between +/+ and Dup/+ pups at each level at both P8 and P12 (**Figure S5AB**), except for sequences at H3 and H4 at P8 (**Figure S5A**).

These findings suggest that Dup/+ pups and +/+ pups generated similarly predictable call sequences, despite the observation that Dup/+ pups emitted a greater number of calls than their +/+ counterparts at P12. At P8, Dup/+ pups emitted three- and four-call sequences more unpredictably than +/+ pups.

Dup/+ and +/+ pups did not differ in the number of calls per sequence (**Figure S6A**). By contrast, Dup/+ pups emitted longer sequences than +/+ pups at P8, but not at P12 (**Figure S6B**). Dup/+ pups emitted more sequences, compared to +/+ pups, at P12, but not at P8 (**Figure S6C**).

### Proportions of Two-Call Sequences

The entropy analysis does not consider the variability of call types among individual pups. For example, if call type A is followed by call type B in one pup while call type C is followed by call type D in another, both scenarios yield an identical entropy score. Consequently, we additionally examined the specific call types in every pair of consecutive calls within the call sequences. We analyzed the proportions of each two-call connection (see **Figure S7**). This analysis revealed that both Dup/+ and +/+ pups emit a greater proportion of specific two-call connections (e.g., harmonic followed by harmonic at P8) and exhibit differentially expressed connections between the genotypes (e.g., at P8, chevron followed by chevron; downward followed by step-up; at P12, harmonic followed by harmonic; short followed by flat). At P12, the most proportionally frequent call connections were short followed by short in +/+ and harmonic followed by harmonic in Dup/+ pups. +/+ pups tended to make connections among simple call types (i.e., downward, short, flat, and upward; blue letter calls), while Dup/+ pups connected more diverse call types, including both simple waves and multiple waves (see more scattered call connections). These findings also indicate that the proportions of various two-call connections within sequences differ between the two genotypes. Indeed, the trajectories of call sequences appear to diverge in the three-dimensional UMAP space (see **Figure S8**). More diverse call transitions between cluster 3 and the other clusters appear in Dup/+ pups and transitions tend to occur more frequently within cluster 3 in +/+ pups, where simple call types aggregate (see **Figures S3** and **S7**).

### Finite State Rule in Two-Call Sequences

Pups display distinct sequences of calls, and temporal call transitions appear to differ between the genotypes (see **Figure S7, S8**). However, in this qualitative analysis, sequences that begin with less frequently emitted call types are underrepresented, while those starting with more frequently emitted call types are overrepresented. For instance, even if a particular call consistently precedes another call type, such a connection may not be well represented in terms of proportions if the first call’s frequency is low relative to the total call count. The Markov property addresses this technical limitation by positing that the future state depends solely on the current state. Consequently, the probability of the second call type in each two-call connection is calculated exclusively based on the first call type. In this framework, the probabilities of transitions from one call to the next are unaffected by the absolute number or proportion of the first call among all emitted calls. Our previous work indicated that mouse pups with *Tbx1* heterozygosity, 16p11.2 hemizygosity, and *Fmr1* deletion—risk gene variants linked to increased susceptibility to neurodevelopmental disorders—exhibit alterations in the finite state rule governing two-call connections within sequences[59, 61, 65].

At P8, both Dup/+ and +/+ pups utilized harmonic, step-up, downward, chevron, and short call types in repeats (e.g., harmonic followed by harmonic). Compared to +/+ pups, Dup/+ pups showed more diverse call connections between different call types (**Figure 1C, P8**).

At P12, +/+ pups exhibited self-repeat connections within harmonic, downward, short, flat, and upward calls, while Dup/+ pups did so within step-up, harmonic, downward, chevron, and short call types (see **Figure 1C, P12**). Both +/+ and Dup/+ pups made fewer, but non-identical connections among different call types.

### Incentive Values of Neonatal Call Sequences for Maternal Approach

We previously demonstrated that the vocalizations of *Tbx1* heterozygous pups did not elicit maternal approach behavior as efficiently as those of wild-type littermates, indicating that the incentive values of call sequences of heterozygous pups were diminished compared to those of wild-type pups [61]. To evaluate how the 15q11-13 duplication alters the incentive values of neonatal vocalizations, we exposed lactating C57BL/6J mothers, 12 days postpartum, to the P12 call sequences of +/+ and Dup/+ pups using our choice tube apparatus (**Figure 2A**). The emitter utilized in this study consisted of a surface-heating thin-film electrode, a nanocrystalline silicon (ns-Si) layer, and a single-crystalline silicon wafer [72]; it efficiently generates ultrasound waves through heat transfer at the surface into the air, rather than mechanical vibration. This feature allows for the maintenance of a constant sound amplitude of up to 160 kHz [72, 73], which is crucial for replaying ultrasound vocal waves exceeding 100 kHz.

**Figure 2.**
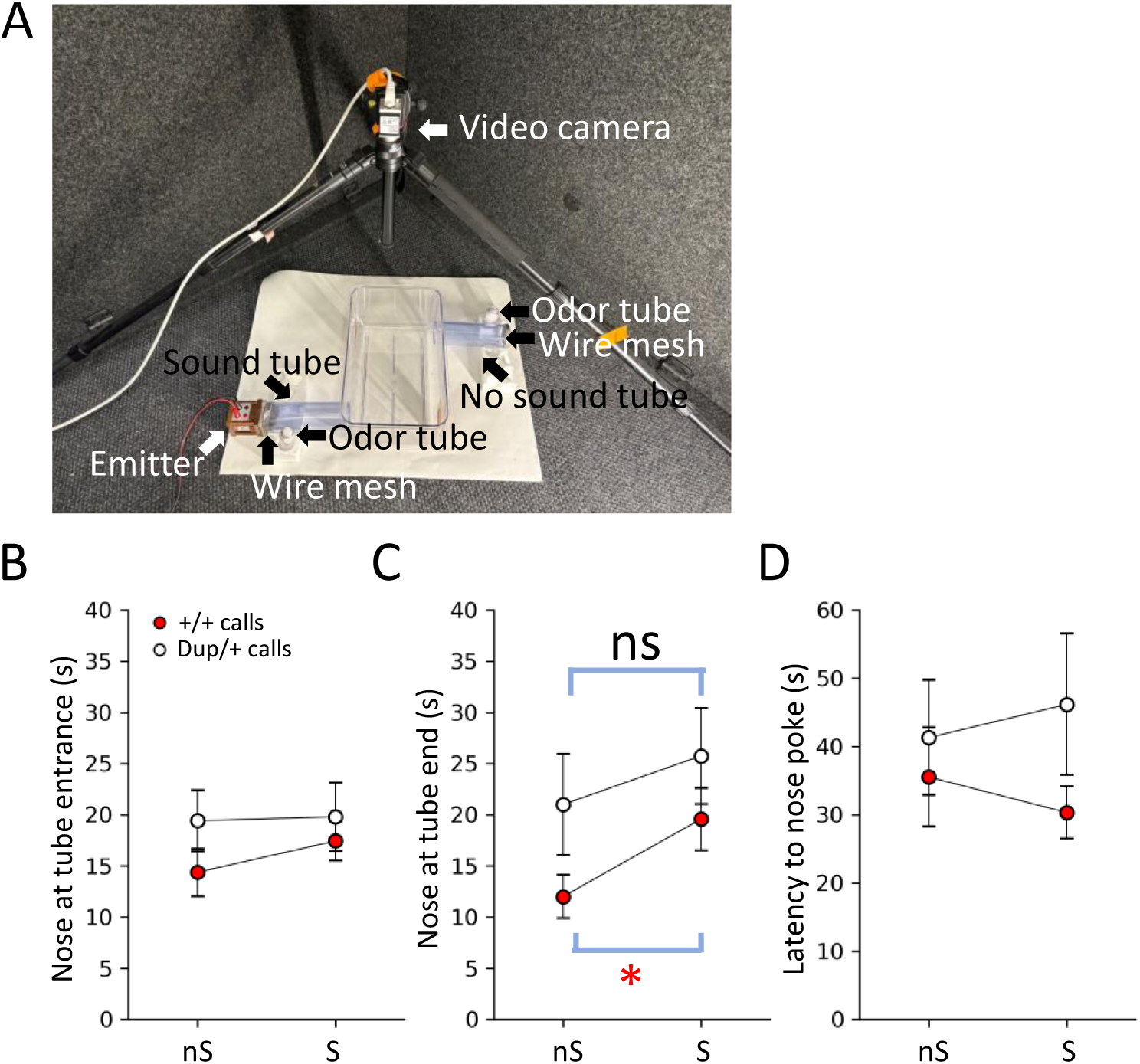
Maternal approach in response to P12 calls of +/+ and Dup/+ pups. **A)** The experimental apparatus. Slits were placed on the walls opposite the tubes to prevent reverberation of sound in the open area. The bedding odor of mothers’ own pups was placed in a plastic tube near the end of both tubes. We analyzed time the mother’s nose peeking at the entrances (**B**) and ends (**C**) of the tubes from which calls were played back (S) or no sound was played back (nS), and (**D**) latency to enter the entrances of the S and nS tubes. **B**) Mothers spent indistinguishable amounts of time peeking at the S and nS tube entrances (+/+ calls, p=0.2998; Dup/+ calls, p=0.9304), as determined by Wilcoxon tests. **C**) Mothers spent more time at the end of the S tube in which +/+ pup calls were played back, compared to the nS tube where no sound was presented (p=0.01923; p=0.0385 (*) after Benjamini Hochberg correction). Mothers spent indistinguishable amounts of time at the ends of the S and nS tubes, when Dup/+ calls were played back from the S tube (Dup/+ calls, p=0.5186). **D**) Mothers exhibited indistinguishable latencies to peek at the entrances of the S and nS tubes in response to +/+ calls (p=0.7381) or Dup/+ calls (p=0.6772). N = 20, C57BL/6J mothers exposed to +/+ calls; N=12, C57BL/6J mothers exposed to Dup/+ calls

We selected call sequences from the P12 time point, as +/+ and Dup/+ pups exhibited the most pronounced differences in the number and proportions of emitted call types at this age (see **Fig. 1A**, **S2**, and **S7**). The representative +/+ and Dup/+ pups were chosen based on their proximity to the median numbers, median proportions, median two-call sequence numbers, and median two-call proportions (see **Supplementary Figure S9-1-S9-4**). For this selection process, we excluded Markov probabilities, as they identify two-call sequences based on finite states; while useful for uncovering hidden rules of call connections, they are not optimal for selecting the most frequently occurring sequences or calls.

While C57BL/6J mothers spent comparable amounts of time peeking at the entrances of both the sound and no-sound tubes **(Figure 2B)**, they spent significantly more time exploring the end of the sound tube compared to the no-sound tube when the representative +/+ vocalizations were presented (**Figure 2C, +/+ calls**). In contrast, the mothers exhibited indistinguishable amounts of time at the ends of both sound and no-sound tubes in response to the representative Dup/+ vocalizations (**Figure 2C, Dup/+ calls**). Additionally, the latencies with which mothers approached the entrances of the sound and no-sound tubes were indistinguishable (**Figure 2D**).

To assess the significance of sound presentations and potential baseline side preferences, we measured these three parameters without call playback; otherwise, the experimental procedure remained identical to that involving sound presentations. The mothers did not display a preference between the two tubes regarding peeking time, exploration time at the ends, or latencies (**Supplementary Figure S10A, B, and C**).

In sum, +/+ calls possess greater incentive values than Dup/+ calls, as evidenced by the longer duration of exploration toward +/+ calls at the end of the sound tube than Dup/+ calls.

### Post-Pubertal Social Behavior

The Dup/+ and +/+ pups tested at P8 and P12 were subsequently evaluated for social behavior upon reaching 10 weeks of age. Molecular mechanisms underlying physical social interaction cannot be effectively recapitulated in a testing environment where a barrier prevents direct reciprocal social interaction between the experimental and stimulus mice; data obtained under such restricted conditions are not reproducible [74, 75]. Indeed, we previously demonstrated that Dup/+ mice engage in less active direct social contact with another mouse compared to +/+ mice in a test environment that allowed direct physical interaction [56]. Consequently, we employed our standard naturalistic home-cage test apparatus, facilitating direct interactions among mice, as previously described[50, 56, 59, 60, 62–65].

Male Dup/+ mice exhibited reduced levels of active affiliative social interaction compared to their +/+ littermates (**Figure 3A**). It is noteworthy that there was considerable score overlap between Dup/+ and +/+ mice, accompanied by significant variance within each genotype.

**Figure 3.**
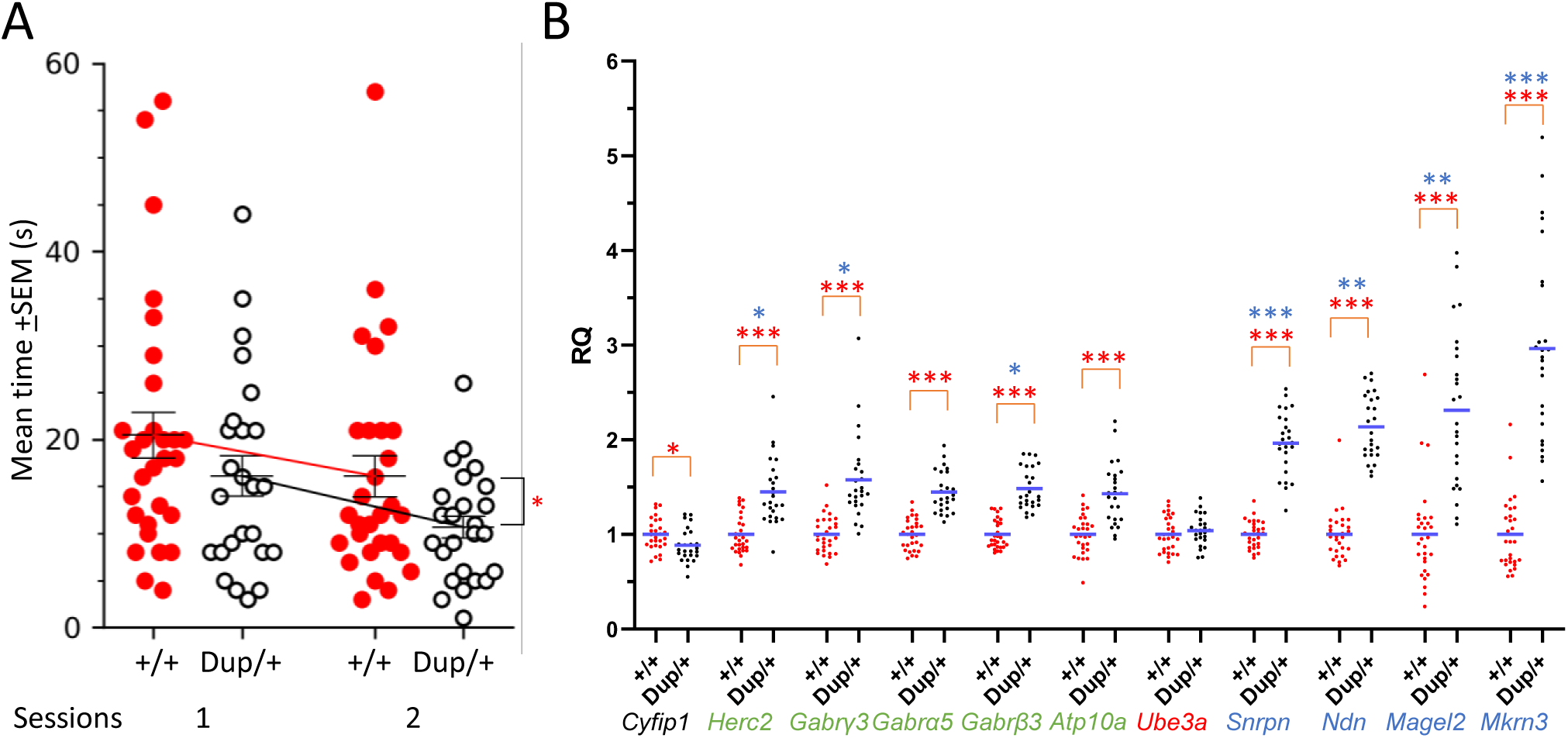
A) Post-pubertal social interaction. The time (mean±SEM) spent in active affiliative social interaction at 10 weeks of age for the +/+ and Dup/+ mice. While both genotypes equally reduced social interaction from session 1 to session 2, Dup/+ mice showed lower levels of social interaction than +/+ mice, as determined by a linear mixed effect model with the individual mouse as a random effect (genotype, F(1,52)= 4.548, p=0.038; session, F(1,52)=6.632, p=0.013; genotype × session, F(1,52)=0.083, p=0.774). For clarity, the mean±SEM of raw data are shown. +/+, n=29; Dup/+, n=25. *, p<0.05. **B**) The relative quantification (RQ) (mean+SEM) of gene expression levels of the prefrontal cortex of individual +/+ (red dots) mice and Dup/+ (black dots) mice, as determined by quantitative reverse transcription polymerase chain reaction (qRT-PCR). Statistically significant differences between +/+ and Dup/+ data are shown as *, p<0.05; **, p<0.01; ***, p<0.001, as determined by Mann-Whitney nonparametric tests; the original p values of those data that remained statistically significant after Benjamini–Hochberg correction at the false detection rate of 5% are shown (see **Table S2-**Figure 3B). The variance of gene expression was statistically significantly larger in Dup/+ than in +/+ mice for *Herc2* (*, p=0.0275), *Gabrγ3* (*, p=0.0167), *Gabrβ3* (*, p=0.0145), *Snrpn* (***, p=0.001), *Ndn* (**, p=0.0054), *Magel2* (**, p=0.0034), and *Mkrn3* (***, p=0.0004), as determined by Levene’s test for the homogeneity of variance. These p values remained statistically significant after Benjamini–Hochberg correction at the false detection rate of 5%. +/+, n=29, Dup/+, n=25.

### Expression of Mouse Ortholog Genes Encoded in 15q11-13 Copy Number Variation in the Brain

The observed expression levels of paternally and maternally inherited genes within the 15q11-13 region do not necessarily correspond to the expected levels in the brain tissues of humans [76] and mice [56]. However, the gene expression in the brain and behavioral variability have not been correlated in the same individuals or mice. We thus assessed the expression levels of ten protein-coding genes corresponding to the murine ortholog of the 15q11-13 region following an evaluation of social behavior at ten weeks of age in the same mice. The prefrontal cortex and the remaining forebrain (excluding the olfactory bulb) were utilized to quantify gene expression levels via quantitative reverse transcription polymerase chain reaction (qRT-PCR). Our primary analysis focused on the prefrontal cortex, given its well-established functional significance in social behavior in mice [77–81].

Expression levels of all genes, except for *Ube3a* and *Cyfip1*, were significantly elevated in the prefrontal cortex of Dup/+ mice compared to their +/+ littermates (see **Figure 3B**). The absence of elevated expression of *Ube3a* in Dup/+ mice, as it is a maternally inherited gene, with its paternal copy silenced by genomic imprinting. Similarly, *Cyfip1* is located outside the duplicated segment, rendering its overexpression unexpected. Non-imprinted genes (*Herc2*, *Gabrg3*, *Gabra5*, *Gabrb3,* and *Atp10a*) are expressed from both paternal and maternal copies in +/+ mice, leading to expression levels of approximately 1.5 relative quantification (RQ) from three copies in paternal Dup/+ mice. Although *Atp10a* was previously presumed to be maternally inherited due to paternal imprinting, evidence indicates that it is expressed from both paternal and maternal copies in the embryonic, neonatal, and adult brains of mice and humans [82–84]. The paternally inherited genes (*Snrpn, Ndn*, *Magel2*, and *Mkrn3*) are expressed as a single copy in +/+ mice, as the maternal copy is silenced by genomic imprinting. Consequently, their expression levels reached approximately 2.0 RQ in paternal Dup/+ mice. Notably, the expression levels of *Mkrn3* were significantly higher than anticipated based solely on duplication. Furthermore, the expression levels of the paternally inherited genes exhibited greater variability than those in +/+ mice (**Figure 3B**).

In the remaining forebrain, all genes, except for *Cyfip1* and *Ube3a*, demonstrated increased expression levels in Dup/+ mice relative to +/+ littermates (see **Figure S11**). Similar to the findings in the prefrontal cortex, *Mkrn3* expression levels were markedly higher than expected based solely on duplication (i.e., RQ=2.0). In comparison to +/+ mice, individual Dup/+ mice exhibited significantly greater variance in the expression of paternally inherited genes in the remaining forebrain (**Figure S11**).

### Predicting Post-Pubertal Social Interaction Scores through Neonatal Social Communication Variables and Post-Pubertal Brain Gene Expression

We previously identified predictive neonatal vocalization parameters for post-pubertal social behaviors in a mouse model of 16p11.2 hemizygous deletion [59]. Given the expression variability of genes located in the 15q11-13 ortholog, alongside variable neonatal social communication, we aimed to investigate the predictive capacity of these two sets of parameters regarding post-pubertal social behavior. If neonatal social communication serves as a prerequisite and brain gene expression acts as a determinant for the development of post-pubertal social behavior, then variability in these neonatal and brain gene expression parameters would predict variability in post-pubertal social behavior.

We aggregated the frequency of each call type (**Figure 1A**), the proportions of each call type (**Figure S2**), the frequencies and proportions of two-call connections (**Figure S7**), the Markov probabilities of two-call connections (**Figure 1C**) at postnatal day 12, and expression levels of genes encoded within the 15q11-13 region in the prefrontal cortex at 10 weeks of age (**Figure 3B**). Subsequently, we employed Lasso regression models to identify predictors of individual variability in post-pubertal social interaction scores. To ascertain the parameters necessary for an optimal model fit that balances maximum likelihood with the risk of overfitting, we utilized the Akaike Information Criterion (**Figure S12**). Following the establishment of cutoff points yielding the best model fit, Lasso regression models elucidated predictors among gene expression and neonatal call metrics (**Figure 4**).

**Figure 4.**
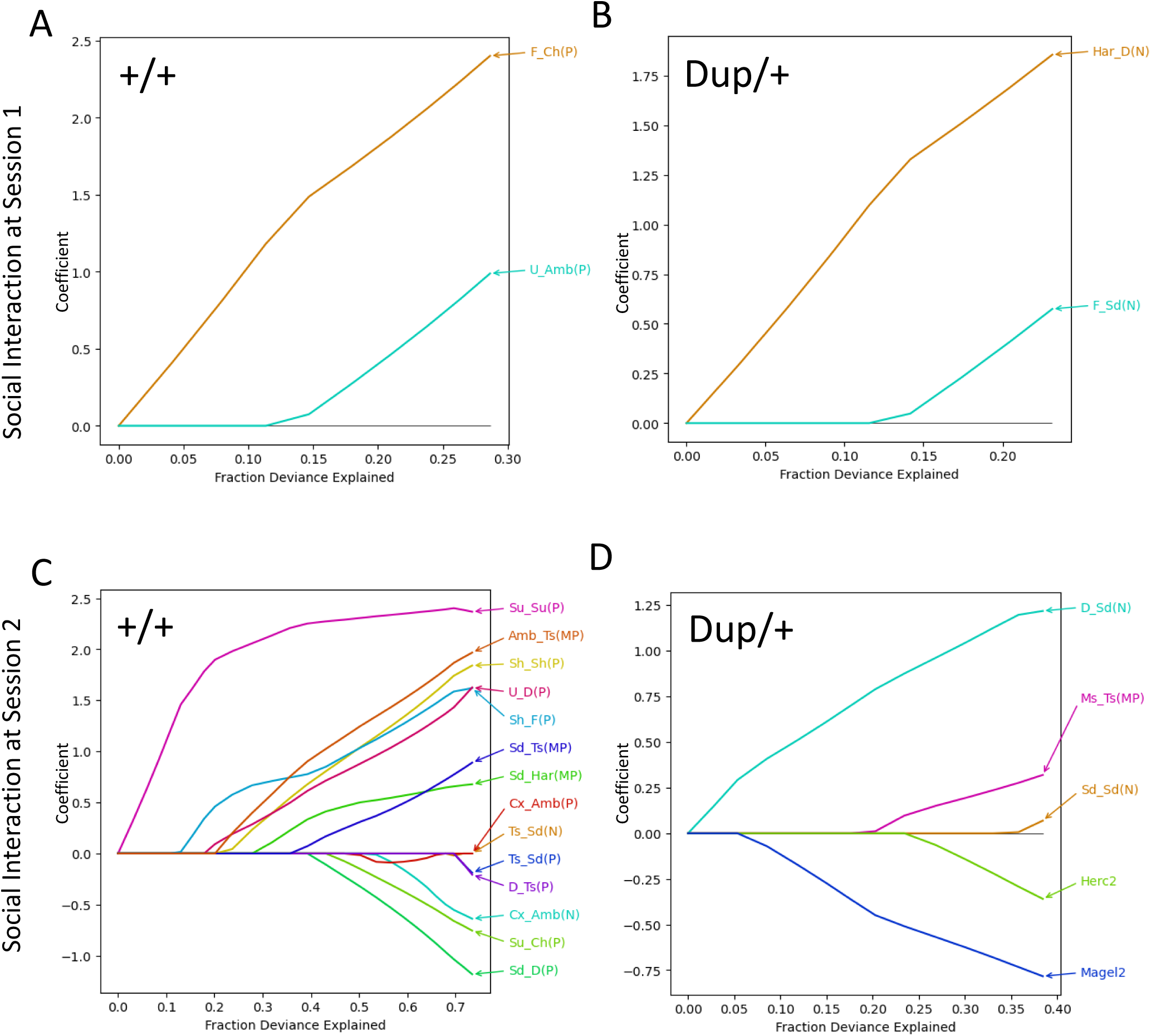
Fraction deviance explained, and coefficients of variables determined by the Lasso regression models of +/+, social interaction session 1 (**A**), Dup/+, session 1 (**B**), +/+, session 2 (**C**), and Dup/+, session 2 (**D**). The colors of lines are arbitrarily assigned. The number (N) and proportions (P) of each call type, number (N) and probabilities (P) of two-call sequences, Markov probabilities (MP) of two-call sequences, and gene expression scores were pooled and used for selection. We used call parameters of P12, as +/+ and Dup/+ pups showed more robust differences in calls on this day than on P8 (see Figure 1A). The dependent variable was affiliative social interaction scores. The fraction of variance explained was chosen and displayed, based on Akaike’s information criteria (see **Figure S12**). Har, harmonic; Su, step-up; Ts, two-steps; Cx, complex; Sd, step-down; Ms, multiple-steps; D, downward; Ch, chevron; Sh, short; F, flat; U, upward; R Ch, reverse chevron; Amb, ambiguous.

The probabilities of transitioning from flat call types to chevron call types (F_Ch(P)) and from upward transitions to ambiguous call types (U_Amb(P)) emerged as predictors for the social interaction scores of +/+ mice during the first social interaction test session (**Figure 4A**). The frequencies of transitions from harmonic to downward calls (Har_D(N)) and from flat to step-down call types (F_Sd(N)) served as predictors for the social interaction scores of Dup/+ mice at session 1 (**Figure 4B**).

More predictors were identified for social interaction scores during session 2. Notably, various probabilities, frequencies, and Markov probabilities of two-call transitions significantly predicted the social interaction scores of +/+ mice (**Figure 4C**). Interestingly, the social scores of Dup/+ mice were predicted by the expression levels of *Magel2* and *Herc2* in the prefrontal cortex, in addition to the frequencies of transitions from downward to step-down call types (D_Sd(N)) and from step-down to step-down call types (Sd_Sd(N)), as well as the Markov probability of multiple steps to two steps (Ms_Ts(MP)) (**Figure 4D**).

Given the sample sizes, a simple cross validation for models is not suitable [85]. A more appropriate approach to confirm the validity of the identified predictive parameters is to eliminate a subset of data from the original data sets and determine how many times the identified predictors appear. We thus used the 5-fold cross validation procedure by creating five smaller data sets after randomly excluding 5-6 mice from the original data set five times and running Lasso analyses. Among gene expression predictors, *Magel2* and *Herc2* appeared three times out of 5 repeats at Session 2 in Dup/+ mice (**Figure S13D, Fold 1-Fold 5**). Vocalization parameters that appeared three time or more were F_Ch(P) and U_Amb(P) in +/+ and Har_D(N) and F_Sd(N) in Dup/+ at session 1; Su_Su(P), Amb_Ts(MP), U_D(P), Sh_F(P), Sd_Ts(MP) in +/+ at session 2 (**Figure S13A, B, C, Fold 1-Fold 5**). It is interesting to observe that the social behavior of Dup/+ at session 2 were better predicted by gene expression than neonatal vocalizations, whereas the opposite were the case for all the other three cases: +/+ at sessions 1 and 2 and Dup/+ at session 1.

We further validated gene expression predictors by analyzing correlation coefficients (**Figure S14**). The expression levels of *Magel2*, *Herc2*, and *Ndn,* in the prefrontal cortex of Dup/+ mice at session 2 exhibited the highest correlation coefficients , with higher expression of these genes being associated with lower social interaction scores among individual Dup/+ mice.

Interestingly, the predictors differed between sessions in both +/+ and Dup/+ mice. This result is not unexpected, as social interaction predominantly encompasses aggression, stress, and anxiety during the initial encounter, while memory-based familiarity and recognition, along with established hierarchy, come into play during Session 2 [86].

## DISCUSSION

Our computational approach demonstrates that individual variability in post-pubertal social behavior arises from call sequences during the neonatal period and correlates with varying expression levels of 15q11-13-encoded genes. This methodology provides a technical framework for predicting the developmental trajectories of social behaviors from the neonatal to post-pubertal stages in both individual mice and humans affected by this and other CNVs. Furthermore, while assessing the variability of CNV-encoded gene expression in the human brain presents significant challenges, our complementary approach in a mouse model facilitates the identification of CNV-encoded driver genes and their associated molecular pathways.

Mouse pups typically exhibit more than ten distinct call types (**Figure S1**), frequencies of which are variably impacted by gene dose alterations associated with neurodevelopmental disorders [59, 61, 65, 68, 87, 88]. The current study extends our previous finding that 15q11-13 Dup/+ pups produce a greater number of calls than +/+ mice by demonstrating that this phenotype is a net result of increased numbers or proportions of certain call types (harmonic, step-up, two-steps, step-down, multiple-steps, downward, chevron, and ambiguous) and decreased numbers or proportions of others (short, upward, and reverse chevron).

Although Dup/+ and +/+ pups maintained a consistent selection of call types within each genotype, as indicated by indistinguishable entropy scores between the two groups at P12 (see **Figure S5**), the genotypes differed significantly in specific call types emitted (see **Figure 1A, 1B**; **Figure S2**), call sequences (**Figure S7**), and finite state-based rules of call sequences (**Figure 1C**). Both the proportion of calls (**Figure S7**) and Markov-type call connections (**Figure 1C**).

The classification of calls based on their acoustic properties identified four major distinct groups (see **Figure 1B; Figures S3-1** to **S3-4**). These clusters only partially accounted for the qualitative classification of call types, as many distinct call types shared similar quantitative properties within each cluster, while others exhibited distinct quantitative characteristics. Moreover, despite qualitative similarities, the quantitative properties of the same call types could differ across different clusters (see **Fig. 1B**; **Fig S3**). Nonetheless, these quantitative differences did not appear to correlate with genotype (**Figure 1B; Figure S3-1** to **S3-4**), suggesting that quantitative, acoustic properties of call sounds were less effective than qualitative properties in detecting genotype-dependent phenotypes.

We evaluated the biological significance of these genotype-dependent properties of neonatal vocalizations. The representative call types and sequences of the Dup/+ genotype were ineffective in eliciting maternal search behavior toward the sound source, as evidenced by the time spent at the end of the sound tube (**Figure 2C**). Previous work with *Tbx1* heterozygous pups indicated that the time spent peeking at the entrance of the sound tube was influenced more by call types than by call sequences; randomized wild-type calls elicited similar maternal responses to the original calls.

However, the randomized wild-type calls failed to sustain maternal search behavior (i.e., time spent at the end of the sound tube) or to prompt a rapid approach to the tube entrance (i.e., latency to approach the sound tube entrance)[61]. These data suggest that call types dictate the initial selection of the tube from which pup calls are played, while call sequences influence the persistence of search behavior in the sound tube and the speed of approach to the source of pup calls. Given the differential reliance of various aspects of maternal approach behavior on components of vocalization, it is significant that the calls of Dup/+ pups were ineffective in eliciting sustained search behavior for the call source. In contrast, neither +/+ nor Dup/+ calls resulted in statistically significant peeking at the entrance or rapid approaches to the sound tube entrance (i.e., latency), although there were non-significant trends towards the +/+ sound tube (see **Figure 2B** and **D**). The incentive values of call sequences of Dup/+ pups seem to be diminished.

The expression levels of duplicated mouse genes orthologous to the human 15q11-13 duplication exhibited variability among individual mice (see variance in **Figure 3B**). Our Lasso regression analysis identified *Magel2* and, to a lesser extent, *Herc2* as potential predictors of individual post-pubertal social interaction scores in Dup/+ mice (**Figure 4D**). These parameters appeared three times in the 5-fold validation despite a potential bias due to the small sample size (**Figure S13, Fold 1-5**). Both *Magel2* and *Herc2* expression levels were negatively correlated with post-pubertal social interaction (see **Figure S14**). Although *Ndn*, another paternally-inherited gene, was not identified by the Lasso regression analysis, its individual expression levels similarly exhibited a high negative correlation coefficient with social interaction scores in Dup/+ mice (**Figure S14**), consistent with our previous findings demonstrating that normalization of *Ndn* restores certain parameters of social behavior in a mouse model of 15q11-13 duplication [89]. Our preclinical data are consistent with human findings that protein-truncating variants of *MAGEL2* and *HERC2* are present in non-CNV carriers with idiopathic cases of autism spectrum disorder (ASD) [8] and schizophrenia[10], although these variants have not consistently reached statistical significance due to their rare nature.

*Magel2* in the prefrontal cortex emerged as the most robust predictor of social interaction, with its degree of overexpression negatively correlating with social interaction levels in session 2 when Dup/+ mice exhibited lower levels of social interaction compared to +/+ mice. A mouse model with overexpressed *Magel2* but normalized *Ndn* demonstrated social interaction levels indistinguishable from wild-type mice, yet remained highly reactive to a stranger mouse[89]. Moreover, isolated normalization of *Ndn* alone did not fully restore the higher number of neonatal vocalizations in our paternal 15q11-13 duplication model [89]. Thus, the role of *Magel2* as an additional potential driver gene for specific parameters of social behavior cannot be discounted. Indeed, the critical role of *Magel2* in social behavior is further substantiated by its deletion, as *MAGEL2* deletion is a recognized risk factor for ASD, intellectual disability, and epilepsy in humans [90]. Paternally inherited *Magel2* deletion in mice adversely affects neonatal vocalizations [91], various aspects of post-pubertal social behavior [92–94] and parenting behavior[95].

Conversely, overexpression of a 1.5 Mb segment encompassing *Ube3a* and *Snrpn* does not influence social interaction in mice[89]. Our data align with this observation, as neither gene was identified as a predictor (see **Figure 4D**). A mouse model with an additional copy of *Ube3a* does not exhibit any phenotypic abnormalities in post-pubertal social behavior; however, a model with two extra copies of *Ube3a* does exhibit such abnormalities, despite neonatal vocalizations remaining unaltered in both models [96, 97]. Another model with four extra copies of *Ube3a* showed no deficits in social behavior [98]. This apparent lack of a dose-dependent phenotype may, in part, be attributable to genetic background, as the former [96] and latter [98] models were generated from FVB and C57BL/6J mice, respectively. Different inbred mouse models exhibit varying susceptibility to gene-dose alterations[3, 4]. Alternatively, the absence of phenotype in the three-chamber apparatus in the four-extra-copy model may result from inherent limitations of this task, which does not directly measure social interaction [75, 99].

We acknowledge the potential for false-negative cases due to conceptual and technical limitations of our approach. Our strategy capitalized on the variance of gene expression levels, which are likely to exhibit higher selection sensitivity for larger variances. However, while *Mkrn3* expression demonstrated significantly higher variance than *Herc2* in the prefrontal cortex of Dup/+ mice, only *Herc2* was selected in our predictive models. Hence, the magnitude of variance alone does not fully account for the outcomes of our predictive models. Although *Herc2* is a non-imprinted gene, its overexpression level might determine the level of social interaction in Dup/+ mice, together with *Magel2*.

It should be noted that technical limitations constrain our interpretation of the qRT-PCR-based approach. Gene expression variability may not be reliably assessed for genes with basally low levels of expression compared to those with high basal expression levels. Despite this interpretative limitation, our computational approach is still applicable to other CNVs for identifying predictors among abundantly expressed genes and neonatal parameters related to social and other behaviors. Some genes expressed in a mouse model of 16p11.2 hemizygous deletion exhibit greater inter-individual variability compared to other encoded genes [59]. Similarly, a mouse model of 22q11.2 hemizygous deletion displays individual variability in gene expression[100].

Our analysis focused on the prefrontal cortex due to its established functional relevance to social behavior [101], but can be applied to other brain regions implicated in other dimensions, such as motor and cognitive capacities, of mental illness. Regional and single-cell gene expression within various brain regions is expected to elucidate the circuits and networks and specific cell types through which driver genes influence social and other behaviors in this and other genetic mouse models of CNVs.

Moreover, our approach can be effectively applied to single-gene cases of psychiatric disorders, as gene expression variance may account for incomplete penetrance and variable expressivity of behavioral dimensions in such instances.

Although neonatal social communication precedes post-pubertal social behavior, the question of whether neonatal social communication is a prerequisite for the development of post-pubertal social behavior has not yet been experimentally evaluated. In this study, a gene-dose alteration of 15q11-13 affected both the types and sequences of neonatal vocalizations as well as post-pubertal social interactions at the individual mouse level. The phenotypic correlations observed across these two developmental time points are consistent with the hypothesis that neonatal social communication is essential for the later development of social behavior.

On the other hand, predictive parameters in gene expression varied between genotypes. Levels of *Magel2*, *Herc2*, and *Ndn* were predictive of individual post-pubertal social interaction levels in Dup/+ mice, whereas none of 15q11-13-encoded genes were correlated with the variability of post-pubertal social interaction in +/+ littermates (see **Figure S14**). This finding raises the intriguing possibility that the variabilities in social interaction among controls and mutants depend on distinct sets of genes. A corollary of this observation is that, although social interaction scores partially overlap between genotypes, the underlying genes responsible for such an overlap may differ between genotypes. As the functional roles of a gene in each phenotype might differ in wild-type and mutant mice, caution is needed in extrapolating gene functions between mice with and without risk gene variants.

Paternally inherited 15q11-13 duplication does not confer complete penetrance. Although it is challenging to estimate the precise percentages of currently identified cases, approximately half of the carriers exhibit a composite phenotype that variably includes intellectual disability, developmental delay, ASD, speech delay, and behavioral problems, while one-third of carriers do not exhibit any of these diagnoses[33]. Severity also varies among individual carriers of paternal 15q11-13 duplication within each diagnosis. Such phenotypic variability cannot be accounted for solely by the chromosomal breakpoints of individual carriers[102]. Our computational approach with a preclinical mouse model is a complementary method to identify the sources of phenotypic variability, thereby contributing to a more comprehensive mechanistic understanding of psychiatric disorders and the development of gene therapy strategies targeting dimensions of mental illness.

## Acknowledgments

Research reported in this publication was partly supported by the Japanese Society for the Promotion of Science KAKENHI (23K06399) and Takeda Science Foundation to K.T.; Japanese Society for the Promotion of Science KAKENHI (23KK0132, 23H04233, 24K22036, 24H00620, 24H01241), Japan Agency for Medical Research and Development (JP21wm0425011), and Japan Science and Technology Agency (JPMJMS2299, JPMJMS229B), Takeda Science Foundation to T.Takumi; National Institutes of Health (R01DC015776, R01MH099660, R03HD108551, R21HD105287) to N.H; and Senshin Medical Research Foundation and the Uehara Memorial Foundation to T.Tanifuji. The content is solely the responsibility of the authors and does not necessarily represent the official views of the National Institutes of Health. The open-access license has been selected.

## Author contributions

1. N. Hiroi and T. Takumi designed the entire study.
2. T. Yamauchi and T. Hiramoto designed and performed qRT-PCR and analyzed data.
3. K. Tamada tested mice for neonatal vocalization and social interaction, prepared brain samples, and genotyped mice.
4. T. Takano, M. Nakamura, L. Stevens, K. Ye, H. Inada, M. Barbachan, T. Tanifuji, and P. Ó Broin statistically and computationally analyzed data and made figures.
5. N. Hiroi wrote the manuscript.
6. K. Tamada, T. Yamauchi, H. Inada, and N. Osumi participated in the writing.
7. G. Kang and M. Esparza managed the technical aspects of mice and sample preparation.
8. T. Kikusui provided and repaired the emitter and designed the recording apparatus for maternal approach.

## Data availability

The datasets generated and analyzed during the current study are available from the corresponding author on reasonable request.

## METHODS

### Ultrasonic vocalization recording

The apparatus and procedure are detailed in our previous publications (1-3). Briefly, male pups were tested for vocalization during 5-minute maternal separation at postnatal day (P) P8 and P12 in a new home cage kept at 23°C to 26°C under a 30-lux light; these postnatal mouse ages correspond to term human infants (4). Testing took place during the light phase between 8:00 AM and 8:00 PM. The test cage was placed in a soundproof box. Ultrasonic vocalizations_were recorded by UltraSoundGate (Avisoft, Germany) connected to a computer equipped with Avisoft-RECORDER software (Avisoft, Germany). We did not set a cut-off threshold for analyzing the frequency range.

### Call type classification

We used VocalMat to classify call types, as this analytical software has the lowest false discovery rate (5%) compared with other published software (5). The outputs of the segmentation process of ultrasonic vocalization candidates were manually labeled in accordance with those of Scattoni et al. (6) and are thus compatible with our previous analyses (2). This software classifies the ultrasonic vocalization candidates into the following call types: complex, step-up, step-down, two-steps, multiple-steps, upward frequency modulation, downward frequency modulation, flat, short, chevron, reverse chevron, and noise. VocalMat tends to classify calls based on the most prominent component but flags them as potential harmonic call types. We manually added harmonics by inspecting the whole spectrum of each call instead of the most prominent component. Calls that were close to a call type but did not satisfy at least one criterion of the call classification were classified as ambiguous (see **Figure S1**).

### Social interaction

Each 10-week-old +/+ or Dup/+ mouse and a stranger C57BL/6J mouse of the same age (±1 week) were randomly selected and were simultaneously placed in a novel test box together (37 cm long × 26 cm wide × 19 cm high; 30 lux) and left there during two 5-minute sessions with a 30-minute inter-session interval in accordance with our published protocol (3, 7). An acrylic plate was placed on the top of the cage during testing. Social behavior was monitored by a CCD camera connected to a Macintosh computer. Behavior was manually rated in accordance with our published procedure (1, 3, 7-11). The number and duration of affiliative social interactions were measured and used for analysis.

### Quantitative reverse transcription polymerase chain reaction (qRT-PCR)

Up to 4 to 5 days following social interaction testing at 10 weeks of age, the mice were euthanized, and the prefrontal cortex, along with the remainder of the forebrain, excluding the olfactory bulb, cerebellum, and brainstem, was extracted. The intervals between social interaction testing and euthanasia were comparable for the +/+ and Dup/+ groups (+/+, Average = 4.8 days (SEM = 0.385); Dup/+, Average = 5.2 days (SEM = 0.445)). A Mann-Whitney U test revealed no significant difference (U = 316, p = 0.428).

qRT-PCR was carried out in accordance with our published procedure (12). Total RNA was extracted from the brain regions using an RNeasy Plus Mini Kit (Cat#74134, Qiagen, Germantown, USA). Complementary DNA (cDNA) was synthesized from total RNA using SuperScript IV VILO Master Mix (Cat#11766050, Invitrogen, Carlsbad, USA). qRT-PCR reactions were performed in triplicate on a QuantStudio 6 Flex Real-Time PCR System (Cat#4485694, Applied Biosystems, Waltham, USA) using the TaqMan Fast Advanced Master Mix (Cat#4444963, Applied Biosystems, Waltham, USA). The Taqman probes are listed in the Supplementary Material (**Table S1**). Data were analyzed using the ΔΔCt method and normalized to the reference gene *18S*.

### Uniform manifold approximation and projection (UMAP)

We used the Python library “umap-learn” (version 0.5.5) to classify calls based on the quantitative parameters of VocalMat, including the length (duration) and bandwidth of each call, the minimum and maximum frequency values in kHz (min_freq_main, max_freq_main, mean_freq_main, min_freq_total, max_freq_total, mean_freq_total), and the sound intensity in dB (min_intens_total, max_intens_total, mean_intens_total) of various components of each call, where “main” and “total” designate the most intense wave component and all wave components, respectively, of each call. We then added call types to each data point in UMAP clusters (13).

### Call sequence analysis

A call sequence was defined as a series of calls with inter-call intervals below the intersection between the theoretical and observed distribution curves of inter-call intervals (2, 3, 13). Accordingly, two calls with longer inter-call intervals than the cross point of the two curves are the last and first calls of two distinct call sequences (**Figure S4**).

### Shannon entropy analysis

We applied Shannon entropy analysis to call sequences of up to length four to determine their overall structure of unpredictability. Statistical analysis of entropy data was carried out using a mixed linear model, with entropy being modeled as a function of both group and level, with each pup having its baseline entropy (**Figure S5**).

### Non-Markov sequences

The standard proportions of two-call pairs within so-defined sequences were also analyzed using our published procedure (2, 3, 13) (**Figure S6**).

### Markov modeling

Two-call pairs within so-defined sequences were analyzed with Markov modeling using our published procedure (2, 3, 13).

### Least absolute shrinkage and selection operator (Lasso) model

We extracted predictive variables from the numbers and proportions of each call type and two-call pairs within sequences for individual social interaction scores using the Lasso regression model (3, 13). We validated the identified predictors by generating five smaller sample sets through the random exclusion of five or six mice, followed by the execution of Lasso regression models on each set.

### Correlation coefficients

The correlation coefficients between the relative expression levels of each gene encoded in our mouse model of paternally inherited 15q11 duplication and the social interaction scores were calculated and their significance was determined.

### Maternal approach

A Plexiglas mouse home cage was modified to measure maternal approach to neonatal vocalizations, using our published apparatus and procedure (2) (see **Figure 2A**). It was stationed in a sound-proof booth. One nanocrystalline silicon (nc-Si) sound emitter was placed facing the wire mesh net-covered end of one of the two tubes. The odor of bedding was collected on a cotton ball, on which three randomly chosen pups from the litter were placed for 3 hrs. The cotton ball was placed in a plastic tube near the emitter and the mesh. Ultrasonic vocal calls were played back using a nanocrystalline silicon (nc-Si) emitter, composed of a surface-heating thin-film electrode, an ns-Si layer, and a single-crystalline silicon wafer. The electrical signals generated were supplied as voltage inputs through a high frequency amplifier to more faithfully reproduce pup vocalizations of high frequencies than commercially available vibration-based speakers. To evaluate the accuracy of sound emitted from our device, ultrasonic sound was monitored by a condenser microphone (CM16/CMPA, Avisoft Bioacoustics, Germany). An amplifier (UltraSoundGate116H, Avisoft Bioacoustics, Germany) analyzed data using an analog-to-digital converter, frequency filters, digital first-Fourier-transform analysis, and signal input-output terminals. Input signals were monitored with spectrum software (Avisoft-SASLab Pro 3.0, Avisoft Bioacoustics, Germany).

We chose calls of a single representative P12 +/+ pup and a single representative P12 Dup/+ pup. The representative calls of each genotype were defined as follows: We computed the difference scores between each pup and the group median in the percentages an numbers of each of the thirteen call types; the absolute values of the difference scores were used to reflect a deviation from the median regardless of the direction (i.e., higher or lower than the median); these scores were summed for all call types. The pup with the smallest sum score was chosen from each genotype (**Figure S9-1** to **S9-4**). The calls of the representative +/+ pup and Dup/+ pup were played back.

This approach was the most practical and feasible one for two reasons. First, calls from many pups cannot be sequentially presented to a single female mother due to a rapid habituation to repeated presentations of calls(14). Second, our pilot study showed that each call sample requires 10 or more mothers to detect statistical significance due to inherently variable responses among different mothers.

Male and female C57BL/6J mice (#000664, Jackson Laboratory, Bar Harber, ME) were paired and pregnant female mice were separated singly when a vaginal plug was noted. The lactating mother was used at 12 day postpartum. We used mothers that had up to four litters, as there was no correlation between the number of litters they produced and the maternal approach parameters.

The basic procedure is detailed in our published work (2): Day 1, habituation in the home cage in the soundproof booth overnight; Day 2, a 5-min test day. Mothers were placed in the test apparatus for 30 min in the soundproof room, and exposed to the representative +/+ or Dup/+ calls. Calls were played back at the end of one of the two tubes; no sound was presented at the other tube. The mother mouse was allowed to freely choose, move, and stay anywhere in the apparatus during the 5-min test. After testing, the mother was returned to the home cage and placed back to the mouse room.

The mother’s behavior was recorded by a video camera. Recorded behavior was classified by the Noldus software (Noldus Information Technology, Leesburg, VA) into the time the mother’s nose peeked at the entrance of each tube, time the mother’s nose stayed at the end of the sound tube (i.e. closest place to the emitter), and the latency to poke the nose to the tubes.

---

**Supplementary Figure S1.**
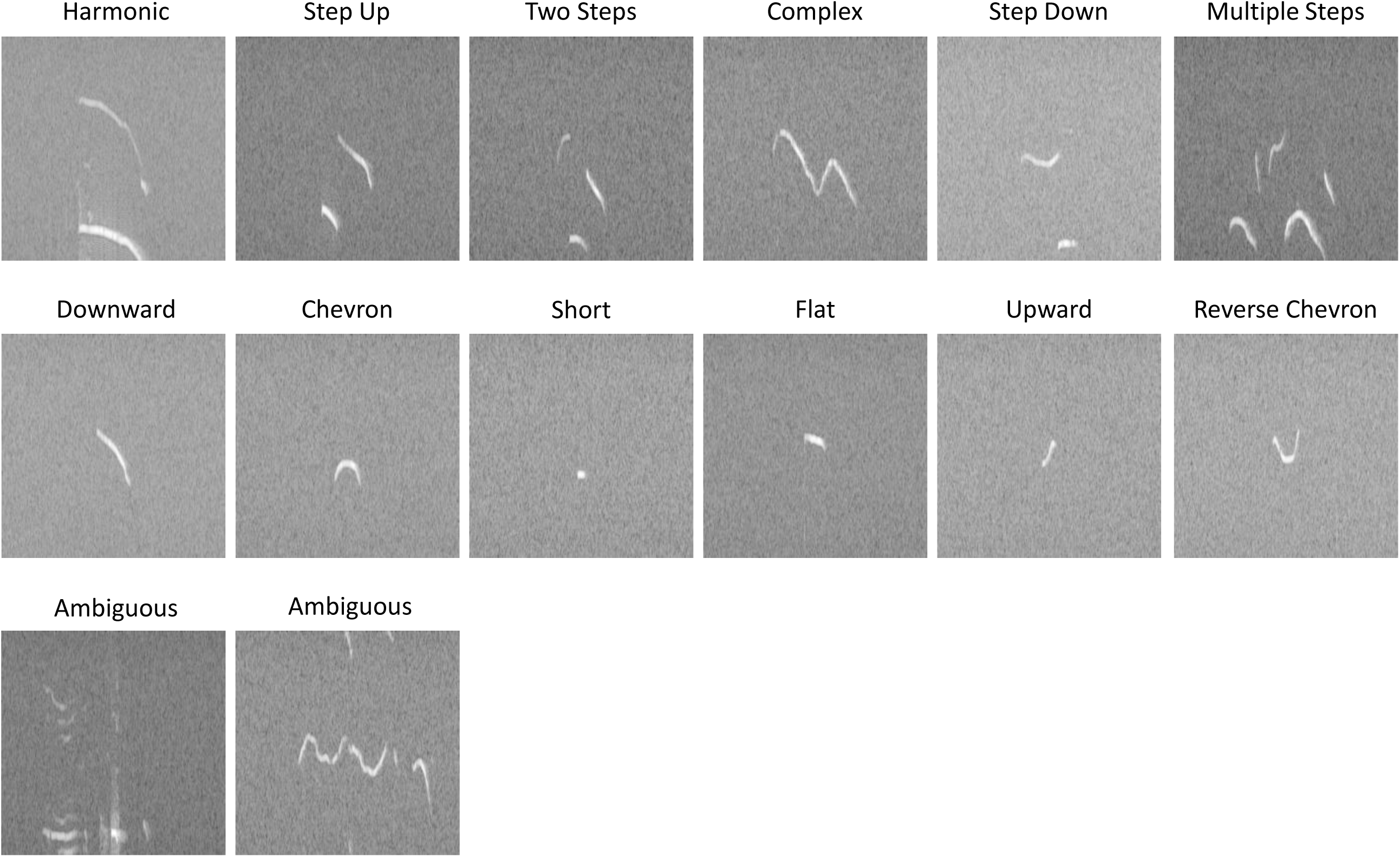
Representative sonogram images of each call type. A modified VocalMat was used to classify pup calls.

**Supplementary Figure S2.**
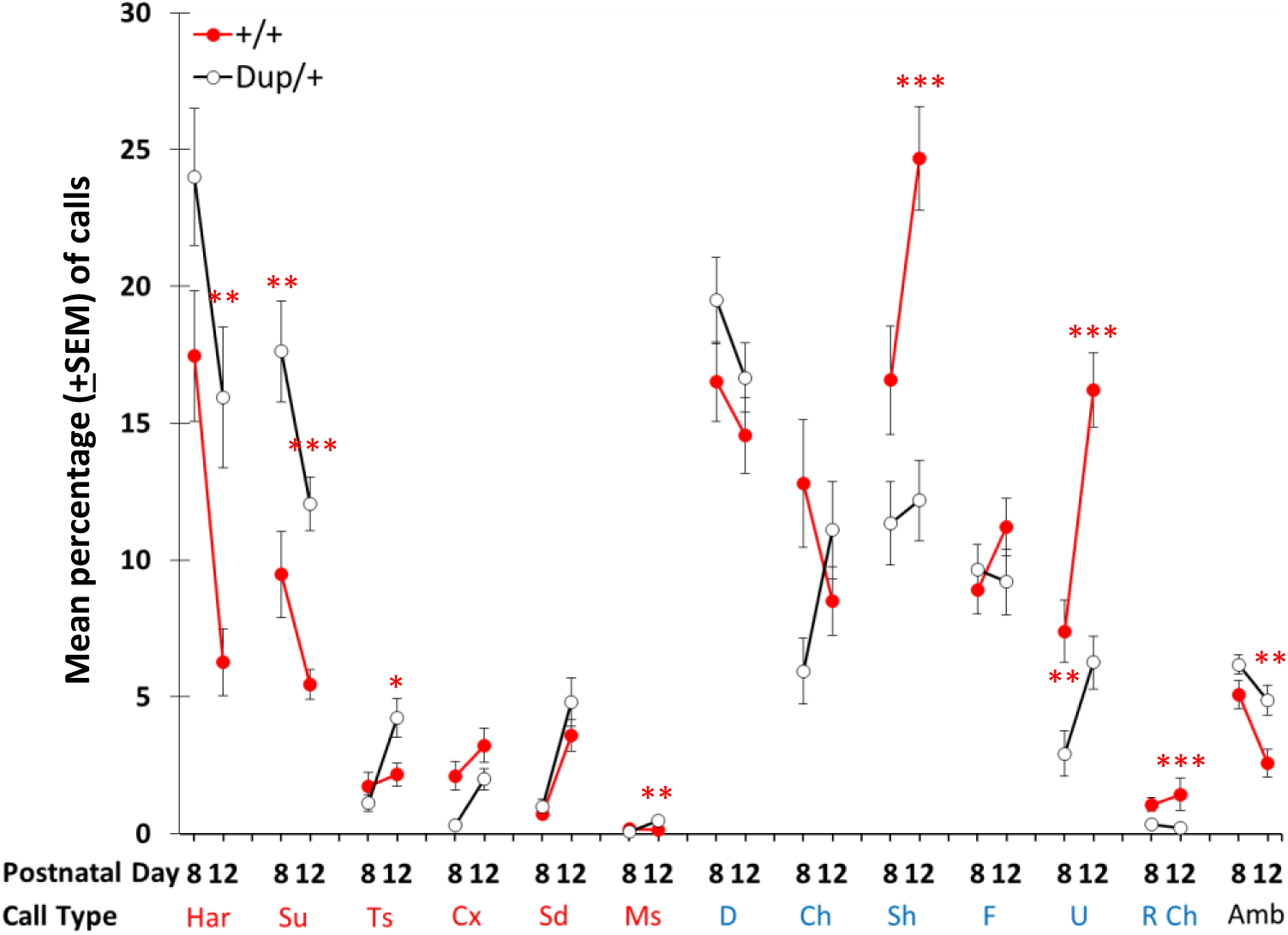
The percentage (mean±SEM) of each call type emitted for Dup/+ (white circle with black line) and +/+ (red) pups P P8 and P12. The assumptions of normality and homogeneity of variance were violated in many cases (see **Table S1-Figure S2**). We applied Mann-Whitney nonparametric tests, and significance was adjusted by Benjamini–Hochberg corrections with a false discovery rate of 5%. Har, harmonic; Su, step-up; Ts, two-steps; Cx, complex; Sd, step-down; Ms, multiple-steps; D, downward; Ch, chevron; Sh, short; F, flat; U, upward; R Ch, reverse chevron; Amb, ambiguous. Red call type names are those of more than one wave or complex waves; blue call type names are those of relatively simple waves(2, 3). P8: +/+, n=29; Dup/+, n=25. P12: +/+, n=28; Dup/+, n=25. *, p<0.05; **, p<0.01. One +/+ pup emitted no call at P12 and call percentages were not calculated.

**Supplementary Figure S3.**
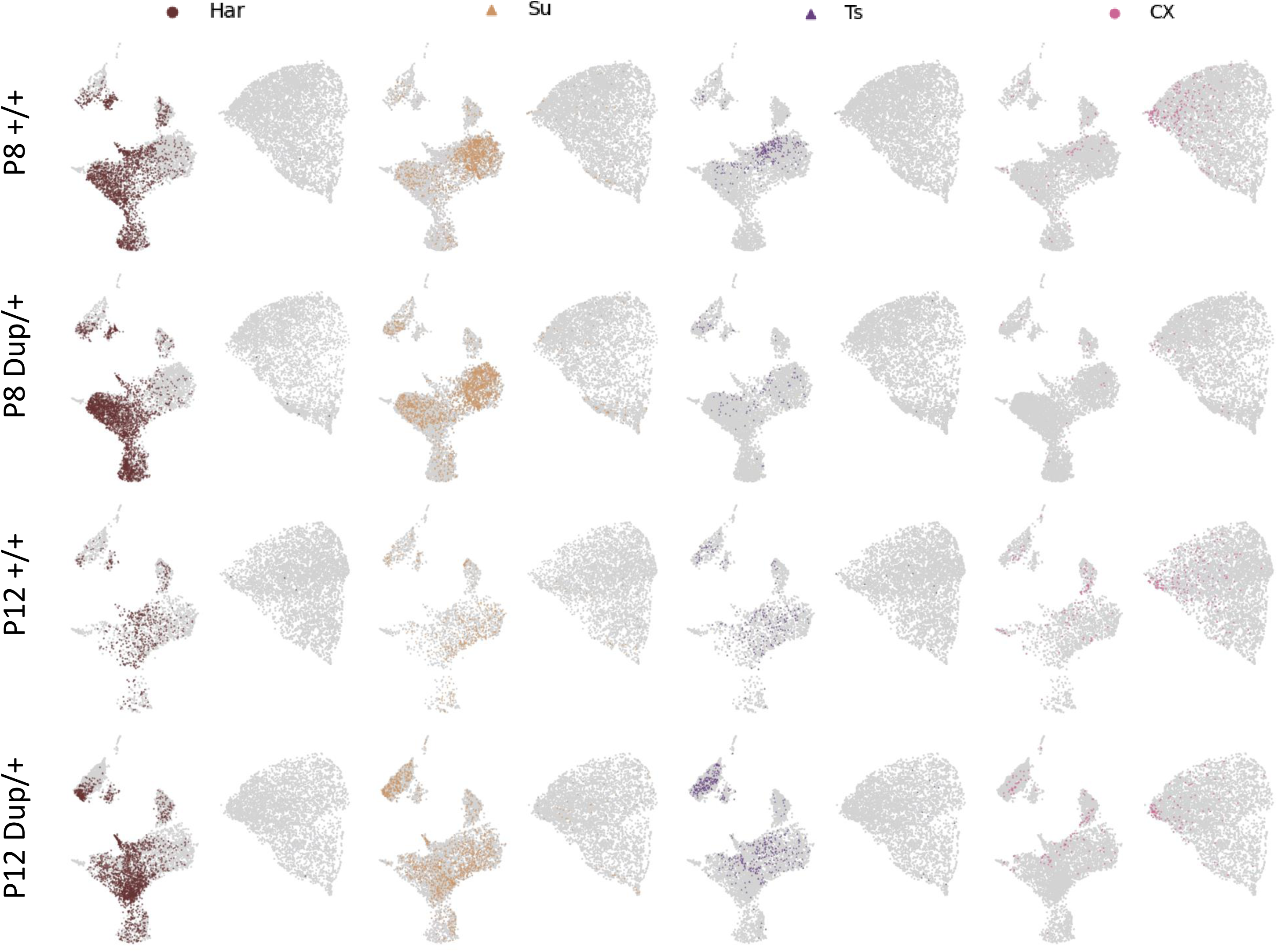
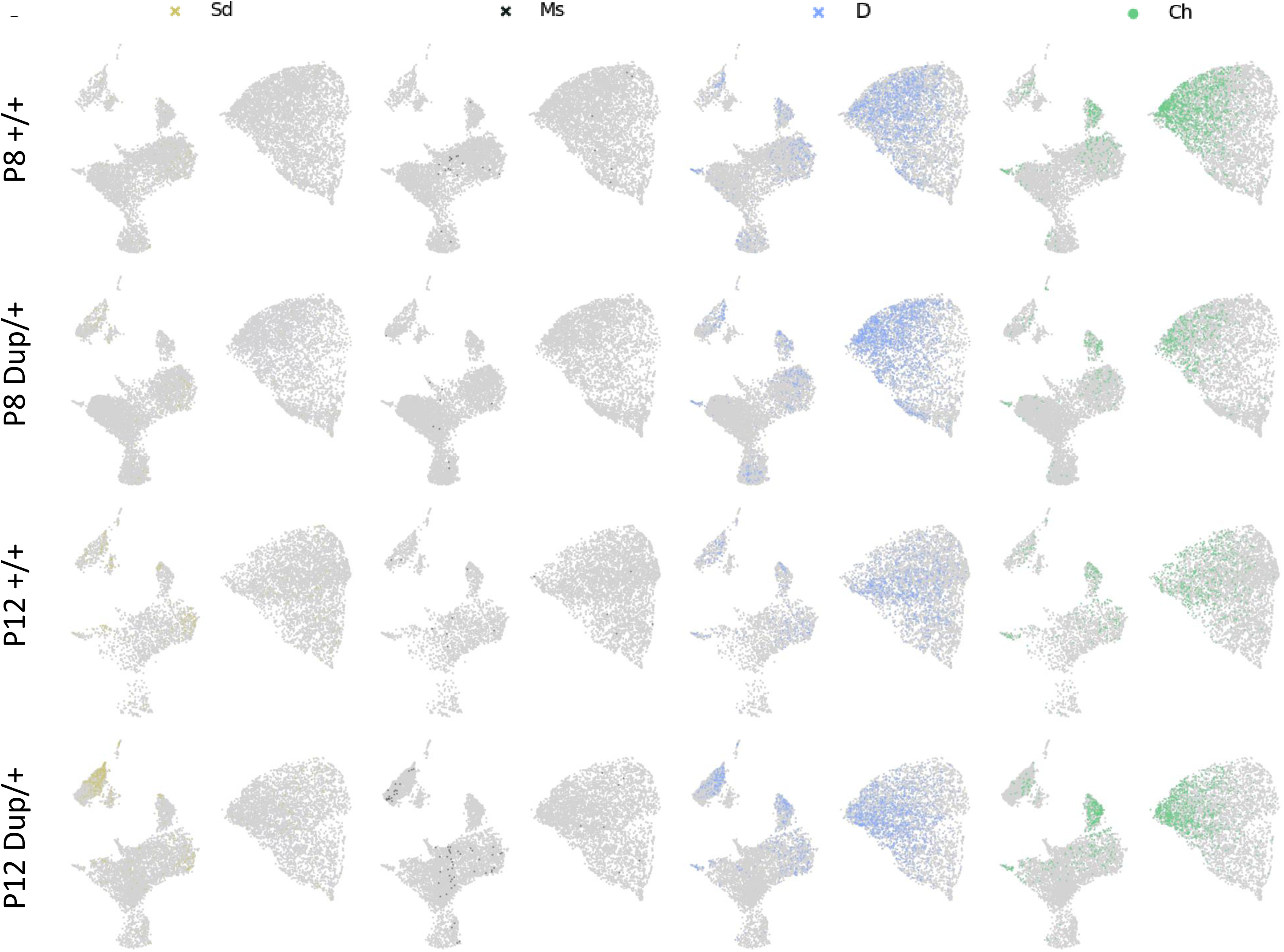
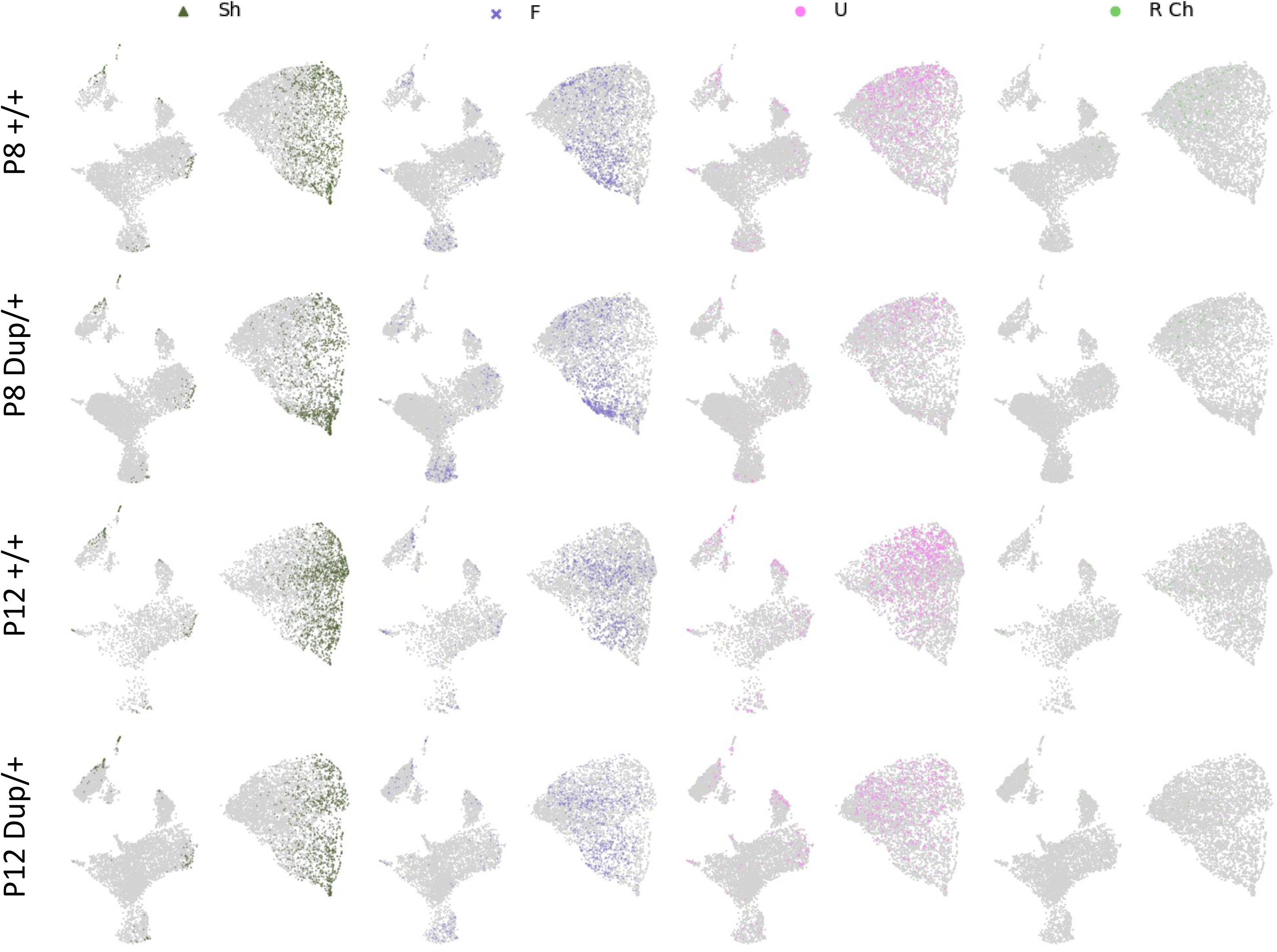
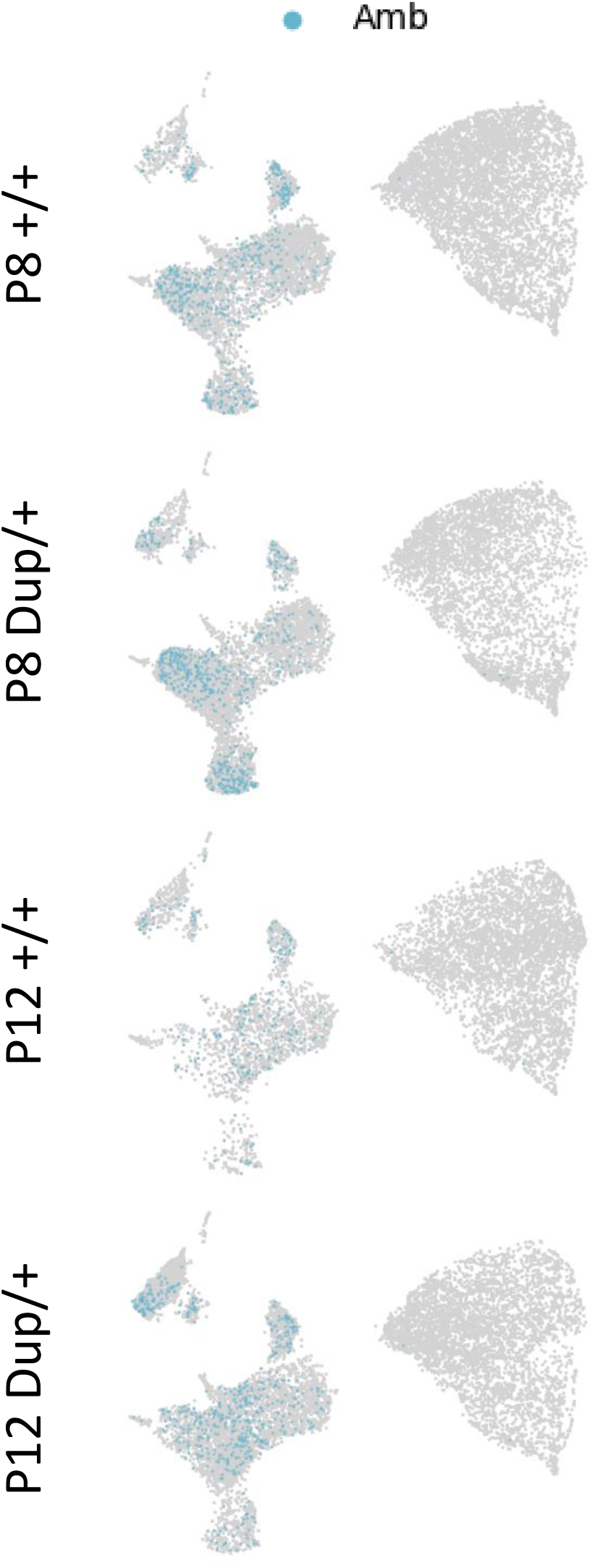
-1 to S3-4. The individual positions of each call type in two-dimensional UMAPs (uniform manifold approximation and projections). P, postnatal day. The distributions of each call type of +/+ at P8, Dup/+ at P8, +/+ at P12, and Dup/+ at P12 are plotted in the UMAP. Har, harmonic; Su, step-up; Ts, two-steps; Cx, complex; Sd, step-down; Ms, multiple-steps; D, downward; Ch, chevron; Sh, short; F, flat; U, upward; R Ch, reverse chevron; Amb, ambiguous.

**Supplementary Figure S4.**
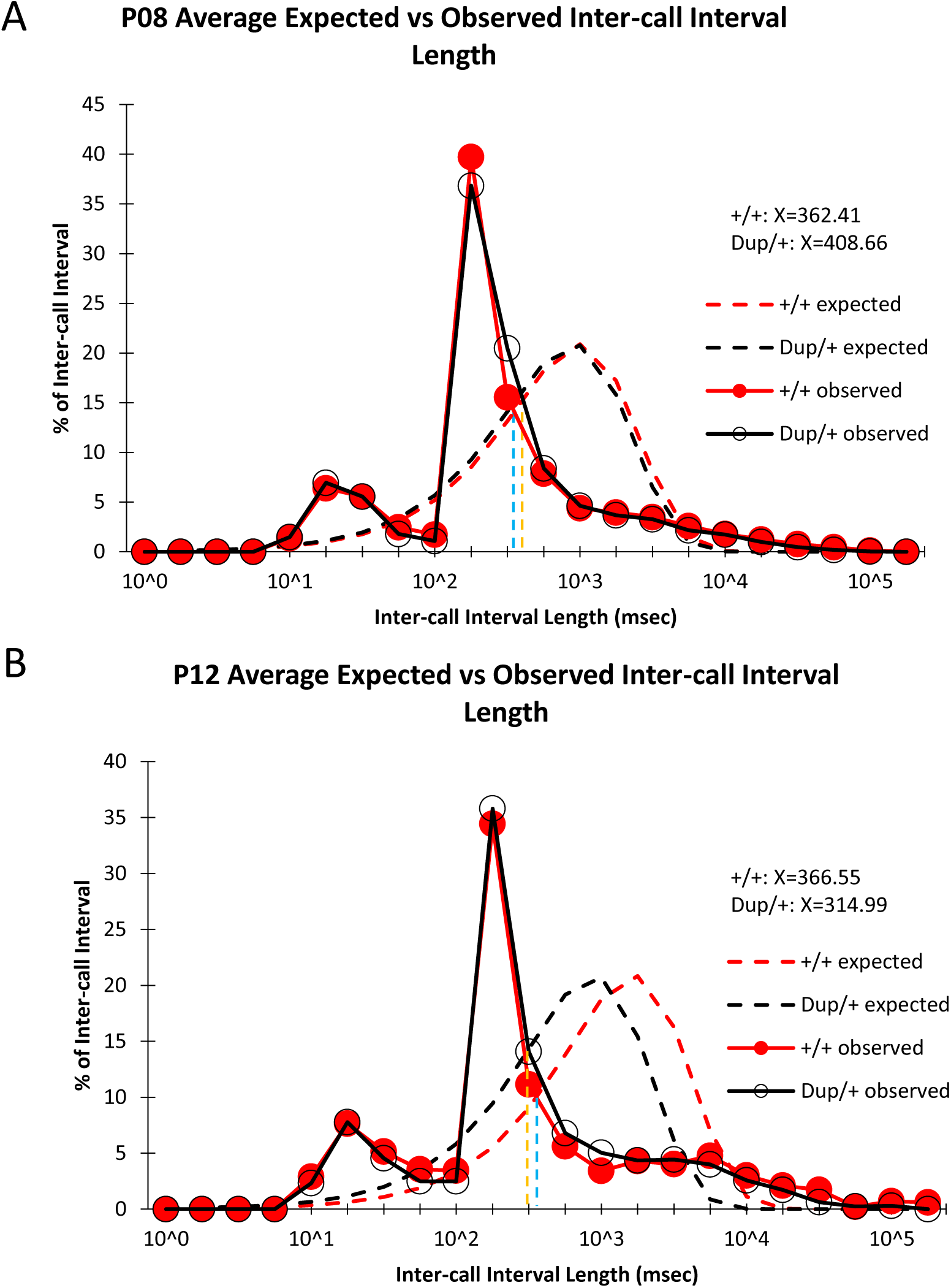
Percentages of inter-call intervals. The observed (black lines) and expected (red lines) inter-call intervals of Dup/+ and +/+ littermates are shown for P8 (**A**) and P12 (**B**). The observed distribution significantly shifted leftward from the theoretical distributions for P8 (+/+, D=0.4759, p<0.0001; Dup/+, D=0.4373, p<0.001) and P12 (+/+, D=0.5238, p<0.0001; Dup/+, D=0.4322, p<0.001), as determined by Kolmogorov–Smirnov tests. Blue and orange vertical dash lines indicate the cross points between the observed and expected inter-call intervals of +/+ and Dup/+, respectively.

**Supplementary Figure S5.**
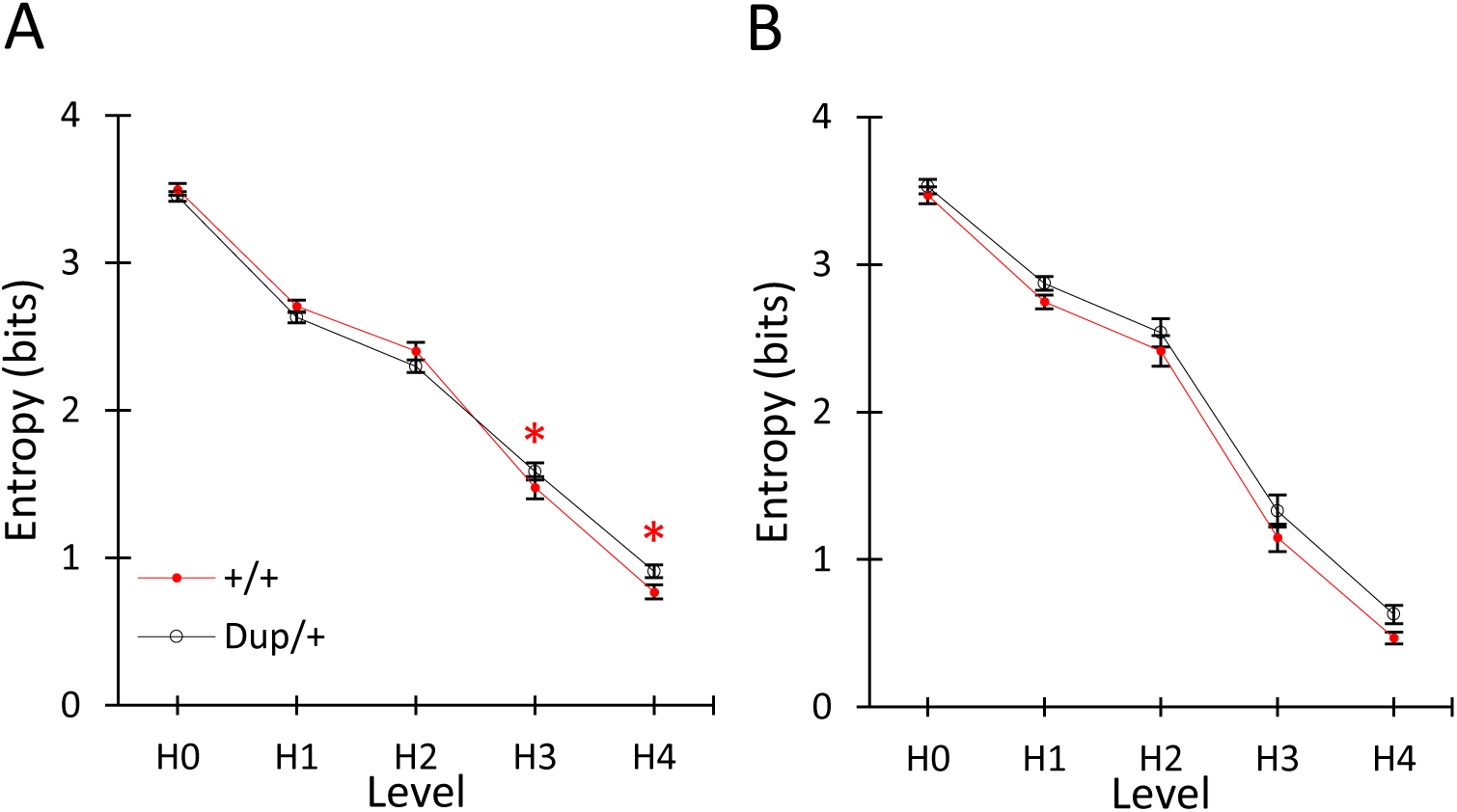
Shannon entropy analysis of the unpredictability of the selection of call types (H0), the choice of distinct calls within the chosen call types (H1), in two-call sequences (H2), three-call sequences (H3), and four-call sequences (H4) at P8 (**A**) and P12 (**B**). Data were analyzed using a linear mixed model fitted using REML (restricted maximum likelihood) with t-test degrees of freedom approximated using the Satterthwaite method. In both genotypes, the degree of randomness declined compared with H0 (P8, H1-H4, all for p<2.0 x10^-16^; P12, H1-H4, all for p<2.0 x10^-16^). +/+ mice and Dup/+ mice did not differ at any H. P, postnatal day. *, p<0.05.

**Supplementary Figure S6.**
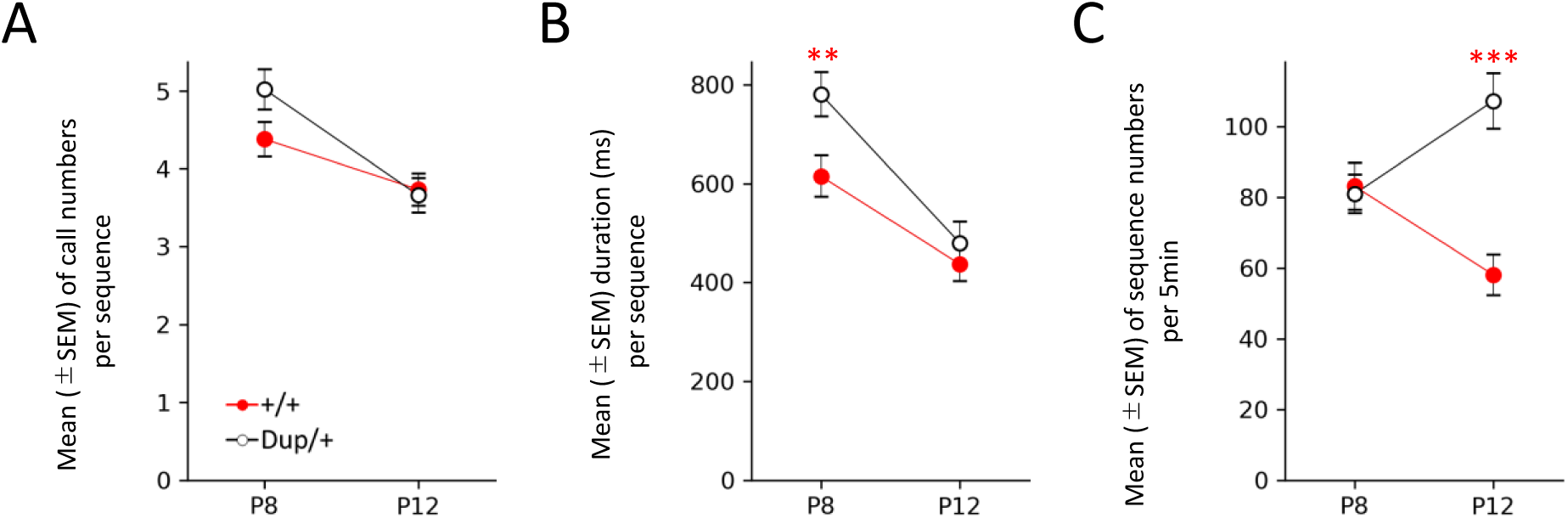
The mean (±SEM) numbers of calls per sequence (**A**), call sequence durations (**B**), and number of sequences emitted per 5 min (**C**). Dup/+ pups and +/+ pups did not differ in the call density (**A**, P8, t(52)=1.903, p=0.063; P12, t(51)=-0.242, p=0.8094), but Dup/+ pups has longer sequence duration than +/+ pups at P8 (**B**, P8, t(52)=2.707, p=0.0091; P12, t(51)=0.777, p=0.4409) and emitted more sequences per the 5 min test at P12 than +/+ pups (**C**. P8, U343.5, p=0.757; P12, U=138.5, p=1.00304E-05). P8: +/+, n=29; Dup/+, n=25. P12: +/+, n=28; Dup/+, n=25.

**Supplementary Figure S7.**
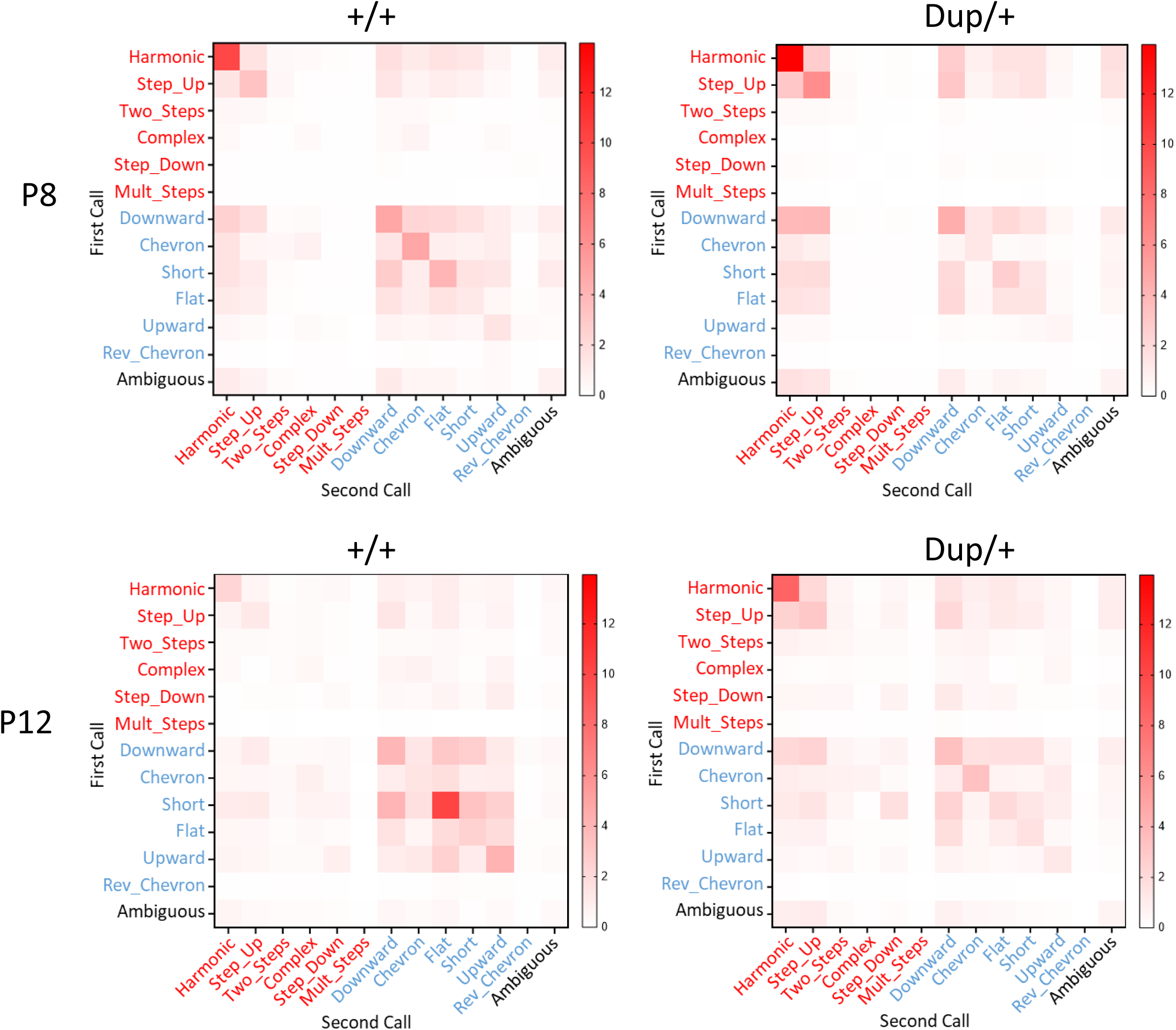
Call connections based on non-Markov proportions of two-call connections at P8 (top) and P12 (bottom) for +/+ (left) and Dup/+ (right) mice.

**Supplementary Figure S8.**
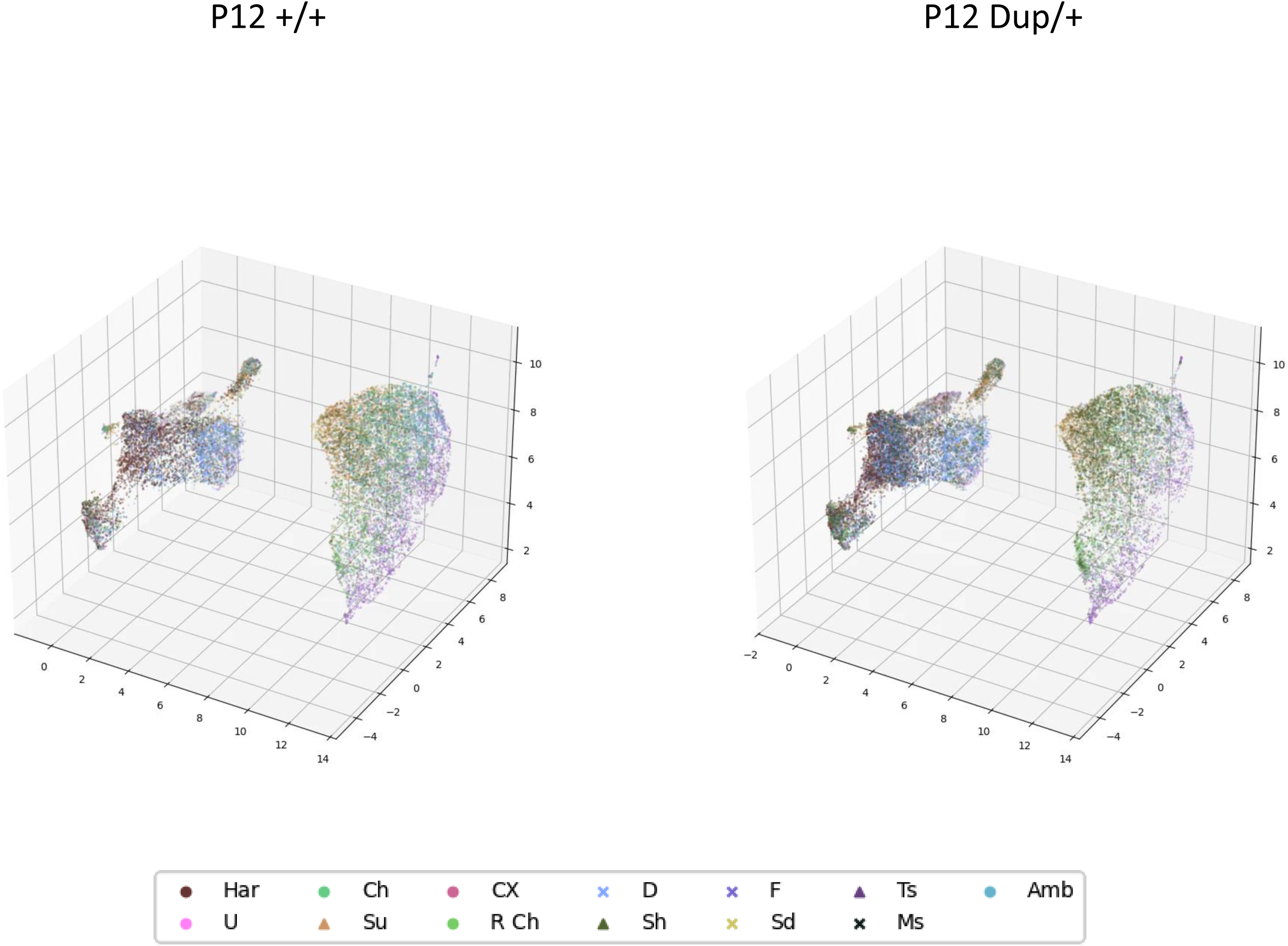
The representative temporal transitions of P12 calls of +/+ (top) and Dup/+ (bottom) mice in three-dimensional UMAPs.

**Supplementary Figure S9.**
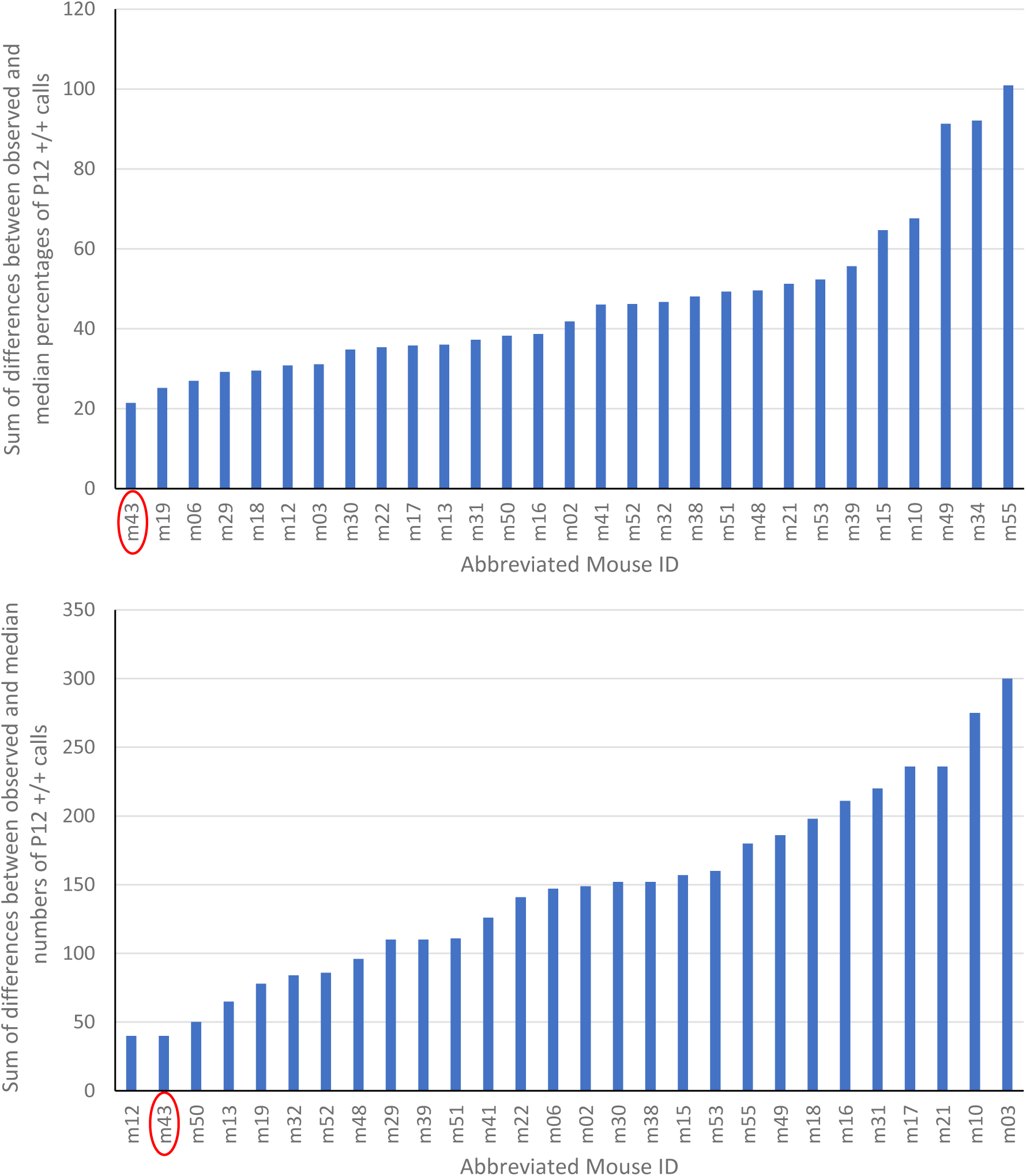
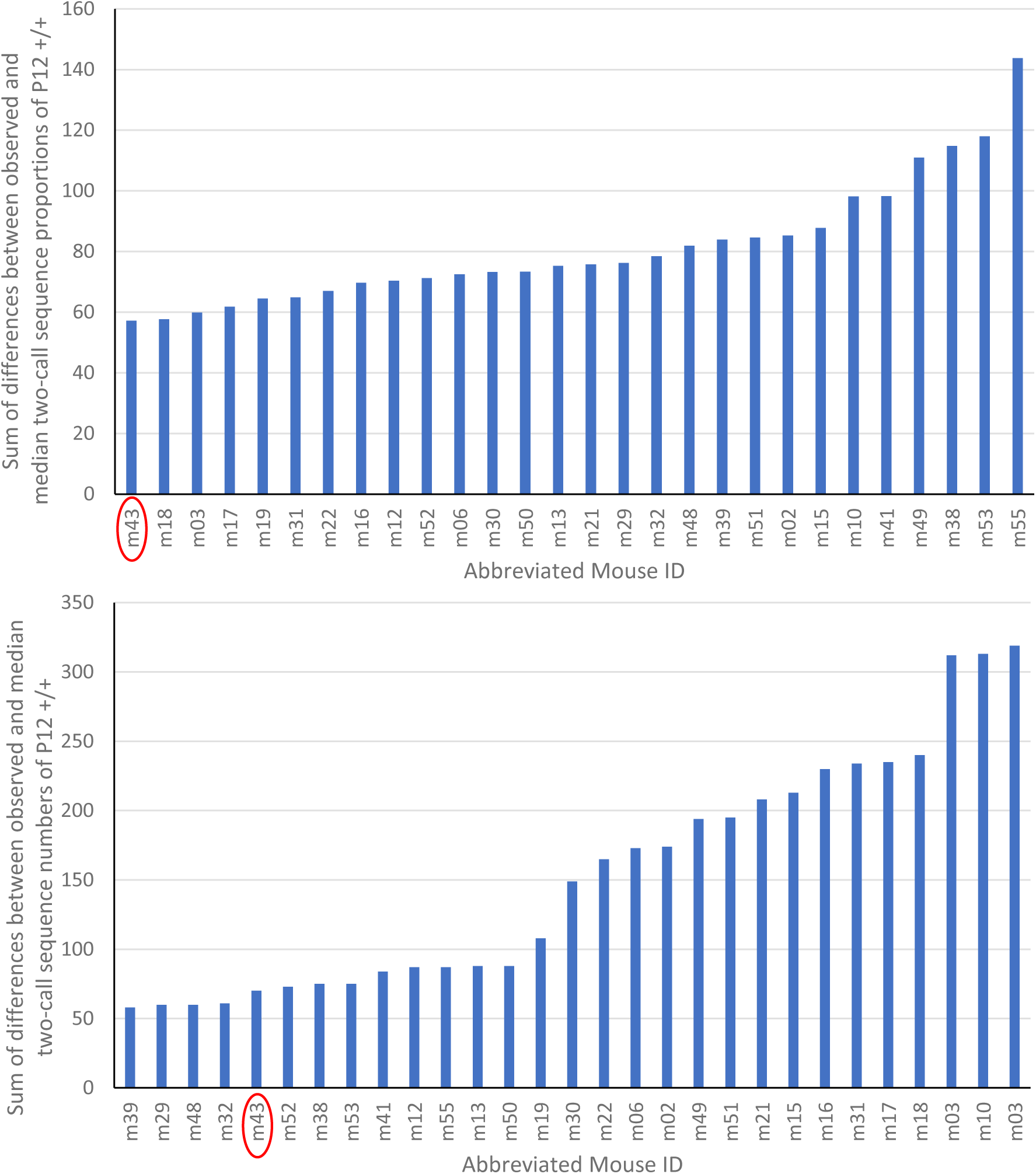
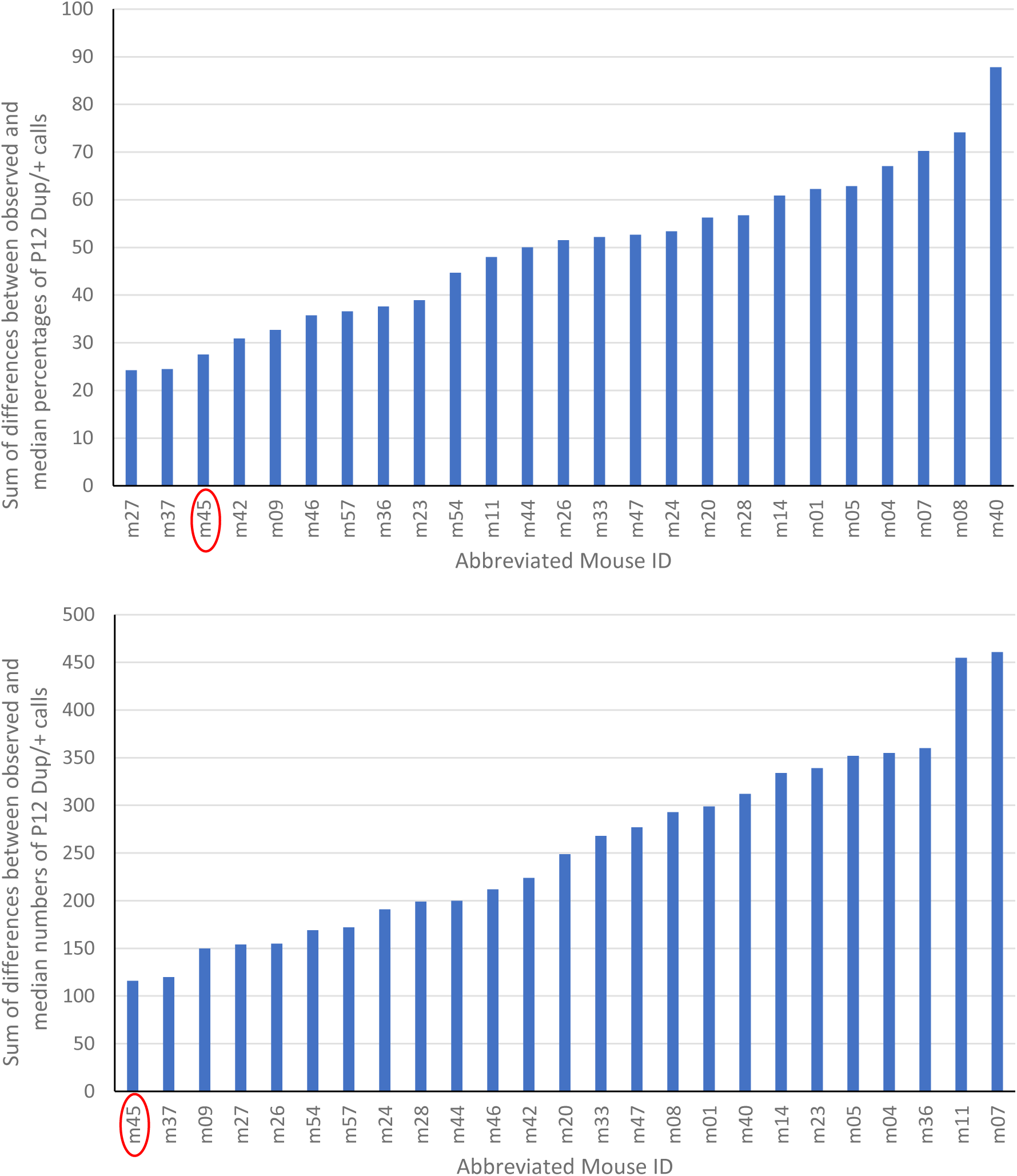
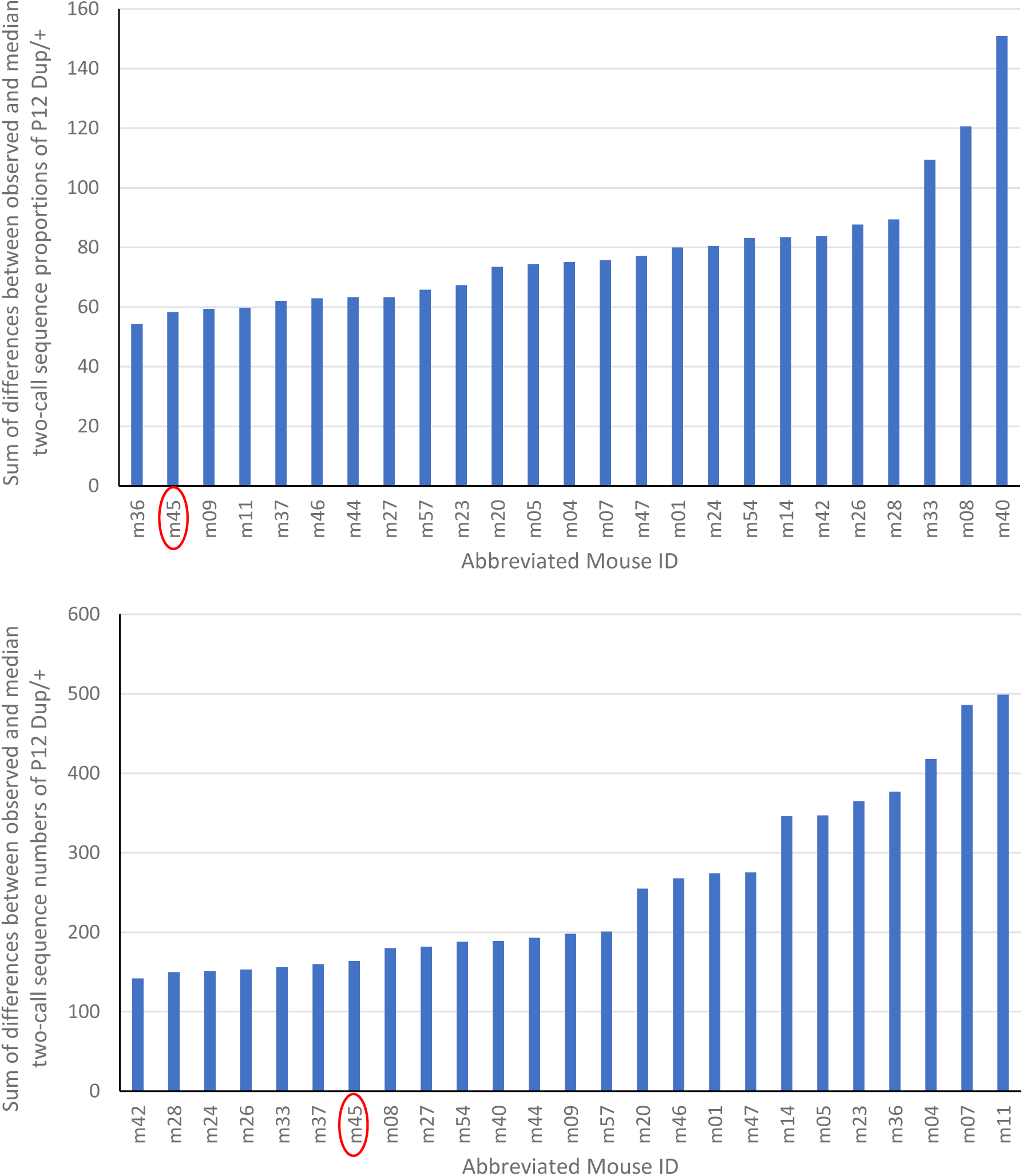
Proximity of calls to the group average. **S9-1**) Sum of differences between observed and median percentages (top) and between observed and median numbers (bottom) of P12 +/+ calls. **S9-2**) Sum of differences between observed and median two-call sequence proportions (top) and between observed and median numbers (bottom) of P12 +/+ calls. Calls of mouse ID m43 had the smallest difference from the group median in all measures. **S9-3**) Sum of differences between observed and median two-call sequence percentages (top) and between observed and median numbers (bottom) of P12 Dup/+ calls. **S9-4**) Sum of differences between observed and median two-call sequence proportions (top) and between observed and median numbers (bottom) of P12 Dup/+ calls. Calls of mouse ID m45 had the smallest difference from the group median in all measures.

**Supplementary Figure S10.**
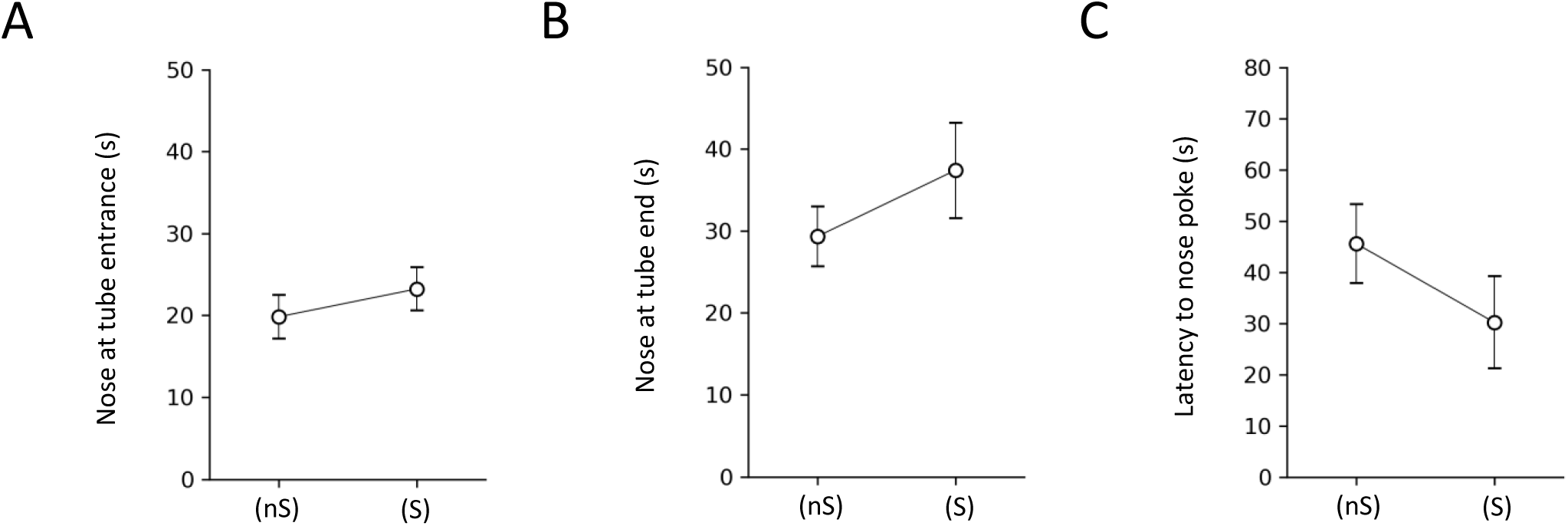
Maternal approach in response to the two tubes ((S) and (nS)) in which no sound was played back as a negative control to determine the baseline preference for the two tubes. We analyzed time the mother’s nose peeking at the two entrances (**A**) and ends (**B**) of the tubes, and (**C**) latency to enter the entrances of the two tubes. Mothers spent indistinguishable amounts of time peeking at the two tubes (**A**, p=0.4254), at the ends of the two tubes (**B**, p=0.3529), or latency to the first nose poke into the entrances of the two tubes (**C**, Dup/+ calls, p=0.051). N = 17.

**Supplementary Figure S11.**
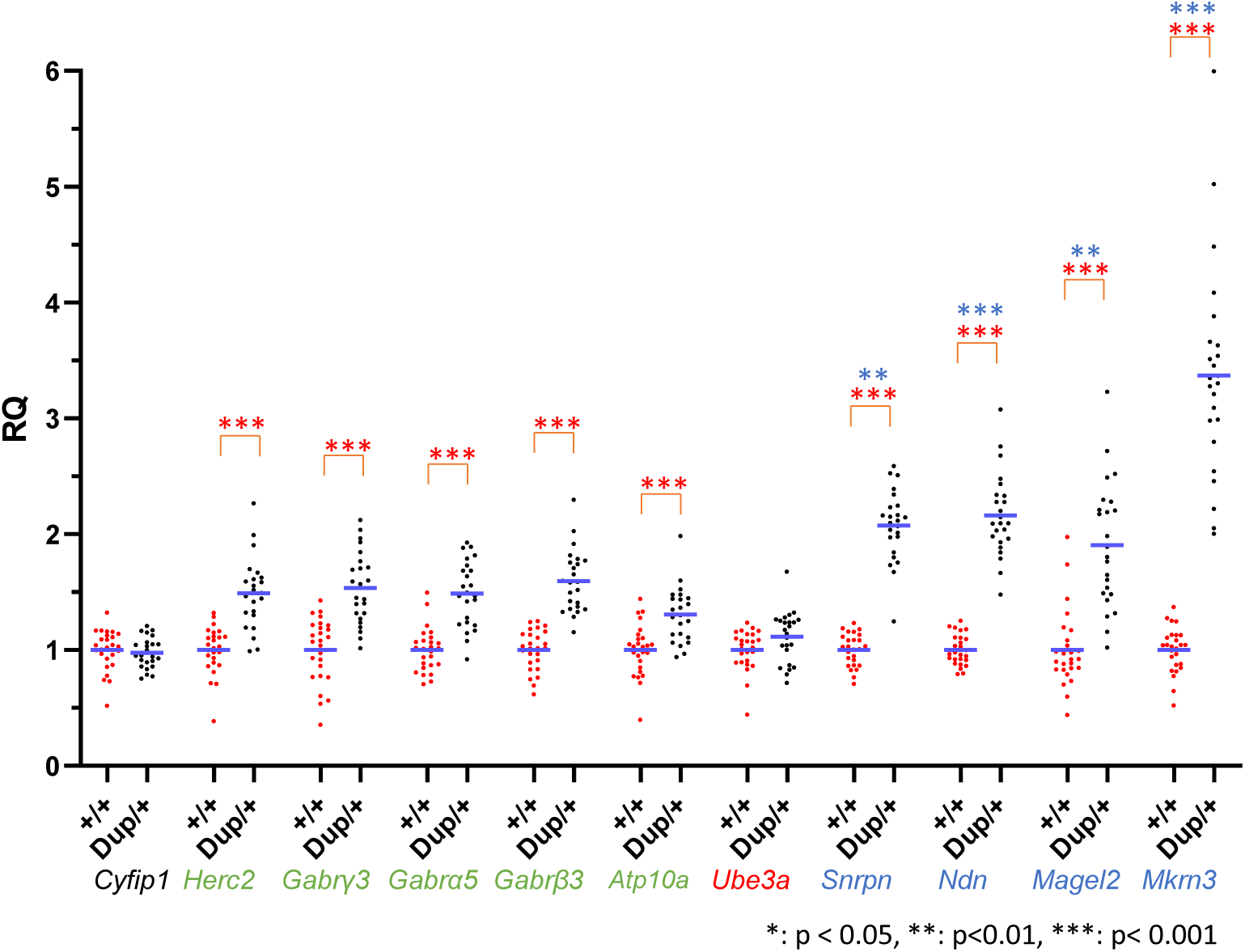
The relative quantification (RQ) (mean+SEM) of gene expression levels of the rest of the forebrain of individual Dup/+ (black dots) mice relative to +/+ (red dots) mice, as determined by qRT-PCR. Statistically significant differences between +/+ and Dup/+ data are shown as *, p<0.05; **, p<0.01; ***, p<0.001, as determined by Mann-Whitney nonparametric tests; the original p values of those data that remained statistically significant after Benjamini–Hochberg correction at the false detection rate of 5% are shown. Gene expression levels were statistically significantly higher in Dup/+ than in +/+ mice for all duplicated genes examined (*, p<0.05; ***, p<0.001). *Cyfip1*, a control gene outside the duplicated chromosomal segment, did not differ between Dup/+ mice and +/+ mice (p=0.4259). The variance of gene expression was significantly larger in Dup/+ than in +/+ mice for *Mkrn3* (***, p=0.0003), *Ndn* (***, p=0.0007), *Snrpn* (**, p=0.0089), *Magel2* (**, p=0.0093), and *Gabβ3* (*, p<0.05). +/+, n=26, Dup/+, n=23. Some forebrain tissues (+/+) were of poor quality and thus were not used for analysis (see **Table S11**, **Column C**, **# of samples**).

**Supplementary Figure S12.**
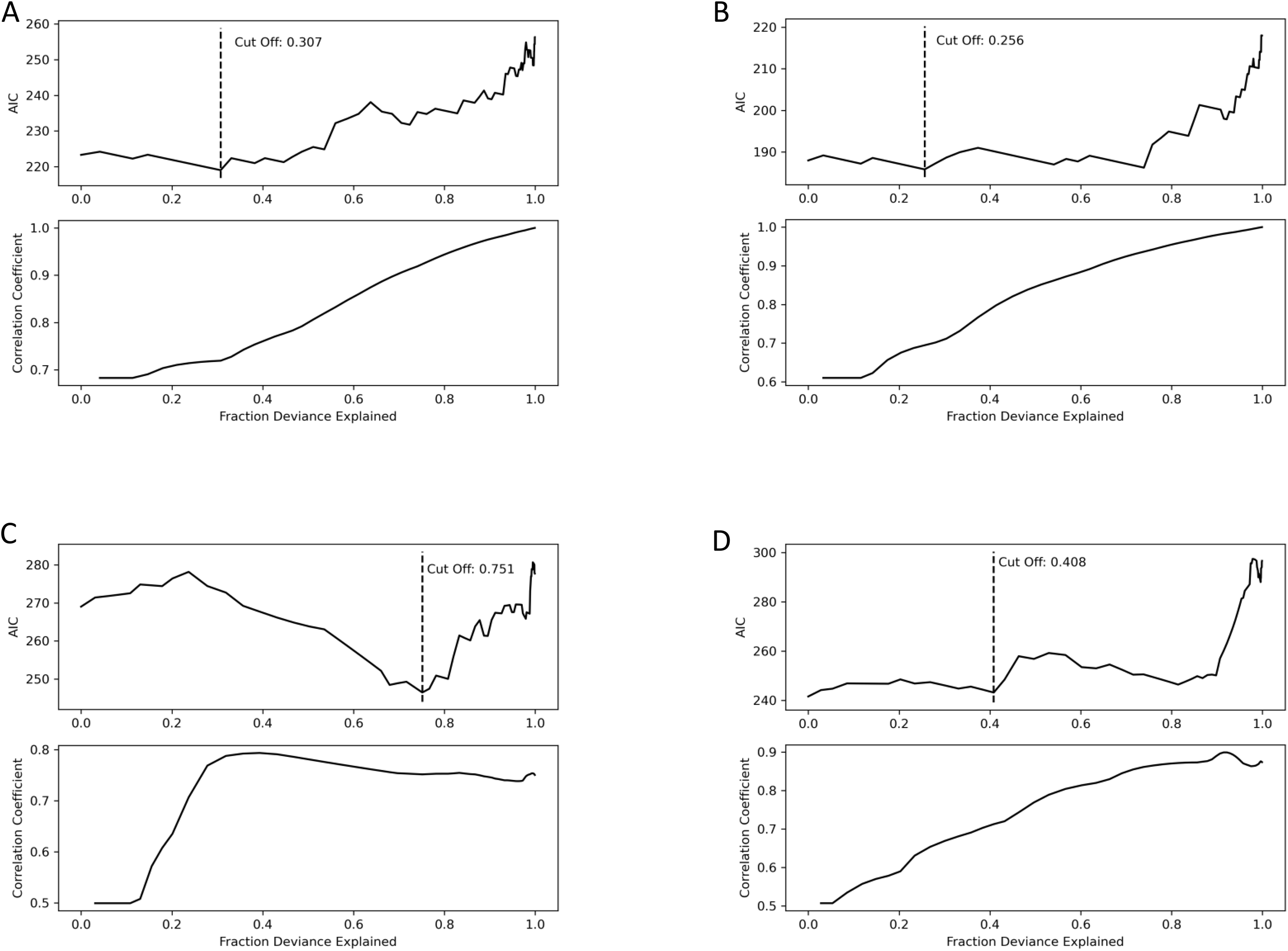
The Akaike Information Criterion determined the number of parameters needed for the best balance between goodness of fitness and model (see Figure 4). The smallest AIC value indicates the best tradeoff point and its corresponding value of Fraction Deviation Explained is shown as “Cut Off” (**A**, 0.307, Social Interaction Session 1, +/+; **B**, 0.256, Social Interaction Session 1, Dup/+; **C**, 0.751, Social Interaction Session 2, +/+; **D**, 0.408, Social Interaction Session 2, Dup/+). At each cut-off, high correlation coefficients (∼0.7) are achieved.

**Supplementary Figure S13.**
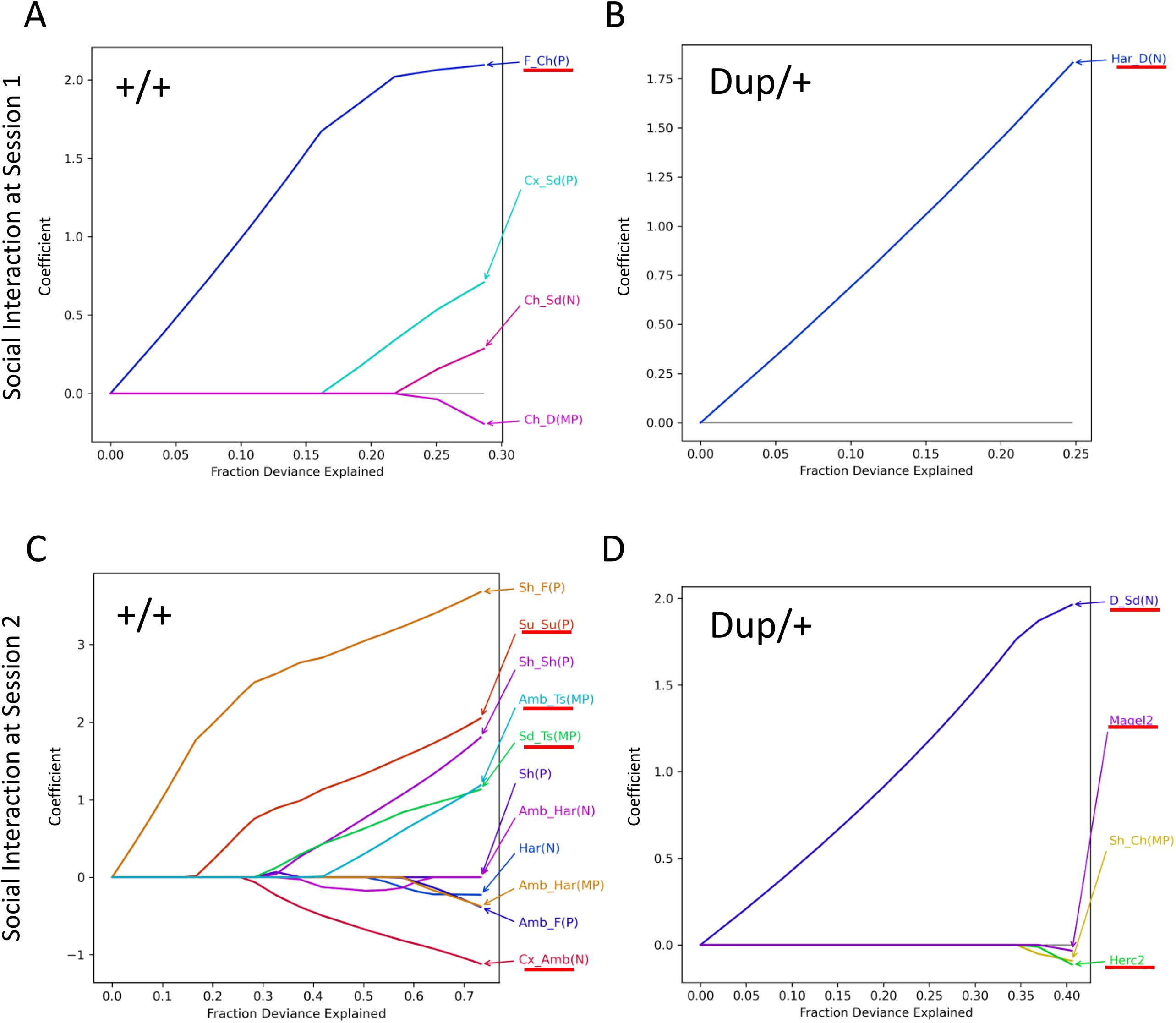
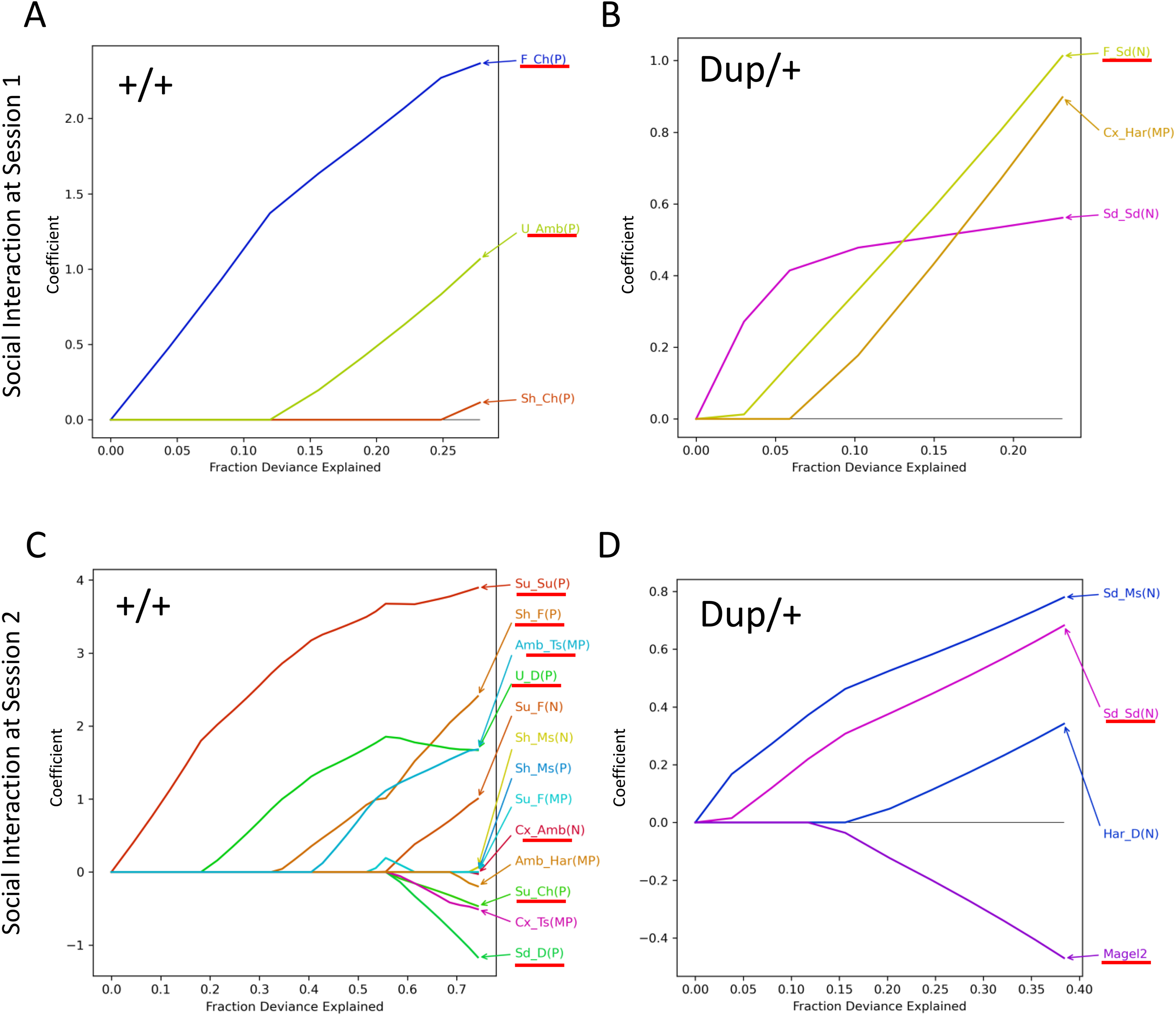
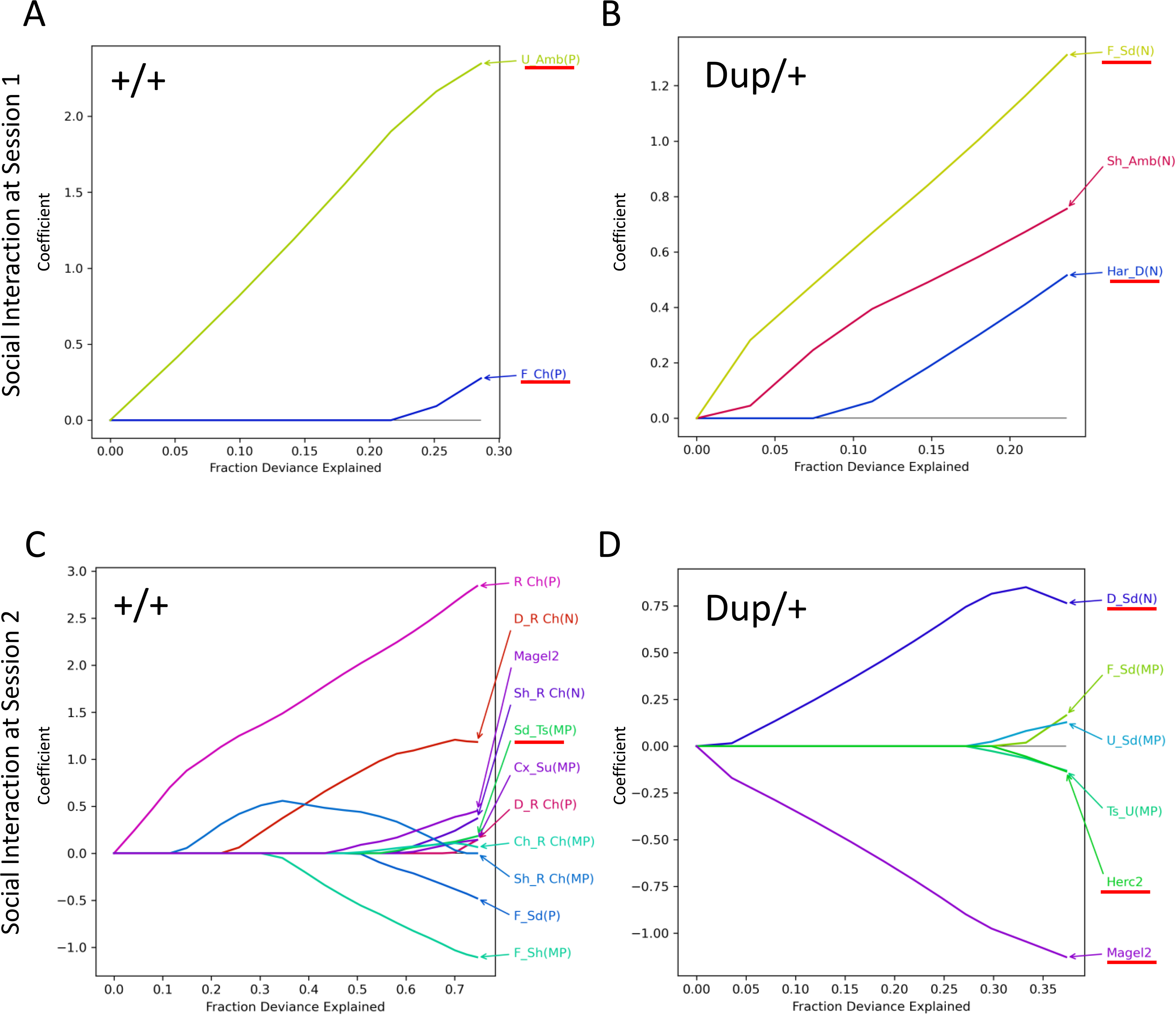
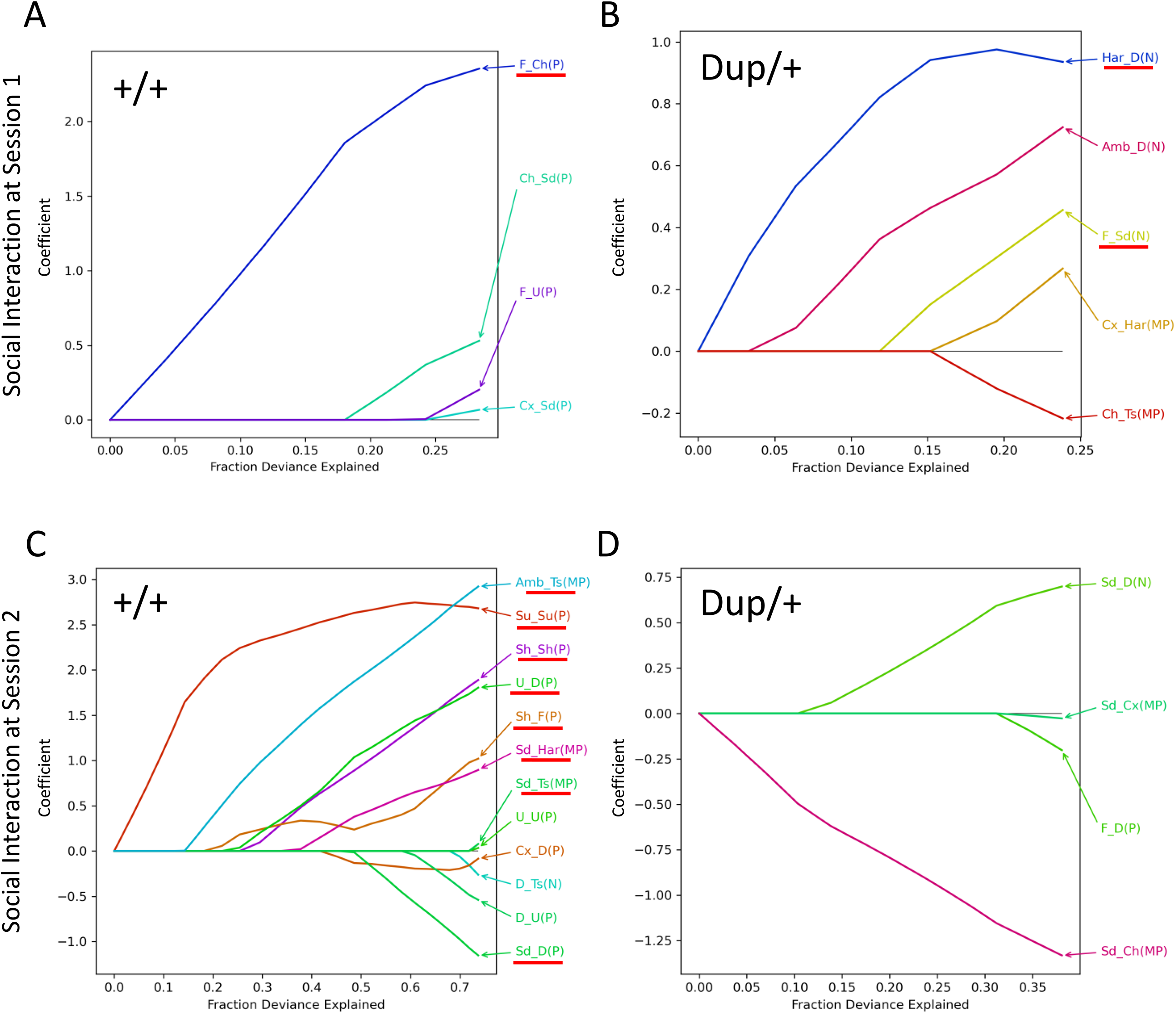
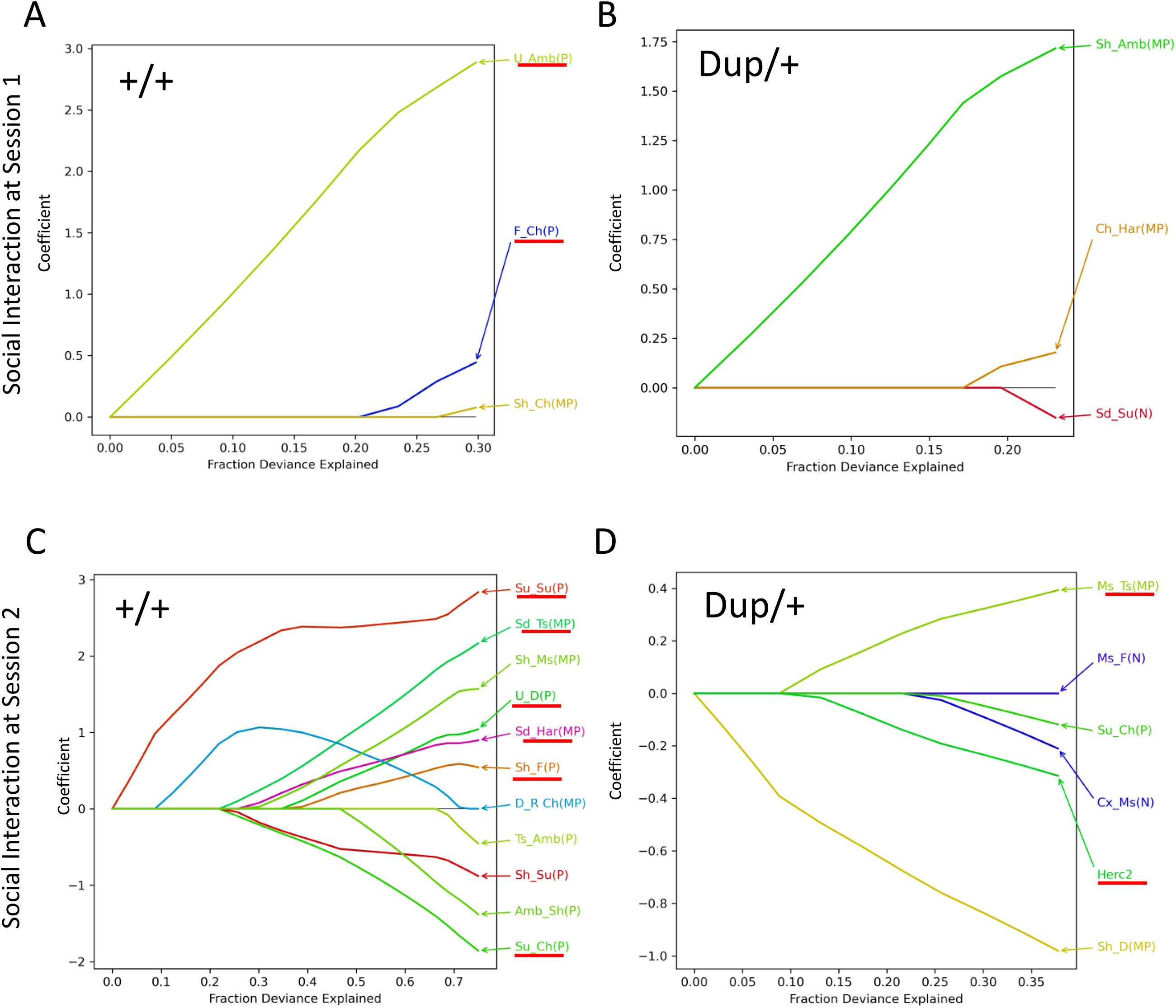
Fraction deviance explained, and coefficients of variables based on 5 Lasso models (**Fold 1** to **5**) of samples from which 5 or 6 randomly chosen mice were excluded from analysis. Social interaction session 1 +/+ (**A**) and Dup/+ (**B**). Social interaction session 2 of +/+ (**C**) and Dup/+ (**D**). The colors of lines are arbitrarily assigned. The number (N) and proportions (P) of each call type, number (N) and probabilities (P) of two-call sequences, Markov probabilities (MP) of two-call sequences, and gene expression scores of the prefrontal cortex were pooled and used for selection. The dependent variable was affiliative social interaction scores. Variables identified in the original Lasso models (see Figure 4) are underlined in red each validation model. Har, harmonic; Su, step-up; Ts, two-steps; Cx, complex; Sd, step-down; Ms, multiple-steps; D, downward; Ch, chevron; Sh, short; F, flat; U, upward; R Ch, reverse chevron; Amb, ambiguous.

**Supplementary Figure S14.**
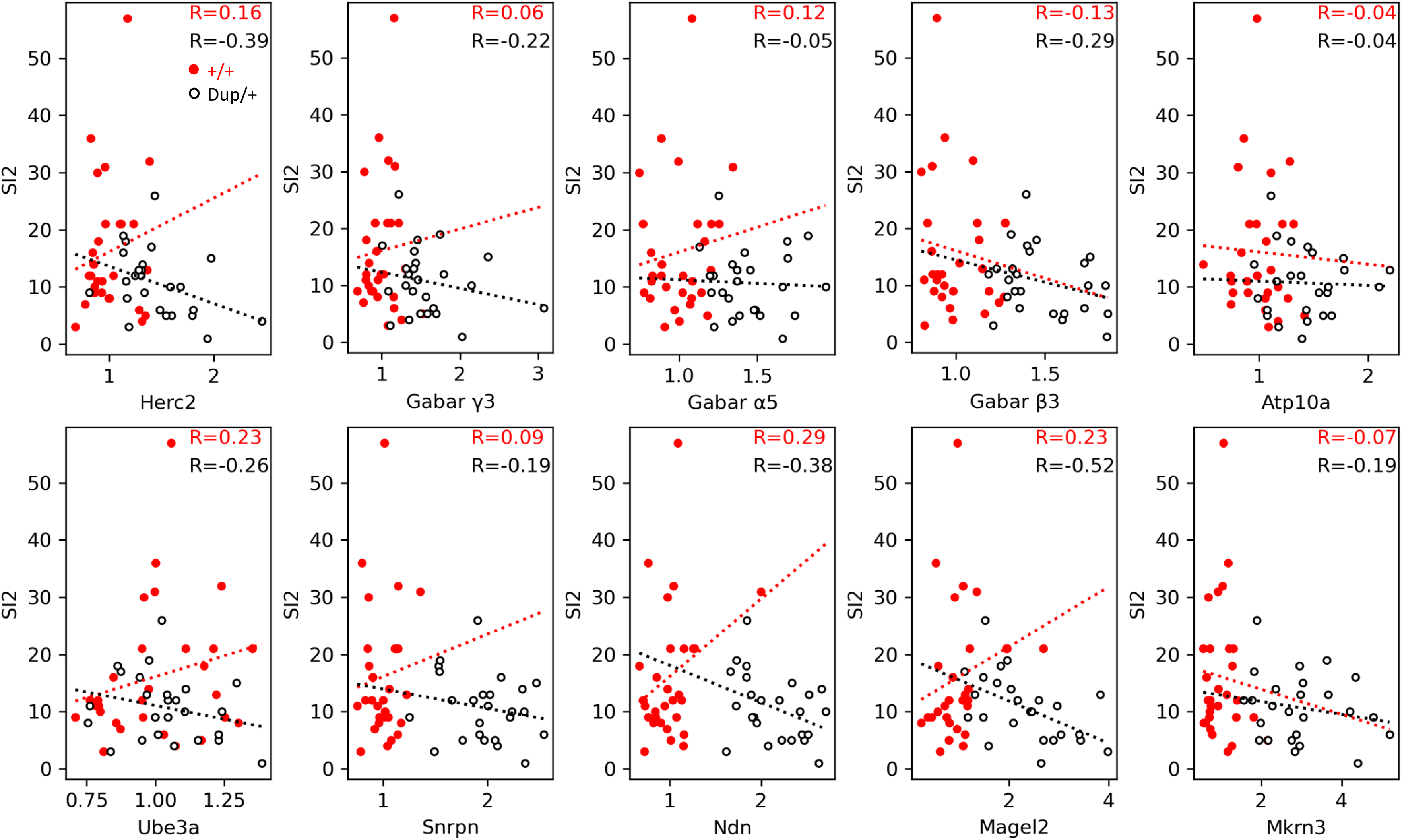
Correlation plots of each mouse based on their gene expression (ratio to the average of +/+) in the prefrontal cortex and social interaction scores at session 2 (SI2 of Y axis) of +/+ (red dots) and Dup/+ (white dots with black lines) mice. Correlation coefficients (R) were separately computed for each genotype. P values that remained significant after correction for multiple comparisons at 5, 10 and 25% FDR are indicated in **Table S2-Figure S14**.

**Table S1.**
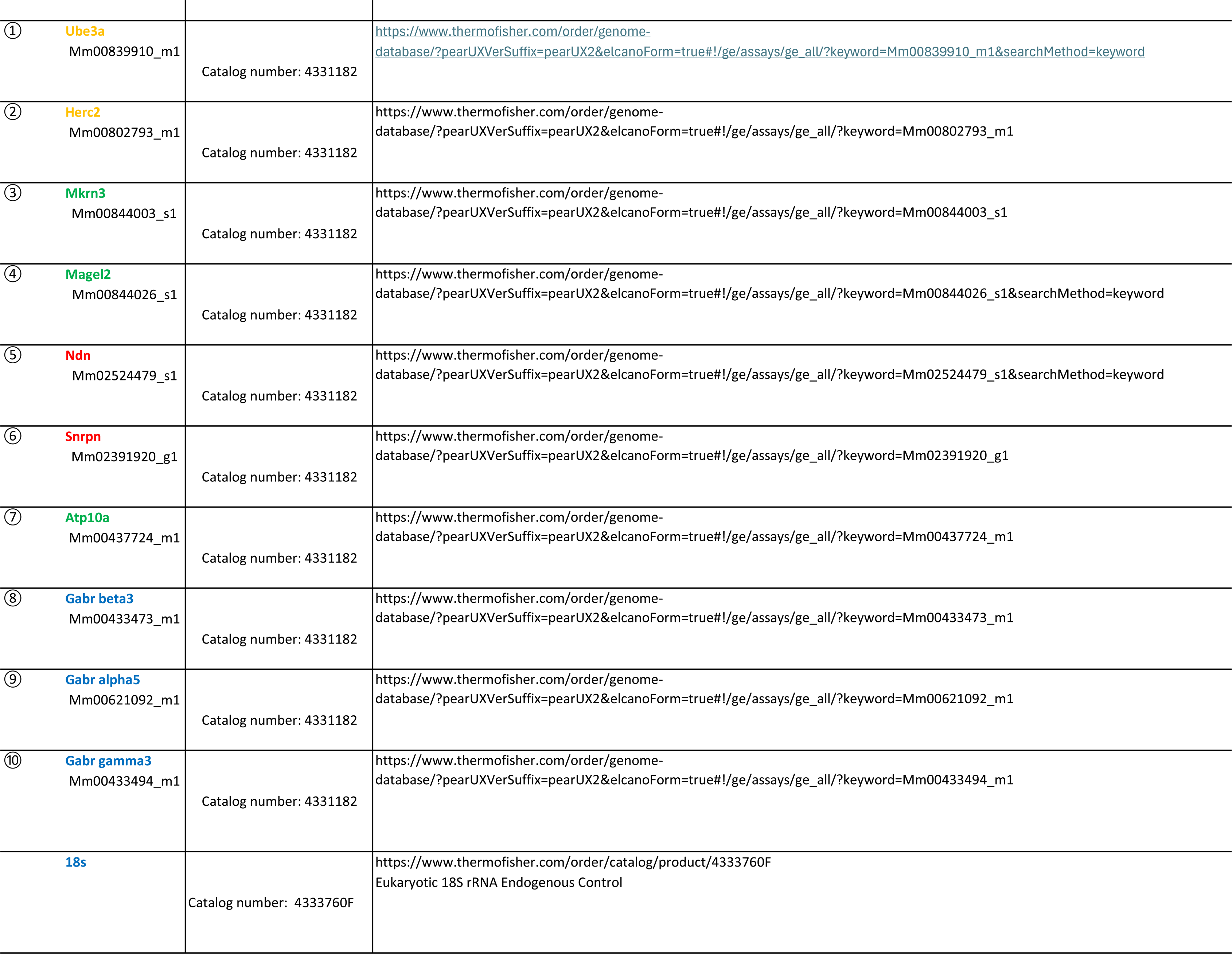

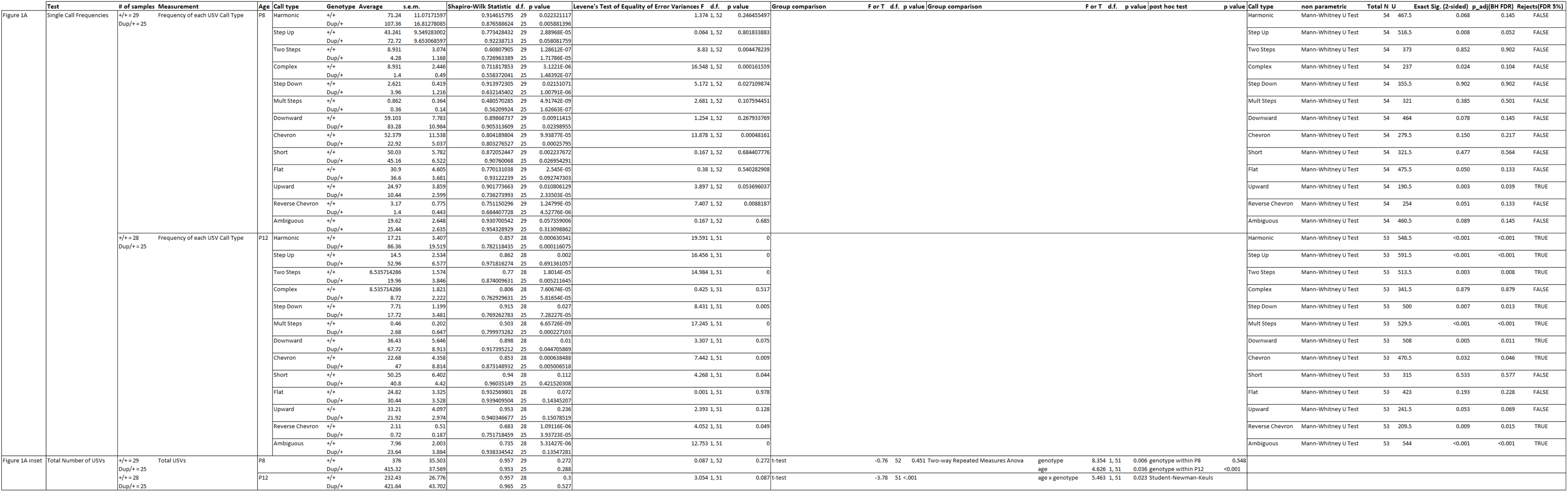
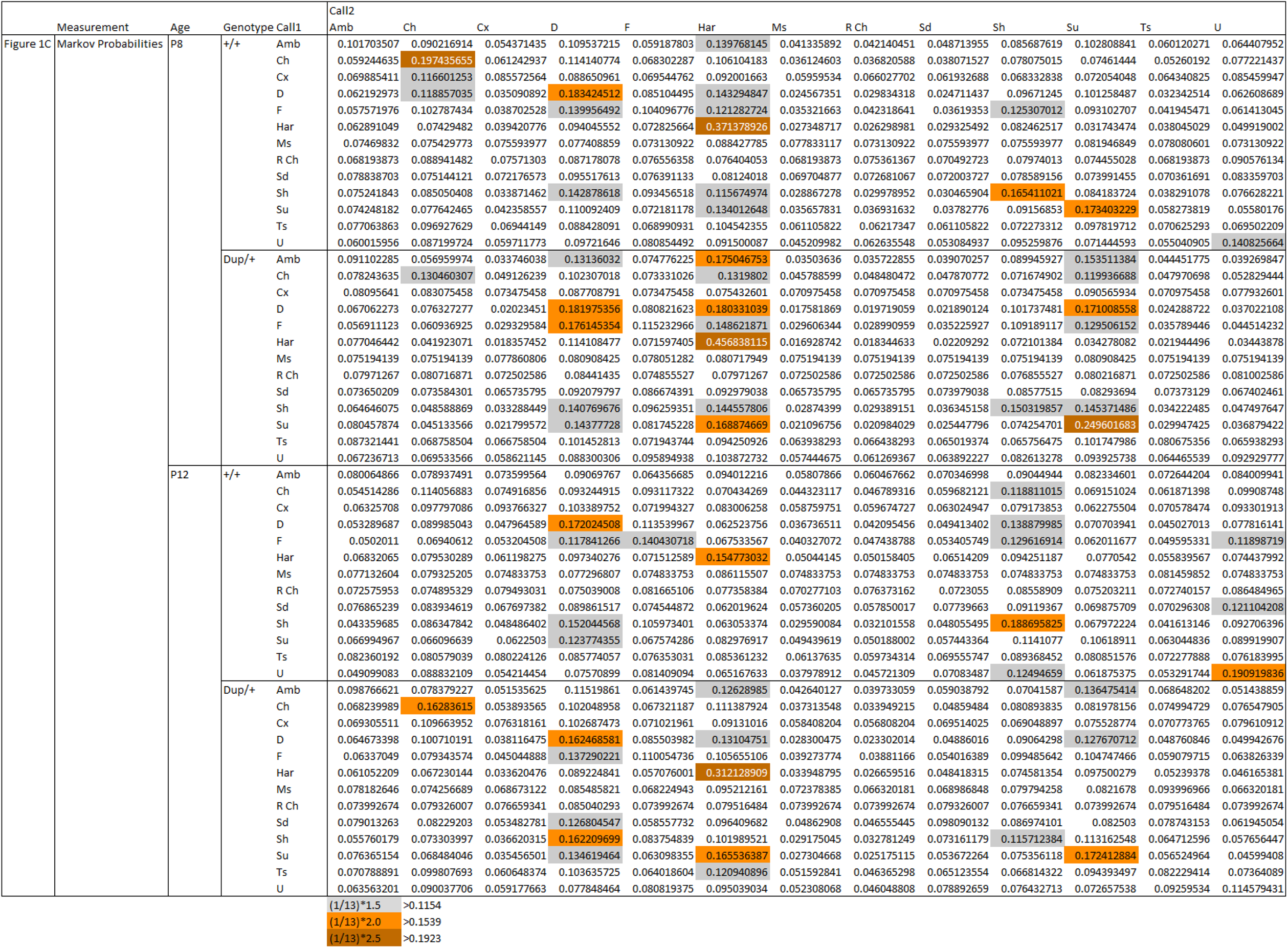
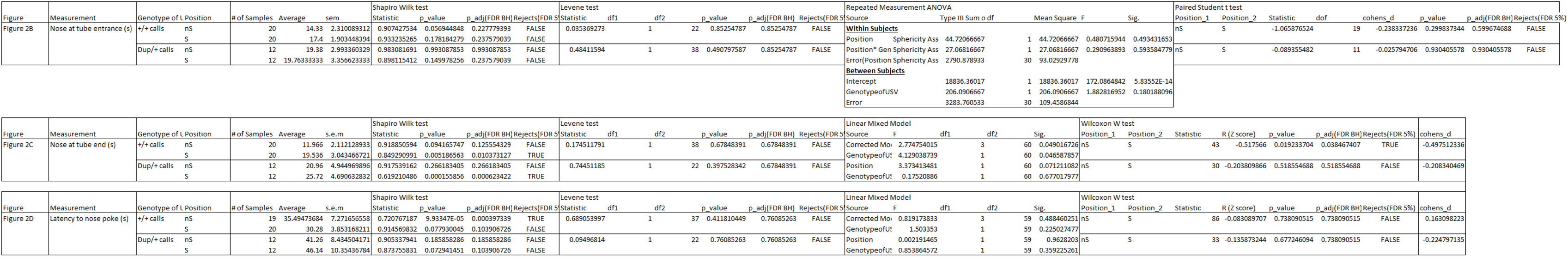
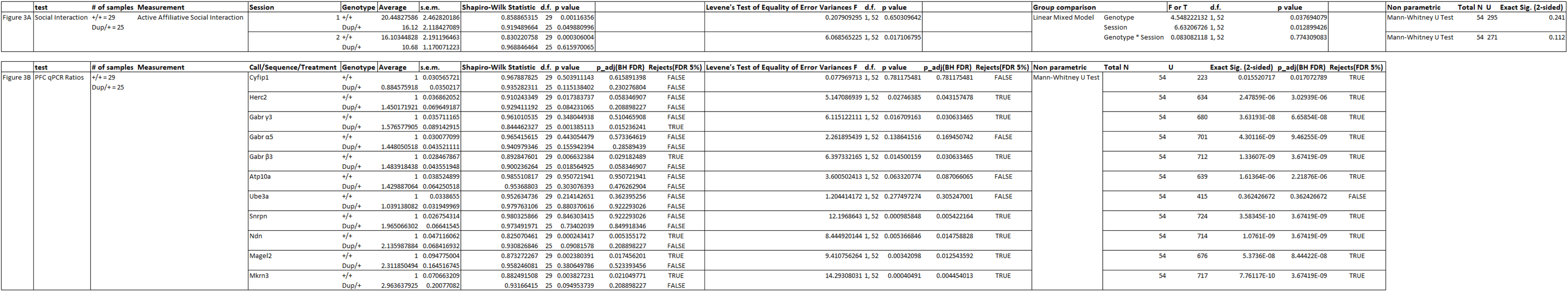
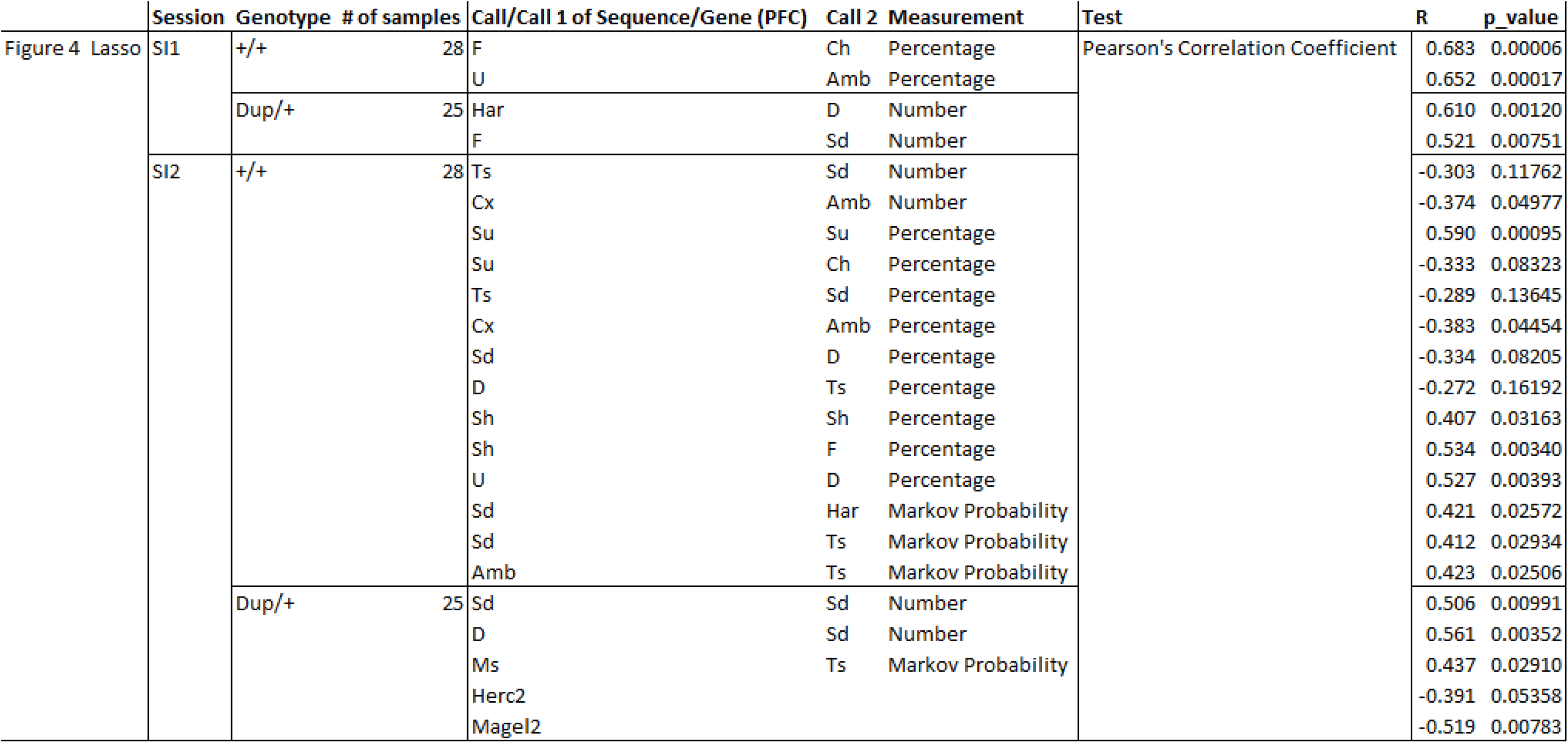
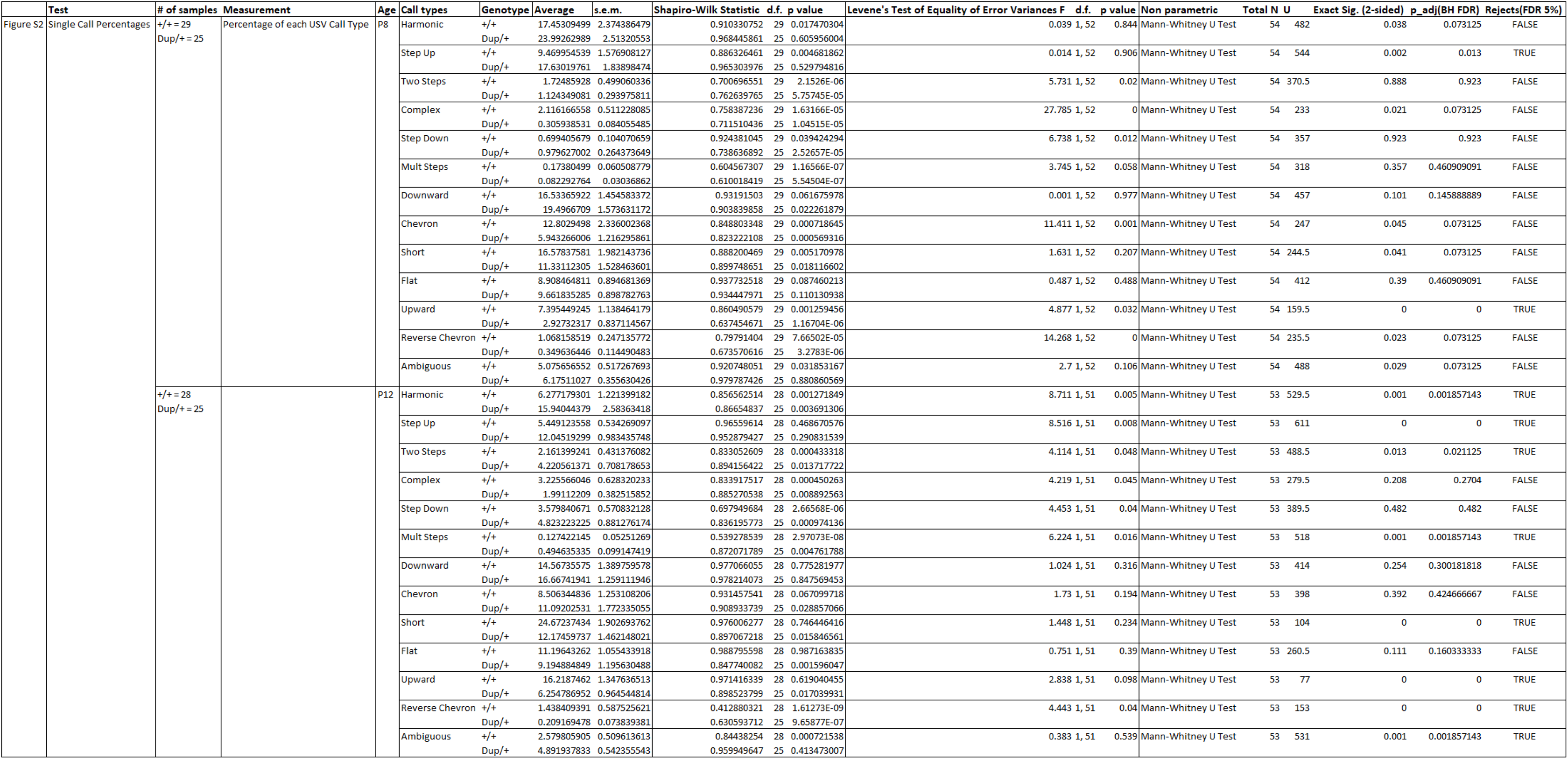
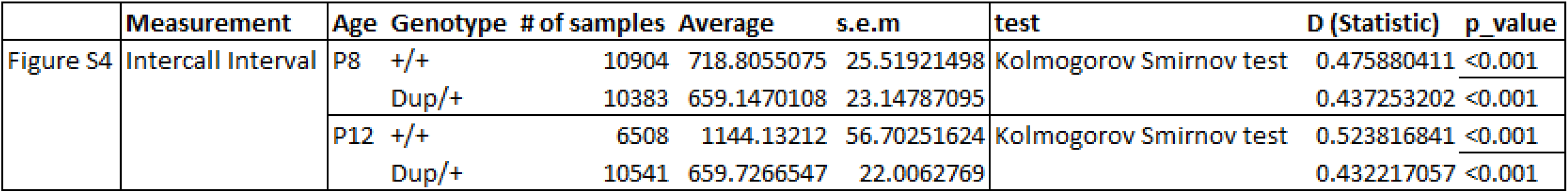
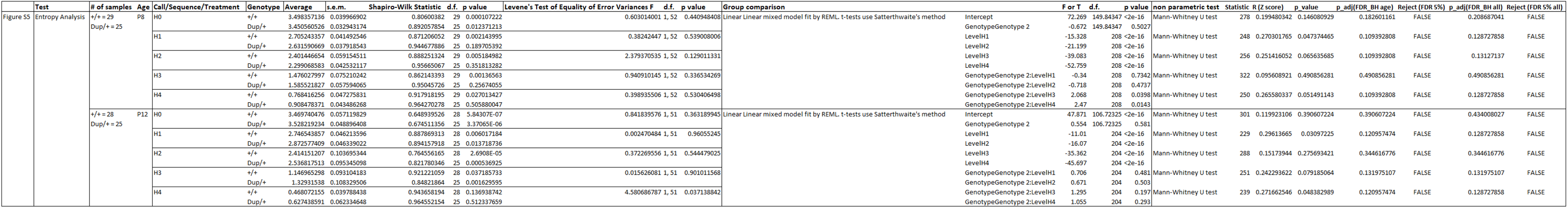
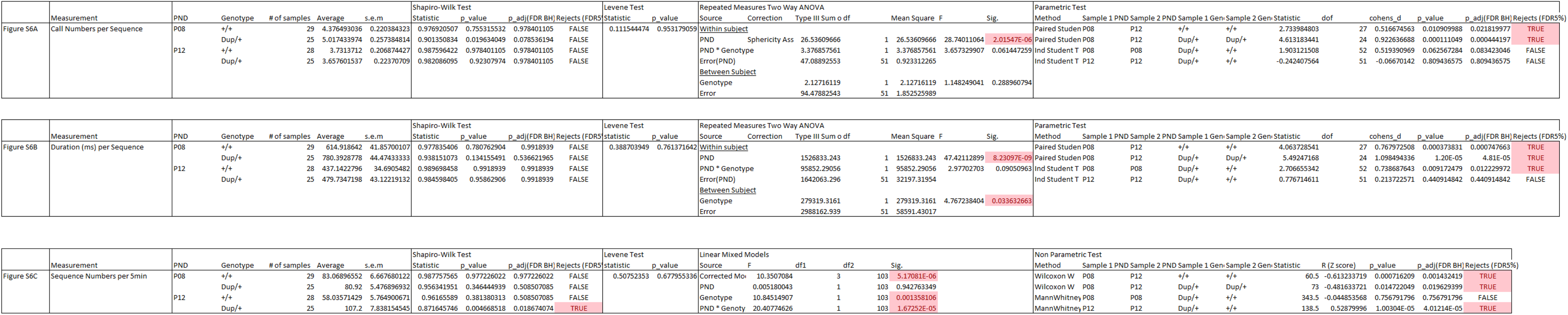
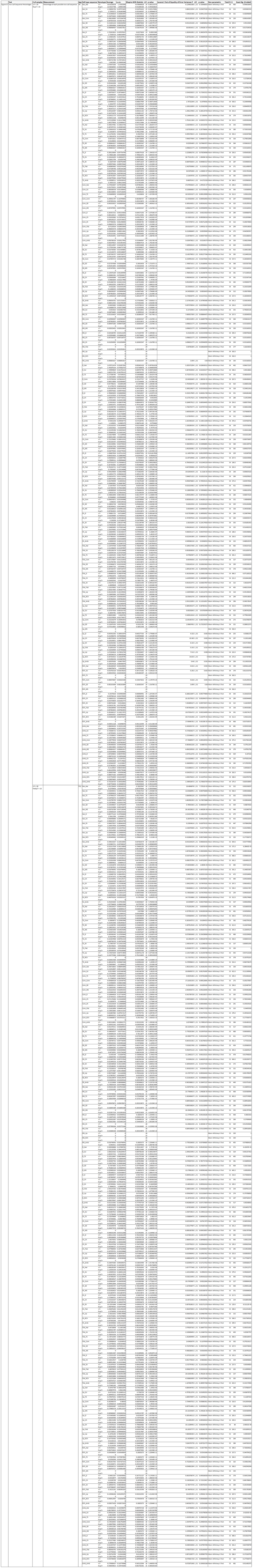
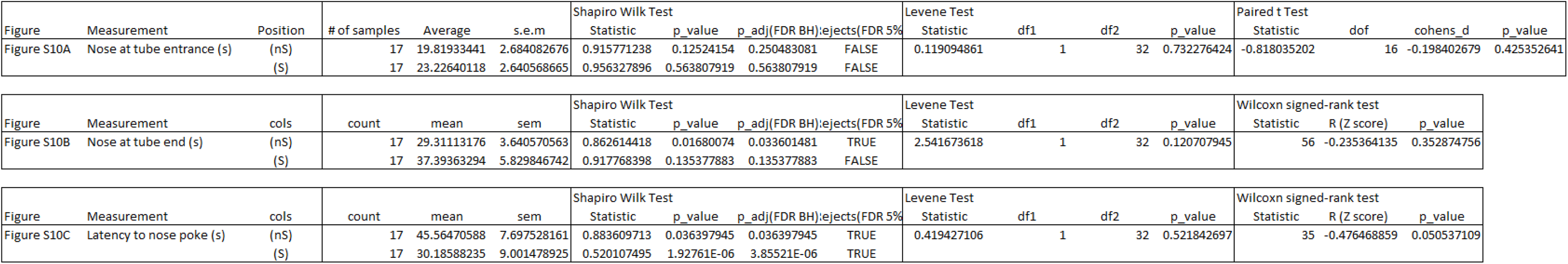
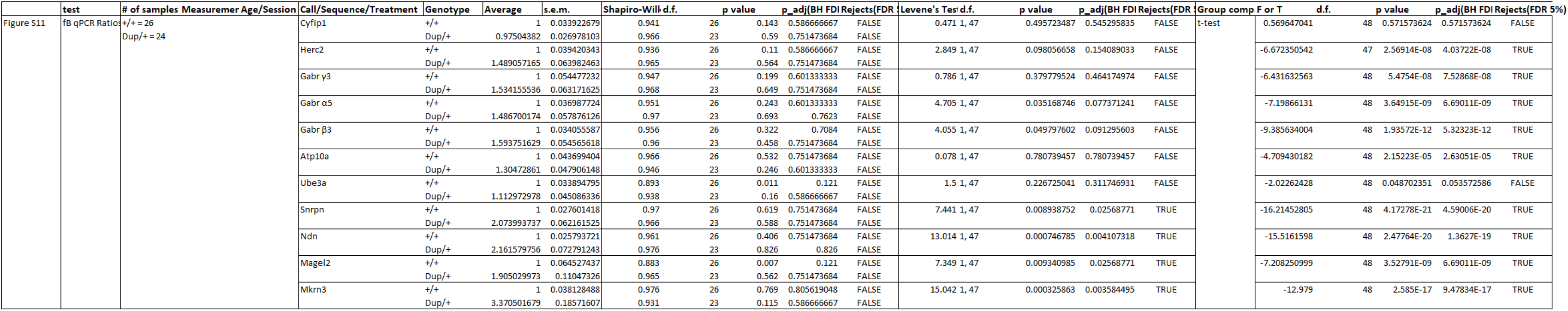
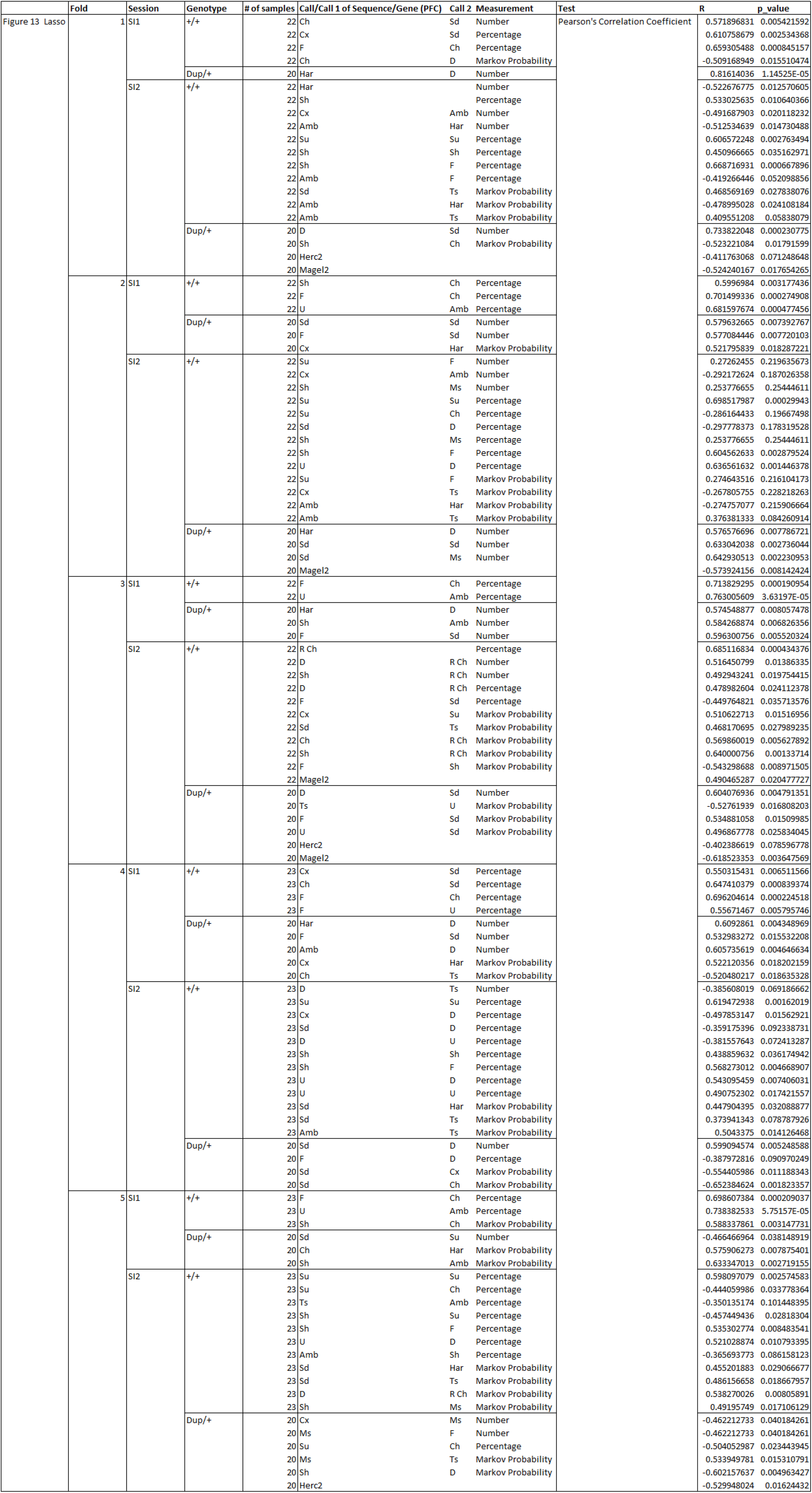
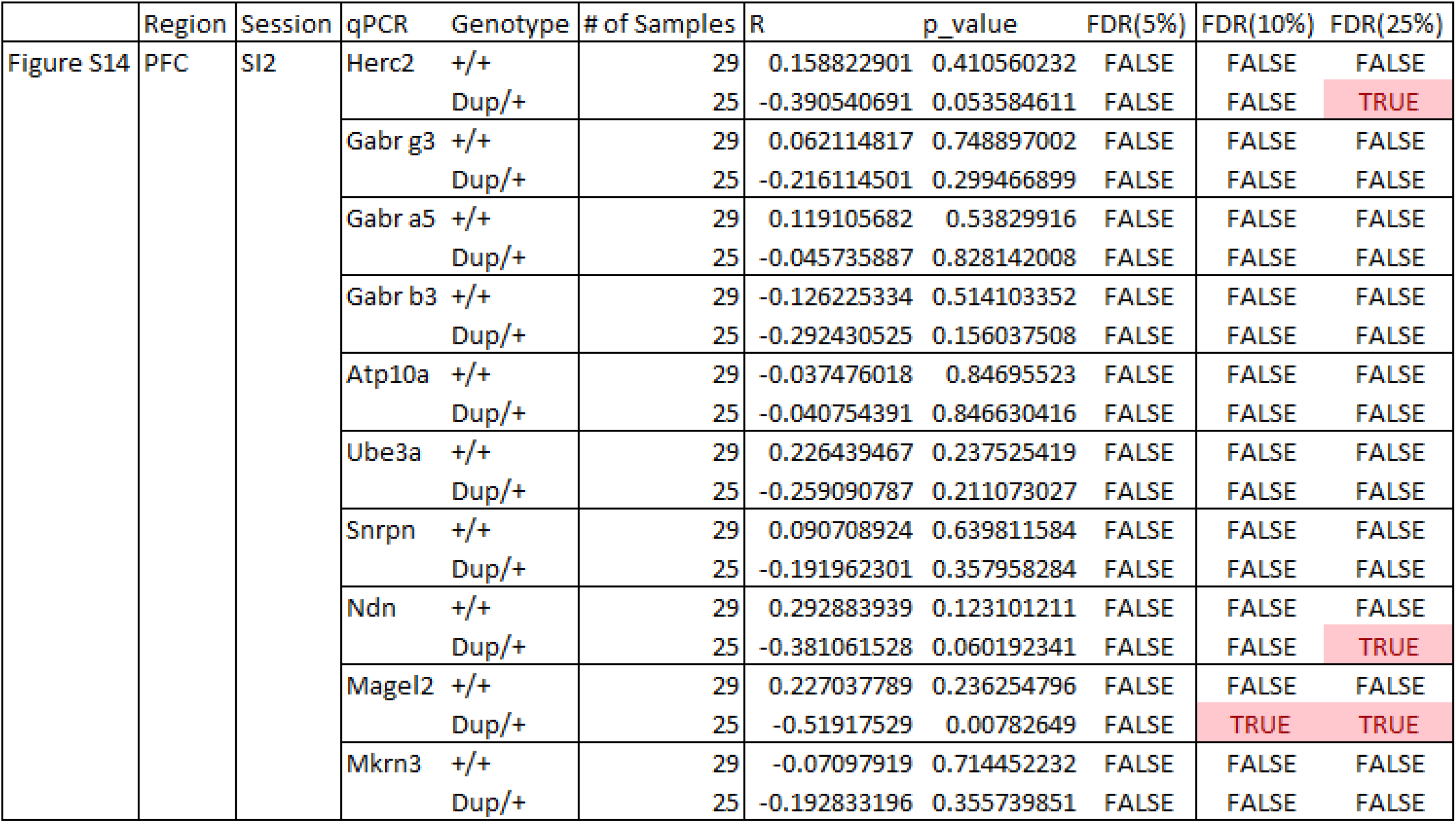

## References

1. Malhotra, D. and J. Sebat, CNVs: harbingers of a rare variant revolution in psychiatric genetics. Cell, 2012. 148(6): p. 1223–1241.

2. Zinkstok, J., et al., The 22q11.2 deletion syndrome from a neurobiological perspective. Lancet Psychiatry, 2019. 6(11): p. 951–960.

3. Hiroi, N. and T. Yamauchi, Modeling and Predicting Developmental Trajectories of Neuropsychiatric Dimensions Associated With Copy Number Variations. Int J Neuropsychopharmacol, 2019. 22(8): p. 488–500.

4. Hiroi, N., Critical Reappraisal of Mechanistic Links of Copy Number Variants to Dimensional Constructs of Neuropsychiatric Disorders in Mouse Models. Psychiatry and Clinical Neurosciences, 2018. 72(5): p. 301–321.

5. Hiroi, N., et al., Copy Number Variation at 22q11.2: from rare variants to common mechanisms of developmental neuropsychiatric disorders. Mol. Psychiatry, 2013. 18: p. 1153–1165.

6. Hiroi, N., et al., Mouse models of 22q11.2-associated autism spectrum disorder. Autism, 2012. S1(001): p. 1–9.

7. Satterstrom, F.K., et al., Large-Scale Exome Sequencing Study Implicates Both Developmental and Functional Changes in the Neurobiology of Autism. Cell, 2020. 180(3): p. 568–584 e23.

8. Fu, J.M., et al., Rare coding variation provides insight into the genetic architecture and phenotypic context of autism. Nat Genet, 2022. 54(9): p. 1320–1331.

9. Zhou, X., et al., Integrating de novo and inherited variants in 42,607 autism cases identifies mutations in new moderate-risk genes. Nat Genet, 2022. 54(9): p. 1305–1319.

10. Singh, T., et al., Rare coding variants in ten genes confer substantial risk for schizophrenia. Nature, 2022. 604(7906): p. 509–516.

11. Trost, B., et al., Genomic architecture of autism from comprehensive whole-genome sequence annotation. Cell, 2022. 185(23): p. 4409–4427 e18.

12. Collins, R.L., et al., A cross-disorder dosage sensitivity map of the human genome. Cell, 2022. 185(16): p. 3041–3055 e25.

13. Ozonoff, S., et al., The broader autism phenotype in infancy: when does it emerge? J. Am. Acad. Child Adolesc. Psychiatry, 2014. 53(4): p. 398–407.

14. Raza, S., et al., Relationship Between Early Social-Emotional Behavior and Autism Spectrum Disorder: A High-Risk Sibling Study. J Autism Dev Disord, 2019.

15. Gur, R.E., et al., Neurocognitive development in 22q11.2 deletion syndrome: comparison with youth having developmental delay and medical comorbidities. Mol. Psychiatry, 2014. 19(11): p. 1205–1211.

16. Esposito, G., N. Hiroi, and M.L. Scattoni, Cry, baby, cry: Expression of Distress as a Biomarker and Modulator in Autism Spectrum Disorder. Int J Neuropsychopharmacol, 2017. 20(6): p. 498–503.

17. Lawford, H.L.S., et al., Acoustic Cry Characteristics of Infants as a Marker of Neurological Dysfunction: A Systematic Review and Meta-Analysis. Pediatr Neurol, 2022. 129: p. 72–79.

18. LaGasse, L.L., A.R. Neal, and B.M. Lester, Assessment of infant cry: acoustic cry analysis and parental perception. Ment Retard Dev Disabil Res Rev, 2005. 11(1): p. 83–93.

19. Kikusui, T. and N. Hiroi, A Self-Generated Environmental Factor as a Potential Contributor to Atypical Early Social Communication in Autism. Neuropsychopharmacology, 2017. 42(1): p. 378.

20. Battaglia, A., The inv dup (15) or idic (15) syndrome (Tetrasomy 15q). Orphanet J Rare Dis, 2008. 3: p. 30.

21. Butler, M.G., Imprinting disorders in humans: a review. Curr Opin Pediatr, 2020. 32(6): p. 719–729.

22. Marini, C., et al., Clinical and genetic study of a family with a paternally inherited 15q11-q13 duplication. Am J Med Genet A, 2013. 161A(6): p. 1459–64.

23. Cook, E.H., Jr., et al., Autism or atypical autism in maternally but not paternally derived proximal 15q duplication. Am J Hum Genet, 1997. 60(4): p. 928–34.

24. Hoppman-Chaney, N.L., et al., Partial hexasomy for the Prader-Willi-Angelman syndrome critical region due to a maternally inherited large supernumerary marker chromosome. Am J Med Genet A, 2010. 152A(8): p. 2034–8.

25. Ingason, A., et al., Maternally derived microduplications at 15q11-q13: implication of imprinted genes in psychotic illness. Am. J. Psychiatry, 2011. 168(4): p. 408–417.

26. Mao, R., et al., Characteristics of two cases with dup(15)(q11.2-q12): one of maternal and one of paternal origin. Genet Med, 2000. 2(2): p. 131–5.

27. Al Ageeli, E., et al., Duplication of the 15q11-q13 region: clinical and genetic study of 30 new cases. Eur J Med Genet, 2014. 57(1): p. 5–14.

28. Roberts, S.E., et al., Characterisation of interstitial duplications and triplications of chromosome 15q11-q13. Hum Genet, 2002. 110(3): p. 227–34.

29. Schroer, R.J., et al., Autism and maternally derived aberrations of chromosome 15q. Am J Med Genet, 1998. 76(4): p. 327–36.

30. Bolton, P.F., et al., The phenotypic manifestations of interstitial duplications of proximal 15q with special reference to the autistic spectrum disorders. Am J Med Genet, 2001. 105(8): p. 675–85.

31. Browne, C.E., et al., Inherited interstitial duplications of proximal 15q: genotype-phenotype correlations. Am J Hum Genet, 1997. 61(6): p. 1342–52.

32. Depienne, C., et al., Screening for genomic rearrangements and methylation abnormalities of the 15q11-q13 region in autism spectrum disorders. Biol Psychiatry, 2009. 66(4): p. 349–59.

33. Bisba, M., et al., *Chromosome 15q11-q13 Duplication Syndrome: A Review of the Literature and 14 New Cases*. Genes (Basel), 2024. 15(10).

34. Mohandas, T.K., et al., Paternally derived de novo interstitial duplication of proximal 15q in a patient with developmental delay. Am J Med Genet, 1999. 82(4): p. 294–300.

35. Veltman, M.W., et al., A paternally inherited duplication in the Prader-Willi/Angelman syndrome critical region: a case and family study. J Autism Dev Disord, 2005. 35(1): p. 117–27.

36. Ungaro, P., et al., Molecular characterisation of four cases of intrachromosomal triplication of chromosome 15q11-q14. J Med Genet, 2001. 38(1): p. 26–34.

37. Bolton, P.F., et al., Chromosome 15q11-13 abnormalities and other medical conditions in individuals with autism spectrum disorders. Psychiatr Genet, 2004. 14(3): p. 131–7.

38. Urraca, N., et al., The interstitial duplication 15q11.2-q13 syndrome includes autism, mild facial anomalies and a characteristic EEG signature. Autism Res, 2013. 6(4): p. 268–79.

39. Isles, A.R., et al., Parental Origin of Interstitial Duplications at 15q11.2-q13.3 in Schizophrenia and Neurodevelopmental Disorders. PLoS Genet, 2016. 12(5): p. e1005993.

40. Parijs, I., et al., Population screening for 15q11-q13 duplications: corroboration of the difference in impact between maternally and paternally inherited alleles. Eur J Hum Genet, 2024. 32(1): p. 31–36.

41. Palmer, D.S., et al., Exome sequencing in bipolar disorder identifies AKAP11 as a risk gene shared with schizophrenia. Nat Genet, 2022. 54(5): p. 541–547.

42. Tamada, K. and T. Takumi, Neurodevelopmental impact of CNV models in ASD: Recent advances and future directions. Curr Opin Neurobiol, 2025. 92: p. 103001.

43. Gur, R.C. and R.E. Gur, Social cognition as an RDoC domain. Am J Med Genet B Neuropsychiatr Genet, 2016. 171B(1): p. 132–41.

44. Gur, R.C., et al., Neurocognitive performance in family-based and case-control studies of schizophrenia. Schizophr Res, 2015. 163(1-3): p. 17–23.

45. Corcoran, C.M., et al., Emotion recognition deficits as predictors of transition in individuals at clinical high risk for schizophrenia: a neurodevelopmental perspective. Psychol Med, 2015. 45(14): p. 2959–73.

46. Kohler, C.G., et al., Facial emotion perception differs in young persons at genetic and clinical high-risk for psychosis. Psychiatry Res, 2014. 216(2): p. 206–12.

47. Gur, R.C., et al., Neurocognitive growth charting in psychosis spectrum youths. JAMA Psychiatry, 2014. 71(4): p. 366–374.

48. Forsingdal, A., et al., Can Animal Models of Copy Number Variants That Predispose to Schizophrenia Elucidate Underlying Biology? Biol Psychiatry, 2019. 85(1): p. 13–24.

49. Hiroi, N., et al., A 200-kb region of human chromosome 22q11.2 confers antipsychotic-responsive behavioral abnormalities in mice. Proceedings of the National Academy of Sciences of the United States of America, 2005. 102(52): p. 19132–19137.

50. Suzuki, G., et al., Over-expression of a human chromosome 22q11.2 segment including TXNRD2, COMT and ARVCF developmentally affects incentive learning and working memory in mice. Human Molecular Genetics, 2009. 18(20): p. 3914–3925.

51. Gur, R.E., et al., A neurogenetic model for the study of schizophrenia spectrum disorders: the International 22q11.2 Deletion Syndrome Brain Behavior Consortium. Mol Psychiatry, 2017.

52. Stefansson, H., et al., CNVs conferring risk of autism or schizophrenia affect cognition in controls. Nature, 2014. 505(7483): p. 361–6.

53. Kendall, K.M., et al., Cognitive performance and functional outcomes of carriers of pathogenic copy number variants: analysis of the UK Biobank. Br J Psychiatry, 2019. 214(5): p. 297–304.

54. Kendall, K.M., et al., Cognitive Performance Among Carriers of Pathogenic Copy Number Variants: Analysis of 152,000 UK Biobank Subjects. Biol Psychiatry, 2017. 82(2): p. 103–110.

55. Chawner, S., et al., Genotype-phenotype associations in children with copy number variants associated with high neuropsychiatric risk in the UK (IMAGINE-ID): a case-control cohort study. Lancet Psychiatry, 2019. 6(6): p. 493–505.

56. Nakatani, J., et al., Abnormal behavior in a chromosome-engineered mouse model for human 15q11-13 duplication seen in autism. Cell, 2009. 137(7): p. 1235–1246.

57. Mirzaeian, L., et al., Investigating the influence of estrous cycle-dependent hormonal changes on neurogenesis in adult mice. Steroids, 2024. 212: p. 109513.

58. Hiramoto, T., et al., Transcriptional regulation of neonatal neural stem cells is a determinant of social behavior. BioRxiv, 2023. https://www.biorxiv.org/content/10.1101/2021.11.12.468452v2.

59. Nakamura, M., et al., Computational identification of variables in neonatal vocalizations predictive for postpubertal social behaviors in a mouse model of 16p11.2 deletion. Mol Psychiatry, 2021. 26(11): p. 6578–6588.

60. Yamauchi, T., G. Kang, and N. Hiroi, Heterozygosity of murine Crkl does not recapitulate behavioral dimensions of human 22q11.2 hemizygosity. Genes Brain and Behavior, 2020. 20(5): p. e12719.

61. Takahashi, T., et al., Structure and function of neonatal social communication in a genetic mouse model of autism. Mol Psychiatry, 2016. 21(9): p. 1208–14.

62. Harper, K.M., et al., Alterations of social interaction through genetic and environmental manipulation of the 22q11.2 gene Sept5 in the mouse brain. Human Molecular Genetics, 2012. 21(15): p. 3489–3499.

63. Hiramoto, T., et al., Tbx1: identification of a 22q11.2 gene as a risk factor for autism spectrum disorder in a mouse model. Hum Mol Genet, 2011. 20(24): p. 4775–85.

64. Suzuki, G., et al., Sept5 deficiency exerts pleiotropic influence on affective behaviors and cognitive functions in mice. Human Molecular Genetics, 2009. 18(9): p. 1652–1660.

65. Sakamoto, Y., et al., Prepartum bumetanide treatment reverses altered neonatal social communication but non-specifically reduces post-pubertal social behavior in a mouse model of fragile X syndrome. Genomic Psychiatry, 2025. 1(1): p. 61–72.

66. Ó Broin, P.B., M.V.; Takahashi, T.; Izumi, T.; Ye, K.; Kang, K.; Pouso, P.; Topolski, M.; Pena, J.L.; Hiroi, N., Computational Analysis of Neonatal Mouse Ultrasonic Vocalization. Current Protocols in Mouse Biology, 2018. 8((2)): p. e46.

67. Scattoni, M.L., et al., Unusual repertoire of vocalizations in the BTBR T+tf/J mouse model of autism. PLoS. One, 2008. 3(8): p. e3067.

68. Lai, J.K., et al., Temporal and spectral differences in the ultrasonic vocalizations of fragile X knock out mice during postnatal development. Behav. Brain Res, 2014. 259: p. 119–130.

69. Caruso, A., L. Ricceri, and M.L. Scattoni, Ultrasonic vocalizations as a fundamental tool for early and adult behavioral phenotyping of Autism Spectrum Disorder rodent models. Neurosci Biobehav Rev, 2020. 116: p. 31–43.

70. McInnes, L.H., J.; Melville, J., UMAP: Uniform manifold approximation and projection for dimension reduction., in *arXiv*. 2018. p. 03426.

71. Fonseca, A.H., et al., Analysis of ultrasonic vocalizations from mice using computer vision and machine learning. Elife, 2021. 10.

72. Kihara, T., et al., Reproduction of mouse-pup ultrasonic vocalzations by nanocrystalline silicon thermoacoustic emitter. Applied Physics Letters, 2006. 88(043902): p. 1–3.

73. Uematsu, A., et al., Maternal approaches to pup ultrasonic vocalizations produced by a nanocrystalline silicon thermo-acoustic emitter. Brain Res, 2007. 1163: p. 91–99.

74. Fukumitsu, K., et al., Amylin-Calcitonin receptor signaling in the medial preoptic area mediates affiliative social behaviors in female mice. Nat Commun, 2022. 13(1): p. 709.

75. Jabarin, R., S. Netser, and S. Wagner, Beyond the three-chamber test: toward a multimodal and objective assessment of social behavior in rodents. Mol Autism, 2022. 13(1): p. 41.

76. Hogart, A., et al., Chromosome 15q11-13 duplication syndrome brain reveals epigenetic alterations in gene expression not predicted from copy number. J Med Genet, 2009. 46(2): p. 86–93.

77. Yizhar, O., et al., Neocortical excitation/inhibition balance in information processing and social dysfunction. Nature, 2011. 477(7363): p. 171–8.

78. Sakamoto, T. and J. Yashima, Prefrontal cortex is necessary for long-term social recognition memory in mice. Behav Brain Res, 2022. 435: p. 114051.

79. Tanimizu, T., et al., Functional Connectivity of Multiple Brain Regions Required for the Consolidation of Social Recognition Memory. J Neurosci, 2017. 37(15): p. 4103–4116.

80. Wang, F., et al., Bidirectional control of social hierarchy by synaptic efficacy in medial prefrontal cortex. Science, 2011. 334(6056): p. 693–7.

81. Xu, P., et al., Pattern decorrelation in the mouse medial prefrontal cortex enables social preference and requires MeCP2. Nat Commun, 2022. 13(1): p. 3899.

82. DuBose, A.J., et al., Atp10a, a gene adjacent to the PWS/AS gene cluster, is not imprinted in mouse and is insensitive to the PWS-IC. Neurogenetics, 2010. 11(2): p. 145–51.

83. Kayashima, T., et al., Atp10a, the mouse ortholog of the human imprinted ATP10A gene, escapes genomic imprinting. Genomics, 2003. 81(6): p. 644–7.

84. Kayashima, T., et al., On the conflicting reports of imprinting status of mouse ATP10a in the adult brain: strain-background-dependent imprinting? J Hum Genet, 2003. 48(9): p. 492–493.

85. Varoquaux, G., Cross-validation failure: Small sample sizes lead to large error bars. Neuroimage, 2018. 180(Pt A): p. 68–77.

86. Kogan, J.H., P.W. Frankland, and A.J. Silva, Long-term memory underlying hippocampus-dependent social recognition in mice. Hippocampus, 2000. 10(1): p. 47–56.

87. Premoli, M., M. Memo, and S.A. Bonini, Ultrasonic vocalizations in mice: relevance for ethologic and neurodevelopmental disorders studies. Neural Regen Res, 2021. 16(6): p. 1158–1167.

88. Glass, T.J., et al., Ultrasonic vocalization phenotypes in the Ts65Dn and Dp(16)1Yey mouse models of Down syndrome. Physiol Behav, 2023. 271: p. 114323.

89. Tamada, K., et al., Genetic dissection identifies Necdin as a driver gene in a mouse model of paternal 15q duplications. Nat Commun, 2021. 12(1): p. 4056.

90. Schubert, T. and C.P. Schaaf, MAGEL2 (patho-)physiology and Schaaf-Yang syndrome. Dev Med Child Neurol, 2025. 67(1): p. 35–48.

91. Bosque Ortiz, G.M., G.M. Santana, and M.O. Dietrich, Deficiency of the paternally inherited gene Magel2 alters the development of separation-induced vocalization and maternal behavior in mice. Genes Brain Behav, 2022. 21(1): p. e12776.

92. Bertoni, A., et al., Oxytocin administration in neonates shapes hippocampal circuitry and restores social behavior in a mouse model of autism. Mol Psychiatry, 2021. 26(12): p. 7582–7595.

93. Meziane, H., et al., An Early Postnatal Oxytocin Treatment Prevents Social and Learning Deficits in Adult Mice Deficient for Magel2, a Gene Involved in Prader-Willi Syndrome and Autism. Biol. Psychiatry, 2015. 78(2): p. 85–94.

94. Fountain, M.D., et al., Magel2 knockout mice manifest altered social phenotypes and a deficit in preference for social novelty. Genes Brain Behav, 2017. 16(6): p. 592–600.

95. Higgs, M.J., et al., The parenting hub of the hypothalamus is a focus of imprinted gene action. PLoS Genet, 2023. 19(10): p. e1010961.

96. Smith, S.E., et al., Increased gene dosage of Ube3a results in autism traits and decreased glutamate synaptic transmission in mice. Sci Transl Med, 2011. 3(103): p. 103ra97.

97. Krishnan, V., et al., Autism gene Ube3a and seizures impair sociability by repressing VTA Cbln1. Nature, 2017. 543(7646): p. 507–512.

98. Punt, A.M., et al., Molecular and behavioral consequences of Ube3a gene overdosage in mice. JCI Insight, 2022. 7(18).

99. Silverman, J.L., et al., Behavioural phenotyping assays for mouse models of autism. Nat. Rev. Neurosci, 2010. 11(7): p. 490–502.

100. Saito, R., et al., Comprehensive analysis of a novel mouse model of the 22q11.2 deletion syndrome: a model with the most common 3.0-Mb deletion at the human 22q11.2 locus. Transl Psychiatry, 2020. 10(1): p. 35.

101. Bicks, L.K., et al., Prefrontal Cortex and Social Cognition in Mouse and Man. Front Psychol, 2015. 6: p. 1805.

102. Dennis, N.R., et al., Clinical findings in 33 subjects with large supernumerary marker(15) chromosomes and 3 subjects with triplication of 15q11-q13. Am J Med Genet A, 2006. 140(5): p. 434–41.

## References

1. Hiramoto T, Kang G, Suzuki G, Satoh Y, Kucherlapati R, Watanabe Y, et al. Tbx1: identification of a 22q11.2 gene as a risk factor for autism spectrum disorder in a mouse model. Hum Mol Genet. 2011;20(24):4775–85.

2. Takahashi T, Okabe S, Broin PO, Nishi A, Ye K, Beckert MV, et al. Structure and function of neonatal social communication in a genetic mouse model of autism. Mol Psychiatry. 2016;21(9):1208–14.

3. Nakamura M, Ye K, MB ES, Yamauchi T, Hoeppner DJ, Fayyazuddin A, et al. Computational identification of variables in neonatal vocalizations predictive for postpubertal social behaviors in a mouse model of 16p11.2 deletion. Mol Psychiatry. 2021;26(11):6578–88.

4. Semple BD, Blomgren K, Gimlin K, Ferriero DM, Noble-Haeusslein LJ. Brain development in rodents and humans: Identifying benchmarks of maturation and vulnerability to injury across species. Prog Neurobiol. 2013;106–107:1–16.

5. Fonseca AH, Santana GM, Bosque Ortiz GM, Bampi S, Dietrich MO. Analysis of ultrasonic vocalizations from mice using computer vision and machine learning. Elife. 2021;10.

6. Scattoni ML, Gandhy SU, Ricceri L, Crawley JN. Unusual repertoire of vocalizations in the BTBR T+tf/J mouse model of autism. PLoS One. 2008;3(8):e3067.

7. Hiramoto T, Boku S, Kang G, Abe S, Nagashima M, Barbachane Silva M, et al. Transcriptional regulation of neonatal neural stem cells is a determinant of social behavior. BioRxiv. 2023;https://www.biorxiv.org/content/10.1101/2021.11.12.468452v2.

8. Suzuki G, Harper KM, Hiramoto T, Funke B, Lee M, Kang G, et al. Over-expression of a human chromosome 22q11.2 segment including TXNRD2, COMT and ARVCF developmentally affects incentive learning and working memory in mice. Human Molecular Genetics. 2009;18(20):3914–25.

9. Suzuki G, Harper KM, Hiramoto T, Sawamura T, Lee M, Kang G, et al. Sept5 deficiency exerts pleiotropic influence on affective behaviors and cognitive functions in mice. Human Molecular Genetics. 2009;18(9):1652–60.

10. Harper KM, Hiramoto T, Tanigaki K, Kang G, Suzuki G, Trimble W, et al. Alterations of social interaction through genetic and environmental manipulation of the 22q11.2 gene Sept5 in the mouse brain. Human Molecular Genetics. 2012;21(15):3489–99.

11. Yamauchi T, Kang G, Hiroi N. Heterozygosity of murine Crkl does not recapitulate behavioral dimensions of human 22q11.2 hemizygosity. Genes Brain and Behavior. 2020;20(5):e12719.

12. Hiramoto T, Sumiyoshi A, Yamauchi T, Tanigaki K, Shi Q, Kang G, et al. Tbx1, a gene encoded in 22q11.2 copy number variant, is a link between alterations in fimbria myelination and cognitive speed in mice. Mol Psychiatry. 2022;27(2):929–38.

13. Sakamoto Y, Takano T, Shimoyama S, Hiramoto T, Hiroi N, Nakamura K. Prepartum bumetanide treatment reverses altered neonatal social communication but non-specifically reduces post-pubertal social behavior in a mouse model of fragile X syndrome. Genomic Psychiatry. 2025;1(1):61–72.

14. Hammerschmidt K, Radyushkin K, Ehrenreich H, Fischer J. Female mice respond to male ultrasonic ’songs’ with approach behaviour. Biol Lett. 2009;5(5):589–92.

